# Integrated Multi-omic Profiling Reveals Early Regulatory Events in Dexamethasone Muscle Atrophy

**DOI:** 10.1101/2025.06.12.659253

**Authors:** Suzuka Nakagawa, Aristotelis Misios, Oliver Popp, Philipp Mertins, Jens Fielitz, Ernst Jarosch, Thomas Sommer

**Affiliations:** Max Delbrück Center for Molecular Medicine in the Helmholtz Association, Berlin, Germany Institute for Biology.; Humboldt-University zu Berlin, Berlin, Germany; Berlin Institute of Health, Berlin, Germany; DZHK (German Center for Cardiovascular Research), partner site Berlin, Germany, DZHK (German Center for Cardiovascular Research), partner site Greifswald, Germany, Department of Internal Medicine B, Cardiology, University Medicine Greifswald, Greifswald, Germany; Institute for Biomedical Translation Lower Saxony, Hannover, Germany

## Abstract

Skeletal muscle atrophy and weakness are major contributors to morbidity, prolonged recovery, and long-term disability across a wide range of diseases. Atrophy is caused by breakdown of sarcomeric proteins resulting in loss of muscle mass and strength. Molecular mechanism underlying the onset of muscle atrophy and its progression have been analysed in patients, mice, and cell culture but the complementarity of these model systems remains to be explored. Here, we applied deep-coverage transcriptomic and proteomic profiling to characterize dynamic changes during dexamethasone-induced atrophy in the widely used murine skeletal muscle cell line C2C12. Comparison with published datasets confirmed that muscle differentiation is well recapitulated in C2C12 myotubes. Under dexamethasone treatment, this model was particularly suited to capture early atrophy events. We identified alterations in mitochondrial gene expression and differential alternative splicing events during early-stage myotube atrophy. This dataset complements existing *in vivo* data and provides novel insights into the regulatory processes during skeletal muscle wasting.

## Introduction

Skeletal muscle atrophy is a condition which severely impacts patients’ quality of life by reducing muscle mass and strength. Research into mechanisms of atrophy onset and progression provides identification of critical factors whose activity may be modified to delay or reverse muscle wasting and ultimately promote recovery of patients. Atrophy is tightly associated with proteolytic systems like the ubiquitin proteasome system (UPS) and the autophagy lysosome pathway (ALP) that are known to mediate degradation of sarcomeric proteins (Wing and Goldberg, 1993, Schiaffino et al. 2013). Although several proteins termed atrogenes have been identified as specifically associated with striated muscle atrophy (e.g. *Fbxo32*/Atrogin-1, *Trim63*/MuRF1, *Fbxo30*/MUSA1, *Fbxo21*/SMART, *Trim32*) (Bodine et al. 2001, Sartori et al. 2013, Milan et al. 2015) a robust mechanism on how they contribute to disassembly of sarcomeres or degradation of its constituents is still under investigation. Moreover, transcriptional and translational processes that initiate, accelerate or prevent degradation of sarcomere components are still obscure.

Conventional research on skeletal muscle atrophy mostly relies on results obtained from mouse models and are complemented by various muscle cell lines. Immortalized murine C2C12 myoblasts are capable of differentiating into multinuclear bodies (myotubes) with formation of sarcomeres. These cells can then be used to further induce myotube atrophy by using various stimuli, such as the synthetic glucocorticoid dexamethasone (dex). Glucocorticoids (GC) bind to glucocorticoid receptors (GR) to allow nuclear translocation of the GR complex. Here it upregulates expression of FoxO transcription factors to transcribe atrogenes, such as MAFbx/Atrogin-1 and MuRF1 (Bodine et al. 2001, Sandri et al. 2004, Waddell et al. 2008). In clinical settings, prolonged or high dosage administration of GC e.g. to counter inflammation by inhibiting the NF-κB pathway, can trigger muscle atrophy and induce degradation of sarcomeric proteins (Auphan et al. 1995, Shimizu et al. 2011, Lecker et al. 1999).

Skeletal muscle differentiation, senescence, regeneration and disease states are best studied in tissues where myofibrils containing myotube bundles are exposed to a multitude of endocrine and metabolic signals from surrounding skeletal, neural and vascular structures. Several mRNA and protein datasets have been published for murine or rat muscle tissue describing molecular changes in atrophy. Most of these studies focus on one particular atrophy model with a single screening technique (Lang et al. 2017, Lang et al. 2018, Shavlakadze et al. 2019) while others cross-compare multiple atrophy models, such as sarcopenia, cancer cachexia and dex-induced atrophy with protein and RNA data (Hunt et al. 2021, Hitachi et al. 2021). Abdelmoez et al. (2020) published a cross-comparative analysis of published transcriptomic datasets from mouse (C2C12), rat (L6) and human skeletal muscle cell lines, albeit without atrophy stimuli. The heterogeneity in experimental set-ups for muscle studies are known to give a high variability in results which are difficult to compare (Hughes et al. 2023). Thus, a comprehensive dataset obtained from a commonly-used cell line, which has lower biological heterogeneity compared to animals, is cost-efficient, and is permissive to extensive protocol optimization would provide valuable information for atrophy studies in animals. Additionally, comparison to published mouse dataset of the same atrophy stimulus would highlight the biological relevance and limitation of these models.

Here, we present a matched time course dataset of deep mRNA and whole proteome coverage of C2C12 myotubes treated with dex. Our results give information on cellular events during myoblast to myotube differentiation with strong reproducibility of observations previously seen in mice. Furthermore, our data appear to portray relatively early events following dex exposure, with insights into cellular activities preceding sarcomeric breakdown. We expect this resource to provide new perspectives of atrophy, to better design and interpret future skeletal muscle studies by highlighting characteristics specific to C2C12 cells.

## Results

### Validation of the C2C12 Dexamethasone Atrophy Model

The aim of this study was to quantitate changes in RNA abundance and protein amounts across defined stages of C2C12 myoblast-to-myotube differentiation and subsequent dex-induced atrophy. Dex was chosen as the atrophy stimulus, as several disease states resulting in muscle atrophy (e.g. sepsis, acidosis) have higher levels of circulating endogenous GC and patients treated with dex in clinical settings display muscle atrophy, thus providing insight useful to multiple atrophy contexts (Braun et al. 2011, Lecker et al. 1999). Furthermore, dex can also be used in rodents to induce atrophy, allowing cross-comparison of cell line to animals. To this end, myoblasts were differentiated into myotubes for 7 days followed by treatment with 100 µM dex for 24 or 72 hrs to induce atrophy (Fig. 1A). A 7 day differentiation period was chosen based on literature as the longest commonly used differentiation for C2C12 cells (Cooper et al. 2004, Ferri et al. 2009). The rational was to a) maximise the number of fully differentiated myotubes, b) allow sufficient time for atrophy to occur while maintaining cell adherence, and c) reduce the presence of undifferentiated myoblasts. Dex concentration was guided by published studies, followed by titration experiments. Final conditions were selected based on microscopic observations that revealed the most noticeable diameter loss after 24 and 72 hrs (data not shown).

**Figure 1:**
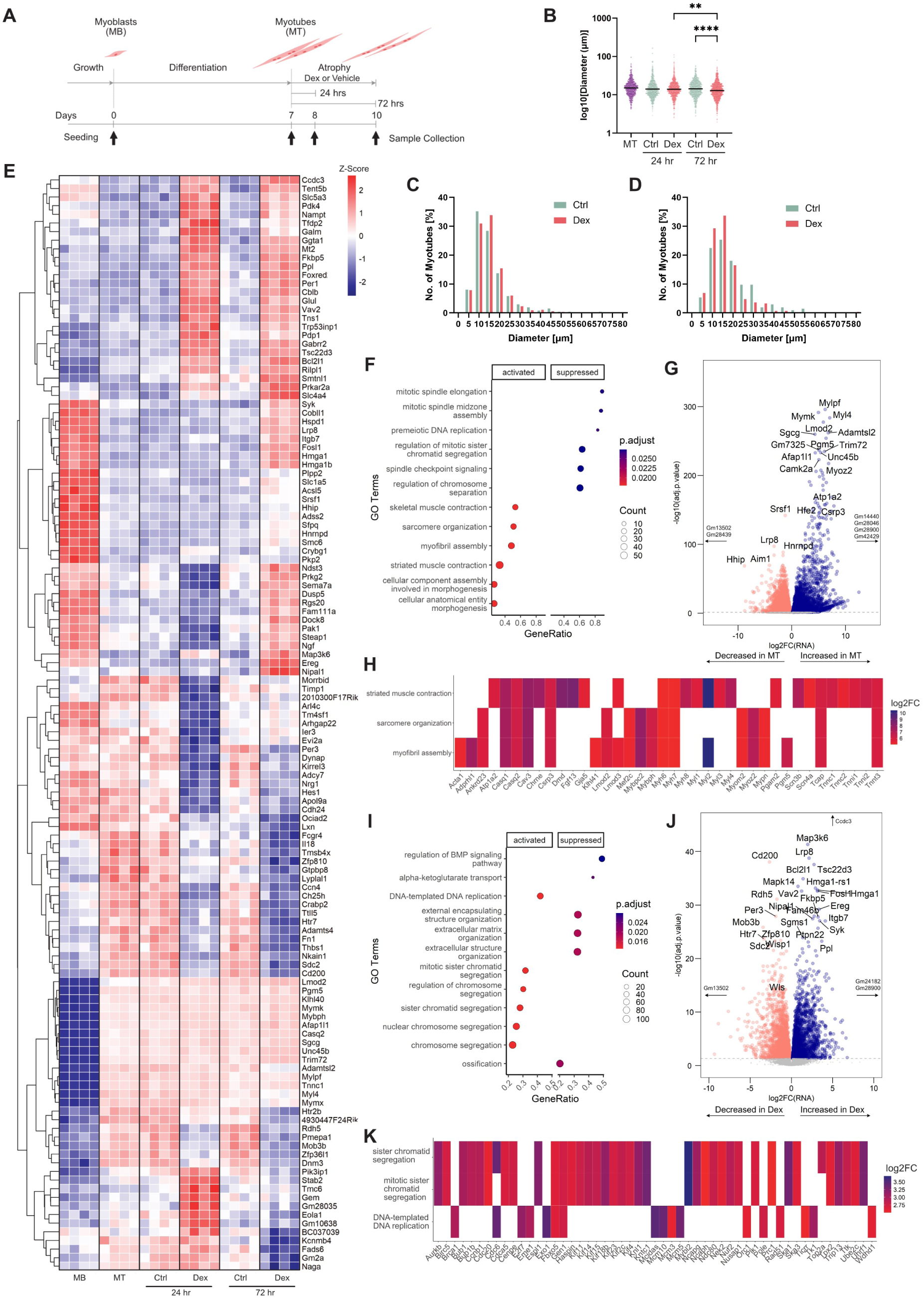
C2C12 cells show differentiation and atrophy phenotypes. Bulk RNA-seq reveals expression of differentiation and atrophy markers. A) Timeline schematic of sample collection for RNA-seq and TMT-MS data collection. C2C12 myoblasts (MB) were differentiated for 7 days into myotubes (MT), before switching to dexamethasone-containing (100µM, dex) atrophy medium or solvent control (0.1% ethanol, vehicle). Arrows indicate days of sample collection. B) Diameter measurements of fast skeletal MyHC positive C2C12s at indicated time points. Each data point indicates a diameter measurement. Five images were analysed per biological replicate, n=3. Lines indicate geometric mean. Significance was assessed by a Kruskal-Wallis test. **≤0.01, ****≤0.0001 C) and D) No. of MyHC positive C2C12s binned for indicated diameters at 24 hr (C) and 72hr (D). Values are shown as % of total. Five images were analysed per biological replicate, n=3. E) Z-score Heat map of normalized transcript counts of top 15 significantly changing genes (adj.p-value <=0.05) from comparisons of MB <> MT, Dex <> Ctrl 24 hr, Dex <> Ctrl 72 hr, Dex 24 hr <> Dex 72 hr. Euclidean clustering is shown. F) GSEA analysis of MT enriched biological pathways compared to MB using clusterProfiler. G) Volcano plot of log2FoldChange in transcript plotted against -log10(adj.p-value) for MT (right, blue) against MB (left, pink). Grey dashed line indicates adj.p-value=0.05. H) Plot of genes belonging to the top 3 activated biological processes from F). Colours correspond to the log2FoldChanges (log2FC) in normalized counts of transcript. Only genes with log2FC≥6 are shown. I) GSEA analysis of Dex over Ctrl at the 72 hr time point. J) Volcano plot of log2FoldChange in transcript plotted against -log10(adj.p-value) for Dex (right, blue) against ctrl samples at 72 hr (left, pink). Grey dashed line indicates adj.p-value =0.05. K) Plot of genes belonging to the top 3 activated biological processes from I). Colours correspond to the log2FoldChanges in normalized counts of transcript. Only genes with log2FC≥2.5 are shown.

We validated our C2C12 differentiation and atrophy model by assessing fusion indices across six conditions: undifferentiated myoblasts, day 7 myotubes, vehicle-treated controls at 24 and 72 hrs, and dex-treated samples after 24 and 72 hrs. Cells were fixed and stained with an anti-MyHC antibody, and fusion indices were calculated as the number of Hoechst-stained nuclei within MyHC-positive cells relative to the total nuclei per field of view. A significant increase of multinuclear cells was observed between myoblasts and all differentiated conditions (p-value <0.0001), with no significant differences among later time points (Supp. Fig. 1A). These results confirm that the protocol reliably produces differentiated myotubes and that subsequent dex-treatment does not significantly alter the fusibility of these cells.

Dex-treatment of myotubes for 24 hr led to a slight, statistically insignificant reduction in MyHC-positive cell diameter compared to vehicle-treated controls (Fig. 1B, 1C). After 72 hr, diameter reduction became more pronounced, as indicated by a shift in geometric mean values (Fig. 1B, 1D), consistent with the onset of atrophy. Further decrease between 24- and 72-hr dex-treated samples supported a progressive atrophy phenotype (Supp. Fig. 1B). In summary, this experimental setup reliably models C2C12 myoblast differentiation followed by dex-induced atrophy, and was used to generate samples for next-generation bulk mRNA sequencing (RNA-seq) to analyse changes in mRNA expression, splicing and mass spectrometry to quantitate differences in protein abundance.

### Transcriptomic analyses reveal tissue-comparable differentiation hallmarks and identifies novel putative contributors of atrophy

From our RNA-seq efforts, we detected ∼30,000 genes across all time points. Principal component analysis (PCA) revealed pronounced clustering of biological replicates for each condition, confirming the reproducibility of transcriptional states (Supp. Fig. 1C). As an initial validation step, we quantitated the expression of known myogenic factors (*Myf5, Myog*) (Rudnicki et al. 1993, Wright et al. 1989) and atrogenes (*Trim63, Fbxo32, Fbxo30*) (Bodine et al. 2001, Sartori et al. 2013) using RT-qPCR (Supp. Fig. 1D). The expression patterns of these genes were consistent with published data (Ferri et al. 2009, Abdelmoez et al. 2020) and in agreement with our RNA-seq results (Supp. Fig. 1E). Notably, *Fbxo30* showed differential expression only in RT-qPCR but not in RNA-seq, likely reflecting differences in detection sensitivity. In general, this indicated that the quantification of our RNA-seq data gave reliable information on actual transcriptional changes.

To characterise gene expression dynamics during differentiation and atrophy, RNA profiles from all post-differentiated time points were compared pairwise to the myoblast condition (adj. p-value≤0.05) and visualized using normalised counts in a heat map (Fig. 1E). In myotubes, we observed significant upregulation of transcripts encoding myoblast fusion regulators (Myomaker/*Mymk*, Myomixer/*Mymx*; Millay et al. 2013, Bi et al. 2017), key myogenic regulators (Klhl40/*Klhl40),* and striated muscle specific chaperones involved in sarcomere organisation (Unc45b/*Unc45b;* Price et al. 2002). Additionally, we detected increased expression of the Ca^2+^-binding protein Troponin C (*Tnnc1*), the ubiquitin E3 ligase *Trim72* that negatively regulates differentiation (Lee et al. 2010), and Camk2a involved in Ca^2+^ signalling (Chin 2005) (Fig. 1E, 1G). Gene set enrichment analysis (GSEA) further revealed upregulation of pathways related to skeletal muscle contraction, sarcomere organization, and myofibril assembly (GO: 0003009, GO:0045214, GO:0030239; Fig. 1F), including enrichment of genes encoding several thick and thin filaments, and Z-disc proteins (Fig. 1H). Conversely, transcripts related to regulation of mitosis (e.g. mitotic spindle elongation; GO:0000022, spindle checkpoint signalling; GO:0031577, regulation of chromosome separation: GO: 1905818) were downregulated in myotubes. This included *Zwint, Zwilch, Birc5, Cenpe,* and *Cdc20* (Supp. Table 1).

In the 24 hr dex-treated sample, pairwise comparison with vehicle-treated controls revealed significant induction of GC-responsive genes such as *Tsc22d3,* a GC induced leucine zipper, and *Fkbp5,* a modulator of GR activity (Supp. Fig. 1F) (D’Adamio et al. 1997, Wochnik et al. 2005). Interestingly, we also observed increased levels of *Cblb*, which encodes an ubiquitin E3 ligase that targets Insulin Receptor Substrate (IRS-1) for degradation promoting FOXO3-mediated atrogene expression (Nakao et al. 2009). Moreover, *Dcun1d3*, a NEDD8 E3 ligase and one of five mammalian homologs of yeast *DcnI* (Keuss et al. 2016), was also increased. Because NEDD8 is a regulator of Cullin-type ubiquitin ligase activity and given the growing list of these complexes reported to be associated with skeletal muscle atrophy, the upregulation of this protein suggests a potential novel regulatory role in skeletal muscle atrophy. Despite the differential expression of ∼4,800 genes at this time point, GSEA analysis did not reveal significant enrichment of specific biological pathways.

In contrast, the 72 hr dex-treated samples showed a distinct transcriptomic response. We observed reduced expression of *Wisp1* and *Wls* (Fig. 1I), suggesting downregulation of Wnt signalling pathway (Cadigan 2008, Jung et al. 2018). We also observed a continued increase of *Tsc22d3* and *Fkbp5* expression, along with several others already observed at 24 hr (*Vav2, Ppl*) (Fig. 1J). In fact, ∼540 genes were significantly upregulated and ∼300 genes downregulated at both 24 and 72 hrs of dex-treatment compared to their respective vehicle treated controls (Supp. Fig. 1G, adj.p-value<0.05, -1<log2FoldChange<0.5). GSEA analysis for the 72 hr time point revealed enrichment of pathways related to DNA replication and mitotic progression. This was due to increased transcription of several kinesin family members (*Kif2c, Kif4, Kif11, Kif14, Kif15*), and genes previously downregulated during myotube fusion (*Birc5, Cdc20, Cenpe*) (Fig. 1K).

### Alternative Splicing Dynamics During Atrophy

To investigate whether myogenic differentiation and dex-induced myotube atrophy affect alternative splicing, we analysed differential alternative splicing (DAS) events using a threshold of transcripts per million (TPM)>1. Across conditions, we detected 38,741 isoforms, with most genes expressing more than 2 isoforms (Fig. 2A). Although GSEA did not yield enriched pathways at 24 hr, DAS analysis revealed biologically meaningful isoform switching in 416 genes, many involving changes in protein domains or intracellular localization (Fig. 2A, 2B). In contrast, only 247 genes underwent isoform switching during differentiation from myoblasts to myotubes (Fig. 2B). Gene enrichment analysis of these 416 genes revealed a strong association with translational and post-translational protein modifications (Fig. 2C, adj.p-value≤0.05), highlighting an additional regulatory layer active during early stages of myotube atrophy.

**Figure 2:**
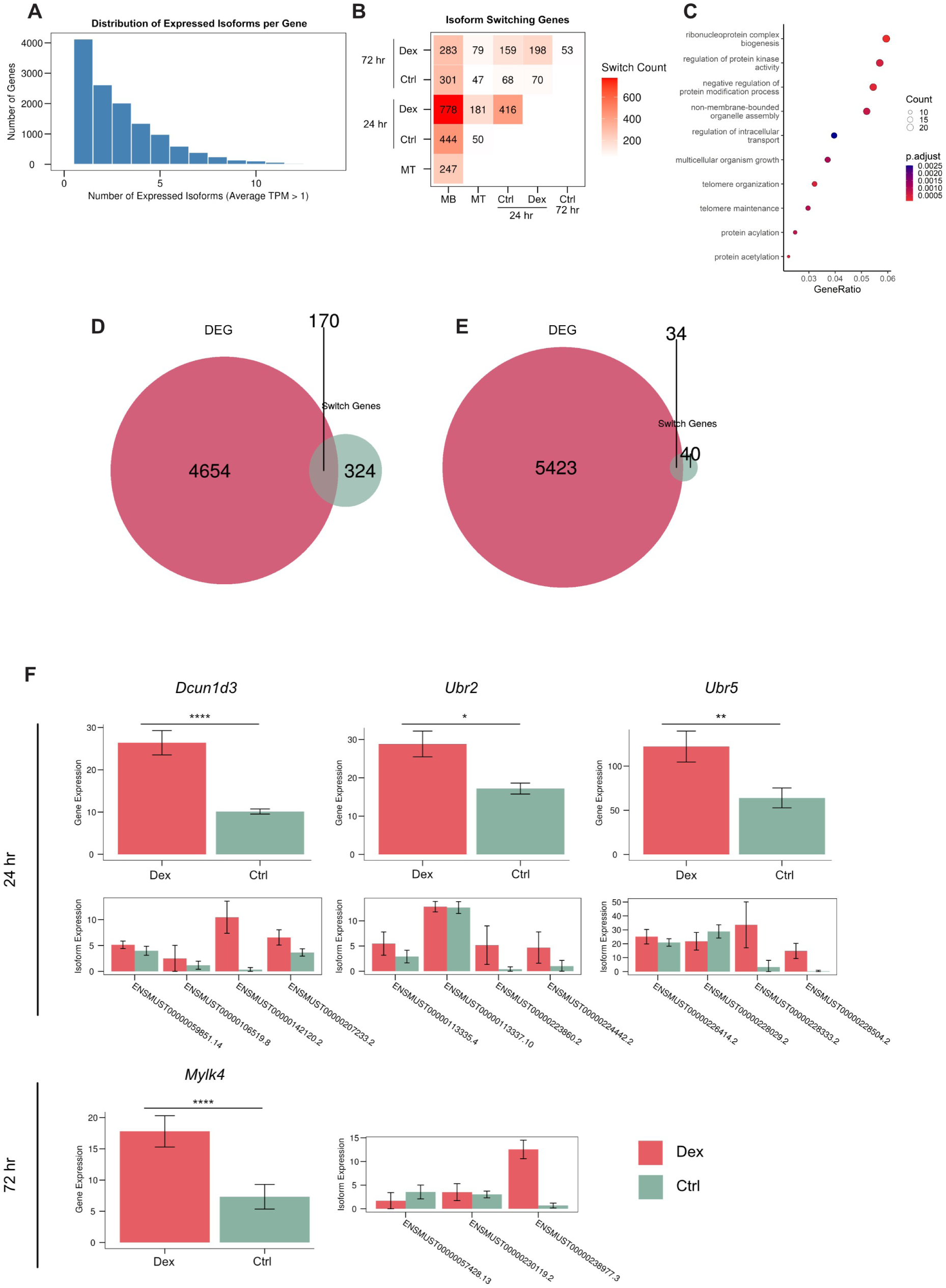
Differential alternative splicing (DAS) events in differentiation and dexamethasone atrophy in C2C12 cells. A) Histogram of number of isoforms per gene identified in the RNA-seq experiment. TPM>1. Counts greater than 15 isoforms are not included for legibility. B) Number of DAS events identified per comparison (adj.p<0.05). Isoform switches were filtered for consequences (see IsoformSwitchAnalyzeR documentation) C) GSEA of significantly DAS genes between Dex and Ctrl 24 hr. D) and E) Venn diagram of genes undergoing DAS and DEG for 24 hr (D) and 72hr (E). This clusterProfiler analysis did not filter for biological consequences, since DEG does not take this into account. F) Gene expression and isoform expression of *Dcun1d3, Ubr2, Ubr5* at 24 hr or *Mylk4* at 72 hr post Dex or vehicle Ctrl treatment.

Next, we examined whether genes undergoing DAS were also identified as differentially expressed genes (DEG) to assess the extent to which isoform abundance influence total gene expression. Our results indicated that approximately 3.5 % of DEGs also exhibited DAS within the first 24 hr of dex-induced atrophy. Interestingly, this overlap dropped to less than 1 % at 72 hrs, despite a rise in the number of DEGs overall. These findings suggest that isoform-specific transcriptional regulation is largely confined to the early phase of atrophy (Fig. 2D, 2E). This temporal specificity was further supported by a reduction in the total number of DAS events at 72 hrs, where only 53 genes exhibited isoform switches between vehicle and dex-treated samples, and 198 genes showed changes between 24 hrs and 72 hrs dex; a number considerably less than that observed at 24 hrs (Fig. 2B).

All genes that underwent both DEG and DAS are provided in supplementary table 2. Noteworthy among them at the 24 hr time point were *Dcun1d3, Ubr2, Ubr5* and at the 72 hr time point, *Mylk4* (Fig. 2F). *Ubr2 and Ubr5*, both encoding ubiquitin E3 ligases, have been previously associated with muscle atrophy and hypertrophy, respectively (Gao et al. 2022, Hughes et al. 2021). Both genes were significantly upregulated following 24 hr dex-treatment (adj.p=0.031 for *Ubr2*, adj.p=0.00031 for *Ubr5*). Interestingly, *Ubr5* displayed isoform changes only in non-coding transcripts, while coding transcripts remained unchanged. In contrast, *Ubr2* exhibited alternative splicing of a second coding isoform (ENSMUST00000113335.4), resulting in a protein variant differing by 13 amino acids compared to the reference sequence (Uniprot ID Q6WKZ8-1 vs. -2). Despite transcriptional upregulation, UBR2 protein levels decreased, whereas UBR5 protein abundance slightly increased under atrophy (Supp. Fig. 1H). At 72 hrs, *Mylk4*, a kinase acting on myosin light chains, whose expression in skeletal muscle increases with dihydrotestosterone treatment and enables Ca^2+^ induced contraction with a cardiac and a skeletal muscle variant (Sakakibara et al. 2021), showed a shift in isoform expression. The skeletal muscle-specific isoform (ENSMUST00000238977.3) increased, while the cardiac variant (ENSMUST00000057428.13) was downregulated, suggesting that C2C12 myotubes differentiated with FBS may predominantly express the cardiac *Mylk4* isoform. A third isoform (ENSMUST00000230119.2), also classified as protein-coding, was expressed at a higher level in controls.

### Mitochondrial Translation and Oxidative Phosphorylation Are Upregulated in Dexamethasone-Induced Atrophy

To study the proteomic landscape, we performed Tandem-Mass Tag Mass Spectrometry (TMT-MS) on proteins from C2C12 cells that were harvested at the same time points used for the RNA-seq experiment. TMT-MS allows multiplexing of a large number of samples and is highly sensitive in detecting low abundance proteins. This analysis enabled quantification of 9,613 proteins including known atrophy markers such as MuRF1/*Trim63*, MAFBx/Atrogin-1/*Fbxo32*, MUSA1/*Fbxo30* (Bodine et al. 2001, Sartori et al. 2013). This high coverage exceeds the typical 3,500-5,000 proteins quantified in mouse tissue, likely due to homogeneity of cells compared to animal tissue preparations, presence of residual undifferentiated mononuclear cells (Lang et al.2018, Hunt et al. 2021) or methodological differences. PCA showed good clustering of biological replicates for the individual samples, reflecting different stages of differentiation and atrophy (Supp. Fig. 2A).

During differentiation, we observed increased expression of sarcomeric proteins (Fig. 3A), with MYH1 being the most abundant, followed by MYH7 and MYH4, suggesting a fiber type composition biased towards type IIx, type I and type IIb fibers, and low levels of type IIa (MYH2). Isoforms of troponin (C, I and T) from both skeletal and cardiac muscle were also detected. MYH6, the second cardiac MyHC isoform, was absent or below detection limit (Fig. 3A). Similarly, myosin light chain isoforms MYL3 and MYL7, representing ventricular and atrial cardiomyocyte forms, respectively (Schiaffino and Reggiani 2011), were not detected. In comparison to the highly abundant sarcomere components, which cluster in the higher intensity ranks, ubiquitin E3 ligases (including atrogenes) spanned a wide range of abundance while myogenic regulators (MyoD1, Myogenin, PAX7) ranked lower, indicating minimal contamination from undifferentiated cells in myotube samples (Fig. 3A).

**Figure 3:**
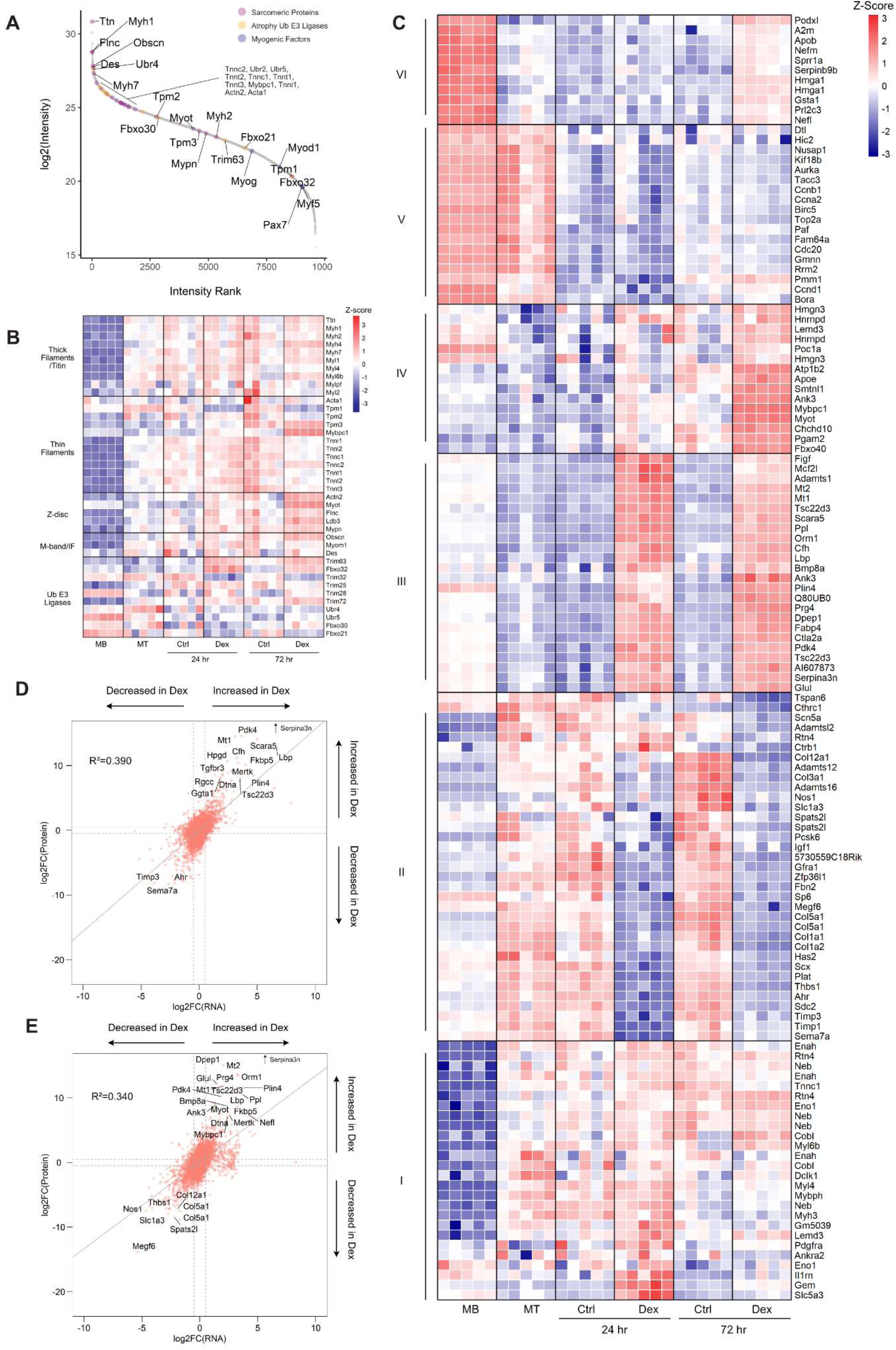
TMT-MS proteomic analyses of differentiation and dex-induced atrophy in C2C12 cells. A) Intensity ranking of sarcomeric proteins, ubiquitin (Ub) E3 ligases and myogenic factors. B) Z-score heat map of sarcomeric components and Ub E3 ligases associated with skeletal muscle atrophy. Genes were not filtered for significance. C) Z-score heat map of median-MAD normalized intensity values of top 15 genes with significant adj.p-value ≤0.05 from TMT-MS data. Up/downregulation in comparisons of: MT<>Dex 24 hr, MT <> Dex 72 hr, Dex 24 hr<> Dex 72 hr, Dex <> Ctrl 24 hr, Dex <> Ctrl 72 hr were pooled together and filtered for unique entries. Euclidean clustering is shown. D) and E) Log2FoldChanges of RNA (x-axis) plotted against those of proteins identified by TMT-MS (y-axis) for Dex <> Ctrl 24 hr (D) and Dex <> Ctrl 72 hr (E). R^2 values were calculated based on linear regression modelling. Serpina3n (*Serpina3n)* has been omitted in both plots, due to the log2FC being significantly further from the rest of the data points.

Contrary to our expectations, no general loss of sarcomeric proteins was observed with dex-treatment, with the exception of Ca^2+^ sensing proteins, including Tropomyosin (TPM1, TPM2) and Troponins (I, C and T) (Fig. 3B). As anticipated, MuRF1 (*Trim63*) and Atrogin-1 (*Fbxo32*) increased along with Serpina3n, a recently proposed atrophy biomarker (Fig. 3B, 3C) (Gueugneau et al. 2018, Hunt et al. 2021). However, other ubiquitin E3 ligases such as TRIM72, TRIM25, UBR4, UBR5, and FBXO30 showed minimal changes. Interestingly, TRIM32 and FBXO21/SMART were downregulated despite being linked to degradation of Desmin and Z-discs (Cohen et al. 2012, Milan et al. 2015) (Fig. 3B). Overall, this suggests that the C2C12 dex atrophy model recapitulates regulatory events from mouse tissue experiments but is less suited for studying sarcomere breakdown by TRIM32 or FBXO21/SMART. Additional UPS components with significant expression changes (adj.p-value<0.05) are detailed in Supp. Fig. 2B.

Notably, the top 15 most regulated proteins during differentiation and atrophy did not include sarcomeric proteins or atrogenes (Fig. 3C). Both atrophy time points shared 485 upregulated proteins (cluster IV), enriched in processes related to mitochondrial translation and mitochondrial gene expression (GO: 0032543, GO: 01450053) (Fig. 4A, 4B). These included mitochondrial ribosomal proteins (MRPL1, MRPL3, MRPS22, MRPS25) and respiratory chain components (COX5A, COX6C, NDUFA9) (Fig. 4A, Supp. Fig. 3A). Interestingly, these changes were not paralleled at the mRNA level, suggesting translation from mRNA pools generated during differentiation (Supp. Fig. 3A). Notably, ERRα and NRF-1 two transcription factors promoting mitochondrial gene expression, were significantly increased (adj.p-value <0.001, Supp. Fig. 3B).

**Figure 4:**
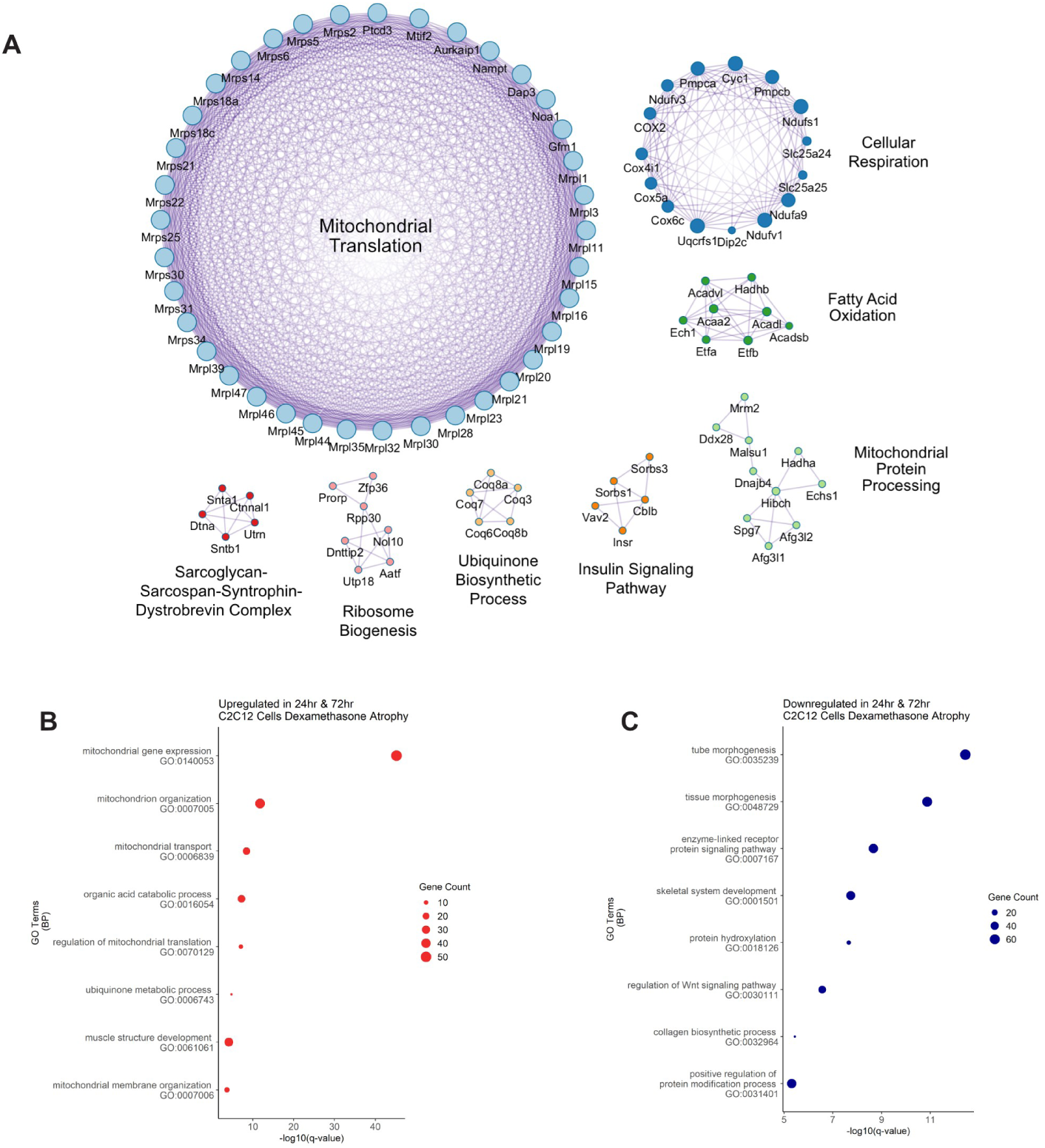
Protein-protein interaction enrichment analysis reveals significant enrichment of mitochondrial processes in dex-induced atrophy of C2C12 cells. A) Protein-protein interaction enrichment analysis (Metascape) of upregulated proteins in both Dex 24 hr and 72 hr with adj.p-value ≤0.05. Figure created with Cytoscape. B) and C) GO enrichment analysis of up/down regulated proteins common to both 24 hr and 72 hr dex atrophy in C2C12 cells with adj.p-value ≤0.05.

In contrast, cytoplasmic ribosomal subunits were substantially lower at both atrophy time points, likely contributing to the general decrease in protein translation observed in atrophic muscle (Supp. Fig. 3C, 3D) (Savary et al. 1998, Gordon et al. 2013). Additional downregulated proteins included those involved in skeletal muscle development and Wnt signaling (Fig. 4C), implicating an impairment of cell cycle progression during dex exposure.

### mRNA and Protein Correlation Reveals Post-Transcriptional Control

We next compared changes in RNA-seq and proteomics data. The 24 hr atrophy time point showed the strongest correlation between mRNA and protein changes (R^2^ = 0.390), followed by 72hrs (R^2^ value = 0.340) (Fig. 3D, 3E). However, the wide spread of mRNA values (x-axis) relative to protein fold-changes (y-axis) at 72 hrs suggests impaired translation or post-translational regulation. Surprisingly, differentiation (myoblast vs. myotube) had the lowest RNA-protein correlation (R^2^ value=0.269, Supp. Fig. 2C), further supporting a delay or disconnect between transcription and translation during myogenic differentiation (Hunt et al. 2021).

### Prolonged Dexamethasone Exposure Better Mirrors *In Vivo* Atrophy Profiles

To assess how closely our C2C12 model mimics *in vivo* responses, we compared our proteomic results with a mouse muscle TMT-MS dataset from dex-treated mice (Hunt et al. 2021). When pooling both atrophy time points, overlap with the mouse dataset was limited. However, separate comparisons showed that the 72 hr C2C12 samples had greater concordance with the *in vivo* model (Fig. 5A), compared to the 24 hr samples (Supp. Fig. 4A).

**Figure 5:**
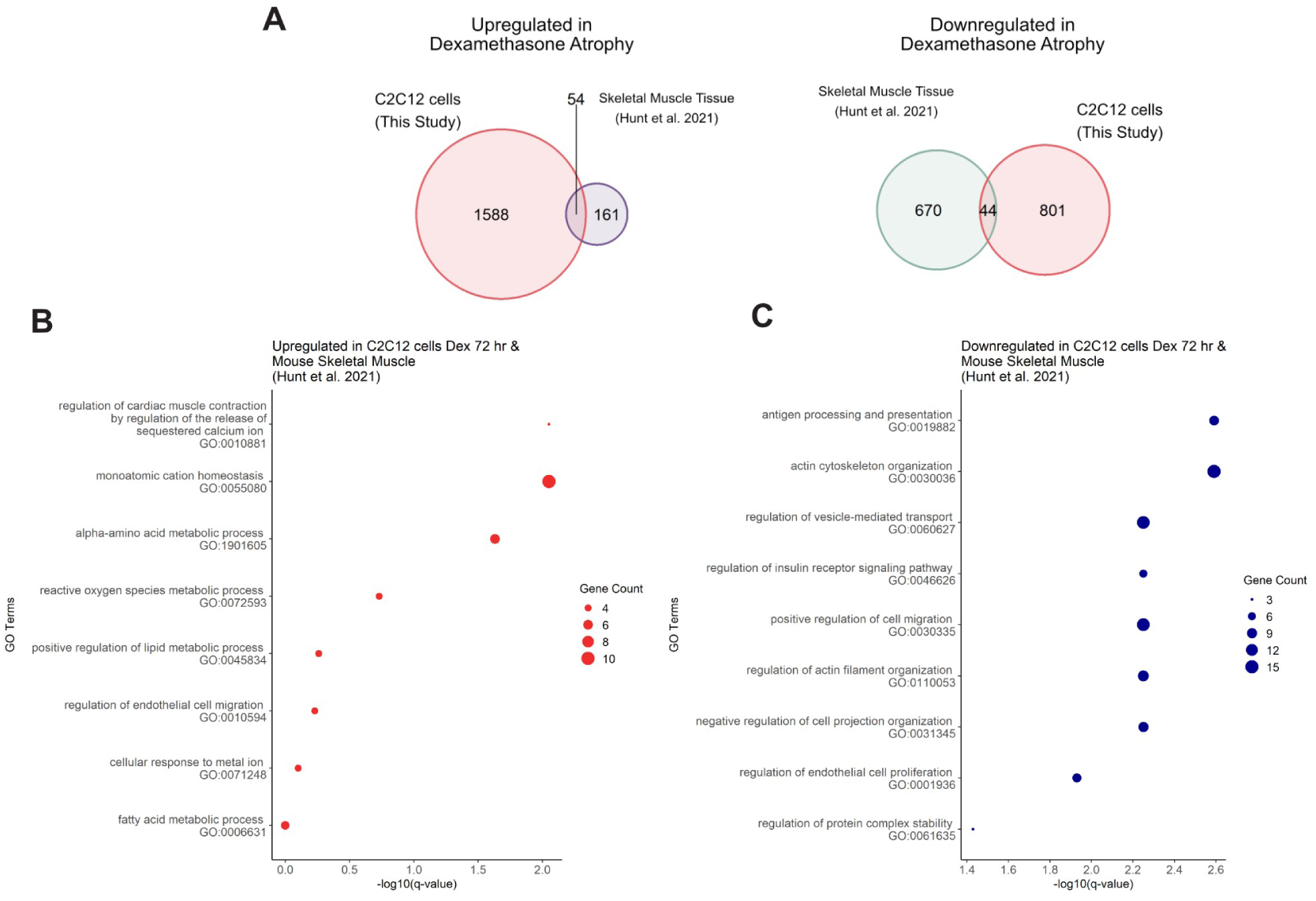
Gene Ontology (GO) analysis of dex-atrophy enriched proteins from this study and in comparison to published mouse skeletal muscle dex atrophy model. A) Overlap of proteins significantly changing in Dex <> Ctrl 72 hr in this study compared to published mouse skeletal muscle dex atrophy from Hunt et al. (2021) (adj.p-value ≤0.05 in both, -1≤log2FC≤0.6 applied for C2C12 TMT-MS dataset, -0.3≤log2FC≤0.3 applied for Hunt et al. TMT-MS data). B) GO analysis of 54 proteins increased in this study and Hunt et al. (2021) dataset in the left Venn diagram from A). C) GO analysis of 44 proteins decreased in this study and Hunt et al. (2021) dataset in the right Venn diagram from A).

We then performed GSEA to identify shared and unique biological processes. Common pathways enriched in both models included metabolic processes, such as that of monoatomic cation, alpha-amino acid, ROS, and lipid or fatty acids (GO: 0055080, GO:1901605, GO:0072593, GO:0045834, GO:0006631) (Fig. 5B). Downregulated pathways included actin cytoskeleton organization, vesicle-mediated transport, and cell migration properties (GO: 0030036, GO:0060627, GO:0030335) (Fig. 5C). However, C2C12-specific enrichments included mitochondrial processes and ribosome biogenesis (GO:0032543, GO:0007005, GO:0042254) (Supp. Fig 4B). Downregulation involved more processes related to the Golgi (Golgi vesicle transport, Golgi organization) and tube morphogenesis (Supp. Fig 4B). Interestingly, in mouse tissue, mitochondrial gene expression was suppressed, along with membrane organization and regulation of vesicle mediated transport, while no specific skeletal muscle related processes were activated (Supp. Fig. 4C). This highlights a biological mismatch between *in vitro* and *in vivo* systems.

## Discussion

Endogenous glucocorticoids (GC) hormones play an essential role in maintaining protein homeostasis in skeletal muscle, balancing protein synthesis and degradation. Clinically, GCs are commonly used to dampen inflammation via its inhibitory effects on the NF-κB pathway (Auphan et al. 1995, Schakman et al. 2013). However, chronic or high-dose GC exposure is known to induce muscle atrophy through transcriptional regulation via the GR, which leads to muscle protein degradation (Shimizu et al. 2011). While animal models have provided important insights into GC-induced muscle atrophy, cell culture model systems offer a scalable and controllable method to study these mechanisms in better resolution. Here, we present a comprehensive multi-omics dataset profiling transcriptomic, proteomic, and splicing dynamics in C2C12 myotubes undergoing dex-induced atrophy.

The C2C12 cell line is a widely used model for studying skeletal muscle development and atrophy. Our data indicate that extending differentiation to day 7, beyond the commonly used 3-5 day window, more faithfully replicates the mature sarcomere environment and better models *in vivo* atrophy responses. This was apparent based on the increased fusion index, myotube diameters, and enrichment of structural and contractile component characteristic for mature muscle. Since the C2C12 cell line cannot be reproducibly cultured past day 10 without loss of adherence or cultured past passage 12-15 without loss of differentiation properties, this defines a practical experimental window for modelling atrophy *in vitro*.

We observed a high degree of overlap between data from mouse tissue in mRNA and protein patterns during the transition of undifferentiated myoblasts to differentiated myotubes. This included enrichment of Z-disc components, thick and thin filaments, along with an increase of myogenic factors such as Myomaker, Myomixer, Myogenin and Myf5. These observations are further supported by measurements of cell morphology. Together, our results suggest that the mRNA and proteomic data from C2C12 cells provide robust information about fusion and differentiation hallmarks of muscle cells and will thereby aid further studies of sarcomere formation.

Upon dex-treatment, we detected an increase of mRNAs and proteins associated with mitochondrial protein expression whereas mRNAs for factors involved in DNA replication and mitosis were reduced. This seemingly contradicts previous findings from mouse tissue that implicated mitochondrial dysfunction as a hallmark of atrophy (Brown et al. 2017, Hunt et al. 2021). This could be explained by differences in the duration of dex-treatment (14 days in mice vs. 3 days in C2C12 cells) or dosage, especially given that lower dex administration in 4 day differentiated C2C12 myotubes showed compromised oxidative capacity (Liu et al. 2016). Our dataset also does not show downregulation of sarcomeric proteins despite expression of atrogenes (MuRF1, Atrogin-1, TRIM72) and GC responsive protein expression (TSC22D3, FKBP5). On the other hand, we also saw a significant decrease in cytosolic ribosome expression, which is in line with an impaired overall protein synthesis that has previously been observed in other atrophy models (Gordon et al. 2013). Presumably our dataset therefore captures earlier events leading up to sarcomeric degradation. Comparison of a mouse dex atrophy model (Hunt et al. 2021) indeed points towards this hypothesis, as we had the greatest overlap with our 72 hr dataset. Other studies also support that C2C12s may replicate “early” atrophy responses as defined in mice: A biphasic expression pattern has been reported for proteasomal subunits (RPN9, Gankyrin) and MuRF1 under denervation atrophy (Gilda et al. 2024) with similar transcriptional changes in cancer cachexia (Morena da Silva et al. 2023).

Although in this study, we defined 24 hr as “early” and 72 hr as “late” relative to C2C12 culture times, we still found clear differences in transcriptomic and proteomic signatures relative to controls. This suggests a dynamic response to atrophy stimuli already in what one may consider “early phase” in animal studies. Whether the degradation of sarcomeric components can then be captured from a day 5 myotube state to maximize atrophy treatment time and better align *in vivo* models would be of interest in future studies. This may also provide insight into transcription/translation cyclical rhythms explaining the biphasic atrophy response observed by others (Gilda et al. 2024, Morena da Silva et al. 2023).

Beyond differential expression, our transcriptomic analysis revealed widespread differential alternative splicing (DAS) at the early phase of atrophy, with fewer events at 72 hrs. Here, we observed a strong shift in isoform landscape without significant gene set enrichment in the GSEA analysis at 24 hrs. While we have not performed RNA-binding protein motif enrichment or functional validation, these isoform switches, especially in genes such as *Dcun1d3*, *Ubr2*, *Ubr5*, and *Mylk4*, may alter protein interactions, localization, and function. For example, the observed isoform-specific shift in *Ubr2* (leading to a 13-amino acid variation) and the reduction in UBR2 protein despite increased transcript levels point to complex post-transcriptional or translational regulation. Such transcription and translation mismatch with a layer of splicing regulation could contribute to the resulting stimulus specific atrophy phenotypes. The rest of the above-mentioned proteins have previously been associated with cachexia or protein anabolism with the exception of DCUN1D3 (Kwak et al. 2004, Gao et al. 2022, Hughes et al. 2021). Given published involvement of Cullin complexes in atrophy, we hypothesize that NEDD8 E3 ligases are involved in this phenotype (Bodine et al. 2001, Nowak et al. 2019, Blondelle et al. 2020).

A meta-analysis of array-based transcriptomic data from mice, rat and human tissues with corresponding cell lines revealed same species to have the best correlation (Abdelmoez et al. 2020). More importantly, C2C12 performed most comparable to human primary cells, in terms of its transcriptional profile for skeletal muscle function and contraction (Abdelmoez et al. 2020). Our data show that dex-treated C2C12 myotubes apparently represent early stages of muscle atrophy well while replicating later atrophy phenotypes poorly. We also identified new proteins to be possibly involved in atrophy, such as DCUN1D3, and shed light on alternative splicing events potentially worth further investigation. Thus, processes like recovery from sarcomere damage and proof of principle studies for atrophy prevention still ultimately rely on the analysis of muscle tissue – yet, for the identification of early indicators for chronic muscle wasting and initial design of therapeutic compounds to counteract muscle wasting, the C2C12 cell culture still has much to offer, saving time-consuming and expensive animal studies.

## Materials and Methods

### Cell culture and dexamethasone-induced atrophy

C2C12 cells (ATCC CRL1722) were cultured in growth medium (GM; 10 % fetal bovine serum (FBS; Gibco, 10270-106), 1 % Penicillin/Streptomycin (P/S; Sigma-Aldrich), high glucose Dulbecco’s modified Eagle’s medium (DMEM; Sigma-Aldrich) and maintained at 50-60 % confluency. For differentiation, cells were switched to differentiation medium (DM, 2 % FBS, 1 % P/S, high glucose DMEM) once they reached 80-90 % confluency (considered day 0) and cultured for 7 days with fresh DM supplied every 2-3 days. To induce atrophy, cells were treated with 100 µM dexamethasone (Sigma-Aldrich) or equivalent dilution of 99.99 % ethanol (Sigma-Aldrich) in DM and cultured for up to 3 more days. Replicates were prepared from various passages and stocks and cultured in a staggered manner to be considered biological replicates. Mycoplasma presence was checked intermittently using the LookOut Mycoplasma Detection PCR kit (Sigma-Aldrich).

### RNA extraction and quality control

Total RNA was extracted using Zymo Direct-zol RNA Miniprep Plus (ZymoResearch) according to manufacturer’s instructions. On collection days, cells were washed twice with D-PBS (Sigma-Aldrich) then lysed with the supplied lysis buffer for 5 min at room temperature before collection with a cell scraper. Samples were snap frozen and stored in -80 °C until extraction. Samples were thawed on ice and total RNA was extracted with all centrifugation steps performed at 20,817 x g and an additional empty centrifugation step before elution. RNA was eluted using supplied nuclease-free double distilled water. An aliquot was taken to measure A260/280 and A260/230 on a NanoDrop2000 (Thermo Scientific, PeqLab Biotechnologie GmbH). Ratios >1.9 for both measurements were deemed as contamination free. Where samples did not pass these criteria, RNA was precipitated using ethanol and resuspended in nuclease-free water. RNA integrity was checked using BioAnalyzer 2000 and RNA 6000 Nano kit with the RNA 6000 ladder (Agilent). A RIN score of >8.0 was chosen as the passing score for subsequent library preparation using the TruSeq Stranded mRNA kit (Illumina) according to manufacturer’s instructions.

### Bulk RNA sequencing

RNA sequencing was performed using the NovaSeq 6000 at the MDC Genomics Technology Platform. Samples were sequenced at S1 mode using paired-end sequencing with 2×100 bp read length and dual indexing. Samples were demultiplexed and read length quantified using salmon, via pseudoalignment against all transcripts and on reference chromosomes in the gencode vM12 mouse transcriptome annotations using parameters: “salmon quant –I ISR –seqBias –gcBias – validateMappings”. RNA-just seq fold changes were calculated using the tximport package with default settings for isoform aware quantification, and DEseq.

### Differential Alternative Splicing and Isoform Switch Analysis

All computational analyses were performed using R (version 4.4.1) and relevant packages as described: Raw RNA-seq reads were pseudo-aligned and quantified using Kallisto (version 0.51.1) with the mouse reference transcriptome corresponding to genome assembly GRCm39. The resulting transcript-level abundance estimates (TPM and estimated counts) were imported into R using the importIsoformExpression() function from the IsoformSwitchAnalyzeR package (v 2.6.0). Isoform-level analysis was also conducted using IsoformSwitchAnalyzeR (version 2.6.0). A switchAnalyzeRlist object was created from Kallisto outputs. Isoform switches were identified between conditions, and potential functional consequences of switching events. The default thresholds of isoform_switch_q_value < 0.05 and differential isoform fraction abs(dIF) > 0.1 were used unless otherwise stated. For the comparison with the Differentially Expressed Genes a threshold of p adjusted value padj<0.05 was used.

### Tandem Mass Tag (TMT) Mass Spectrometry Sample Preparation

On collection days, cells were washed three times with phosphate-buffered saline (PBS), scraped with 1 mL D-PBS into a 1.5 mL microcentrifuge tube, and centrifuged at 500 × g for 3 min at 4°C. The supernatant was discarded, and the cell pellet was snap-frozen in liquid nitrogen before storage at –80°C. For protein extraction, cell pellets were lysed in 8 M urea lysis buffer containing 75 mM NaCl, 50 mM Tris-HCl (pH 8.0), 1 mM EDTA, 2 μg/mL aprotinin, 10 μg/mL leupeptin, 1 mM PMSF, 1:100 (v/v) Phosphatase Inhibitor Cocktail 2, 1:100 (v/v) Phosphatase Inhibitor Cocktail 3, 10 mM NaF, and 20 μM PUGNAc. Lysates were sonicated on ice using a probe sonicator at 20 % amplitude in three cycles of 10 sec on and 20 sec off. Insoluble debris was removed by centrifugation at 14,000 × g for 10 min at 4 °C. Protein concentration was determined using the bicinchoninic acid (BCA) protein assay kit (Pierce, Thermo Fisher Scientific) according to the manufacturer’s instructions.

### TMT Labelling and Peptide Digestion

For each sample, 1 mg of protein was reduced with 5 mM dithiothreitol (DTT) at 37 °C for 45 min, alkylated with 15 mM iodoacetamide (IAA) in the dark for 30 min at room temperature, and quenched with 5 mM DTT for 15 min. Proteins were sequentially digested with Lys-C (1:100 enzyme-to-protein ratio) for 3 h at 37 °C, followed by trypsin digestion (1:50 ratio) overnight at 37 °C. Peptides were desalted using Sep-Pak C18 cartridges (Waters) and dried in a vacuum concentrator. Tandem Mass Tag (TMTpro) 16-plex labelling was performed using two sets of TMTpro reagents (Thermo Fisher Scientific) following the protocol described by Mertins et al. (2018). Each TMT 16-plex experiment included a balanced representation of time points, with one channel designated for a pooled reference sample containing equal proportions of peptides from all other channels. Labelled samples were pooled, desalted, and fractionated using high-pH reversed-phase liquid chromatography on an UltiMate 3000 HPLC system (Thermo Scientific) equipped with an XBridge Peptide BEH C18 column (130 Å, 3.5 µm, 4.6 mm × 250 mm; Waters). Peptides were separated into 24 concatenated fractions.

### Liquid Chromatography-Tandem Mass Spectrometry (LC-MS/MS) Analysis

Peptide fractions were resuspended in 0.1 % formic acid and analysed using an EASY-nLC 1200 system coupled to an Orbitrap Exploris 480 mass spectrometer (Thermo Fisher Scientific). Peptides were separated on a 50 cm × 75 μm inner diameter EASY-Spray column (Thermo Fisher Scientific) packed with 1.9 μm C18 beads (PepMap RSLC). A 110 min HPLC gradient was applied at a flow rate of 250 nL/min. Data acquisition was performed in data-dependent acquisition (DDA) mode, with MS/MS spectra collected at a resolution of 45,000 and an isolation width of 0.4 m/z.

### Analysis of TMT-MS Data

Raw data files were processed using MaxQuant (v.1.6.10.43) and searched against the Mus musculus proteome (2018-07) downloaded from UniProt, including isoforms. Trypsin/Lys-C was specified as the digestion enzyme, with up to two missed cleavages allowed. Carbamidomethylation of cysteine was set as a fixed modification, while oxidation of methionine and acetylation of protein N-termini were specified as variable modifications. A PIF filter of 0.5 was applied. Peptide-spectrum matches (PSMs) and protein identifications were filtered at a false discovery rate (FDR) of 1 %. Only proteins with at least two unique peptides and valid reporter ion intensities across all samples were retained for further analysis.

Protein abundances were calculated from log₂-transformed, corrected TMT reporter ion intensities. Intensities were normalised against the internal reference, followed by sample-wise median-median absolute deviation (MAD) normalisation to correct for systematic variations. Differential protein expression analysis was performed using the limma R package (Ritchie et al. 2015), with Benjamini–Hochberg correction applied for multiple testing. Gene lists belonging to indicated GO terms were retrieved from the Mouse Genome Database (MGD), Mouse Genome Informatics, The Jackson Laboratory, Bar Harbor, Maine. (http://www.informatics.jax.org).

### Gene Set Enrichment Analysis, GO enrichment and Protein-Protein Interaction Enrichment Analysis

For GSEA of RNA data, DESeq2 generated pairwise comparison data was enriched using ClusterProfiler (ver. 4.14.6, Wu et al. 2021) and the DOSE packge (ver. 4.0.1, Yu et al. 2014). Gene lists for significantly increased and decreased proteins were generated using adjusted p-value ≤0.05 and/or -0.6≤logFC≤1 from our TMT-MS data to account for fold-change by 50 %. Protein-protein interaction enrichment analysis was then performed using Metascape (Zhou et al., 2019) and visualized in Cytoscape (Shannon et al., 2003). GO enrichment for protein data was performed using string.db (ver. 12.0) (Szklarczyk et al. 2023).

### RT-qPCR

Aliquots of total RNA isolated for the bulk RNA-sequencing was used to validate relative changes in quantity of certain transcripts. 1 µg of total RNA was reverse transcribed into cDNA using Maxima H Minus (VWR Technologies) and oligo-dT_20_ primer per reaction. Total RNA was incubated with 10mM dNTPs at 65 °C for 5 minutes, then briefly placed on ice. A mix of 5x RT buffer provided with Maxima H Minus was mixed with RiboLock RNase Inhibitor (20 U per reaction, Life Technologies) and added before incubation at 50 °C for 30 minutes. The cDNA mixture was then diluted 1:20 using nuclease-free water (Promega).

Bio-Rad CFX96 was used to quantify SYBR Green-mediated amplification signals. In a 96-well plate (Biozym), cDNA was pipetted with RT-qPCR primers and 2x GoTaq qPCR master mix (Promega). Water blank was used to check for primer-primer dimer formation. Plates were cycled as follows: 95 °C for 2 min, 41x loops of 95 °C for 3 seconds, 62 °C for 30 seconds, ending with a 60-90 °C temperature gradient of 1 °C increment. Plates were read after the 62 °C annealing step and again after the temperature gradient. Signals were calculated using the 2^-ΔΔCt^ method with more than one house-keeping gene. Primers used are listed in the following table:

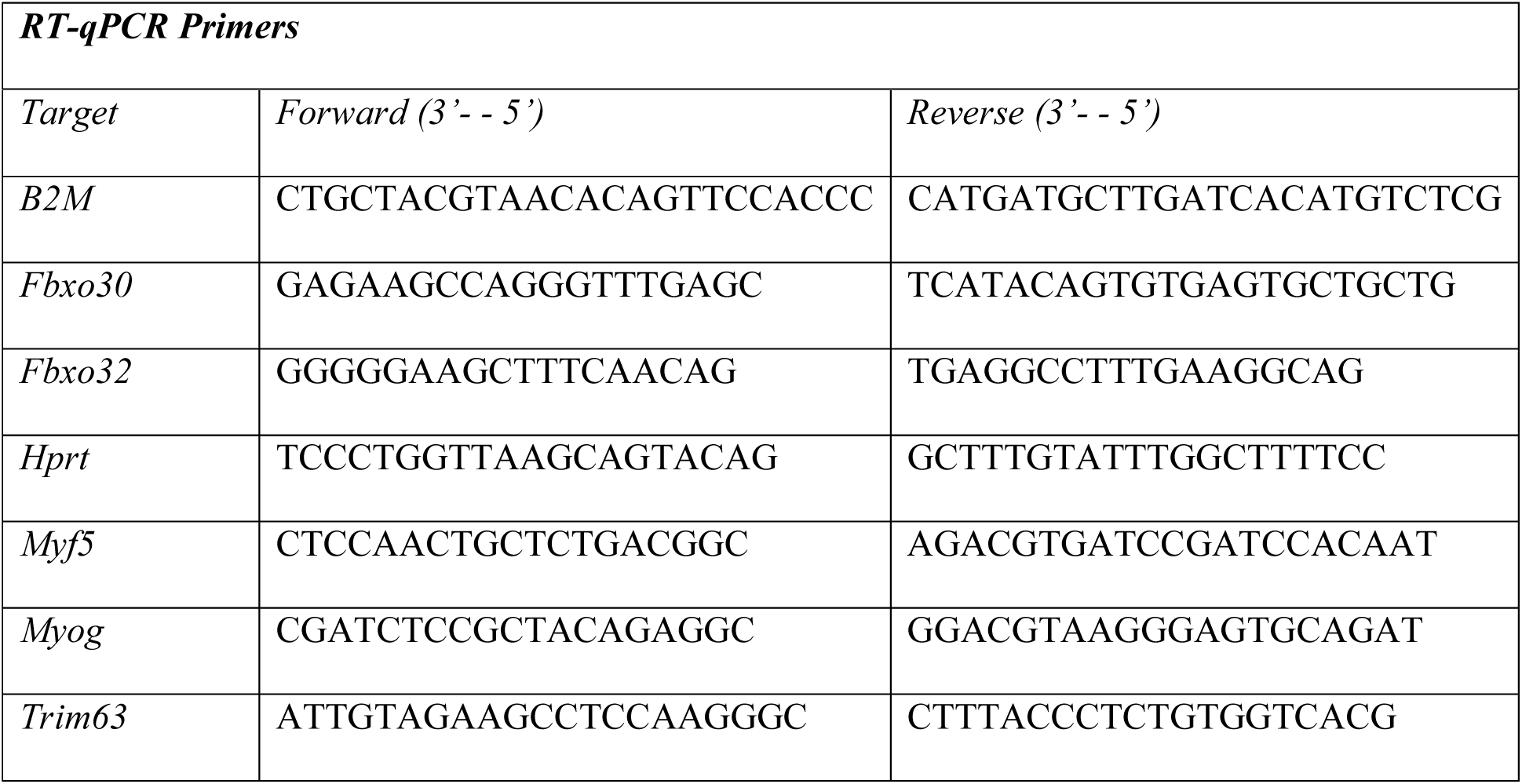

### Immunofluorescence, Fusion Index and Diameter Measurements

Cells were cultured and treated for atrophy induction in ibiTreat 35 mm dishes (ibidi). On days matching the bulk RNA-seq and TMT-MS samples, dishes were washed with D-PBS(-) and fixed with 4 % PFA for 30 min at room temperature. Dishes were washed and permeabilized using 0.4 % Triton X-100 0.5 % BSA D-PBS(++) for 30 min at room temperature. Cells were then washed with 0.5 % Tween-20 D-PBS(++) henceforth and blocked in 0.05 % Triton-X100 3 % BSA D-PBS(++) for >30 min. After washing, samples were incubated overnight with antibodies against fast/type-II myosin heavy chain (MyHC; sigma, M4276) and α-actinin 2 at 4 °C. Following another wash, dishes were incubated with AlexaFluor 568 goat anti-rabbit (Invitrogen) or Cy5 goat anti-mouse (Invitrogen, A10524) at 2 µg/ml concentration in blocking buffer for 1 hr at room temperature. Cells were washed again, before incubation with Hoechst 33342 diluted to 1 µg/ml in D-PBS(++) for 5 min at room temperature. Cells were washed and stored in D-PBS(++) and imaged on a Leica SP8 confocal microscope. All images were analysed using Fiji (Schindelin et al. 2012).

Fusion index was calculated as the number of nuclei inside a MyHC positive cell out of the total number of nuclei in a field of view. Diameter measurements were carried out on MyHC positive cells in the vehicle control or atrophy conditions. Statistical significance comparing the diameter of all conditions was carried out using the Kruskal-Wallis test.

### Data Exploration, Statistical Analyses and Visualization

Data visualization and students moderated t-test for RNA-seq, DAS, isoform switch analyses and TMT-MS data were performed using base R functions, ggplot2 (version 3.5.1), pheatmap (version 1.0.12). Custom plots of gene expression, isoform expression, and isoform usage were generated from processed data. Venn diagrams were plotted using the VennDiagram package (version 1.7.3). Statistical testing and visualization of quantified immunofluorescence data for cell morphology were performed using GraphPad Prism 10 (version 10.4.1).

## Acknowledgements

Authors would like to Corinna Volkwein and Mandy Gerlach for excellent technical help and Benjamin Sünkel for scientific discussion. We also thank the Advanced Light Microscopy facility (MDC) for assistance in confocal imaging; Dr. Tatiana Borodina and Dr. Daniele Yumi Sunaga-Franze at the genomics sequencing facility (MDC) for assistance in RNA-seq and data pre-processing; Prof. Dr. Uta Höpken (MDC) for access to the BioAnalzyer; Dr. Cristina Brischetto (TU-Berlin) for advice on RT-qPCR experiments. We also thank Dr. Dermot Harnett and Prof. Dr. Uwe Ohler (MDC) for data analysis assistance.

## Data availability statement

The original contributions presented in the study are included in the article and supplementary material. RNA-Seq data were deposited into the Gene Expression Omnibus database under accession number [SUBMISSION ID PENDING] and can be accessed from the following URL: [URL PENDING]. The mass spectrometry proteomics data have been deposited to the ProteomeXchange Consortium (http://proteomecentral.proteomexchange.org) via the PRIDE partner repository (Perez-Riverol et al. 2024) with the dataset identifier PXD064871.

## Author Contributions

SN designed and performed experiments, analyzed discussed data, prepared, and edited figures and tables, prepared, and edited the manuscript. AM and OP analysed discussed data, prepared and edited figures and tables. EJ prepared and edited the manuscript. PM conceptualized and supervised the project. TS and JF conceptualized and supervised the project, designed and analyzed experiments, discussed data, edited figures and tables, prepared, edited the manuscript, and acquired funding. All authors edited and approved the final manuscript.

## Online supplementary material

Additional supporting information may be found online in the Supporting Information Section at the end of the article.

---

**Supplementary Figure 1.**
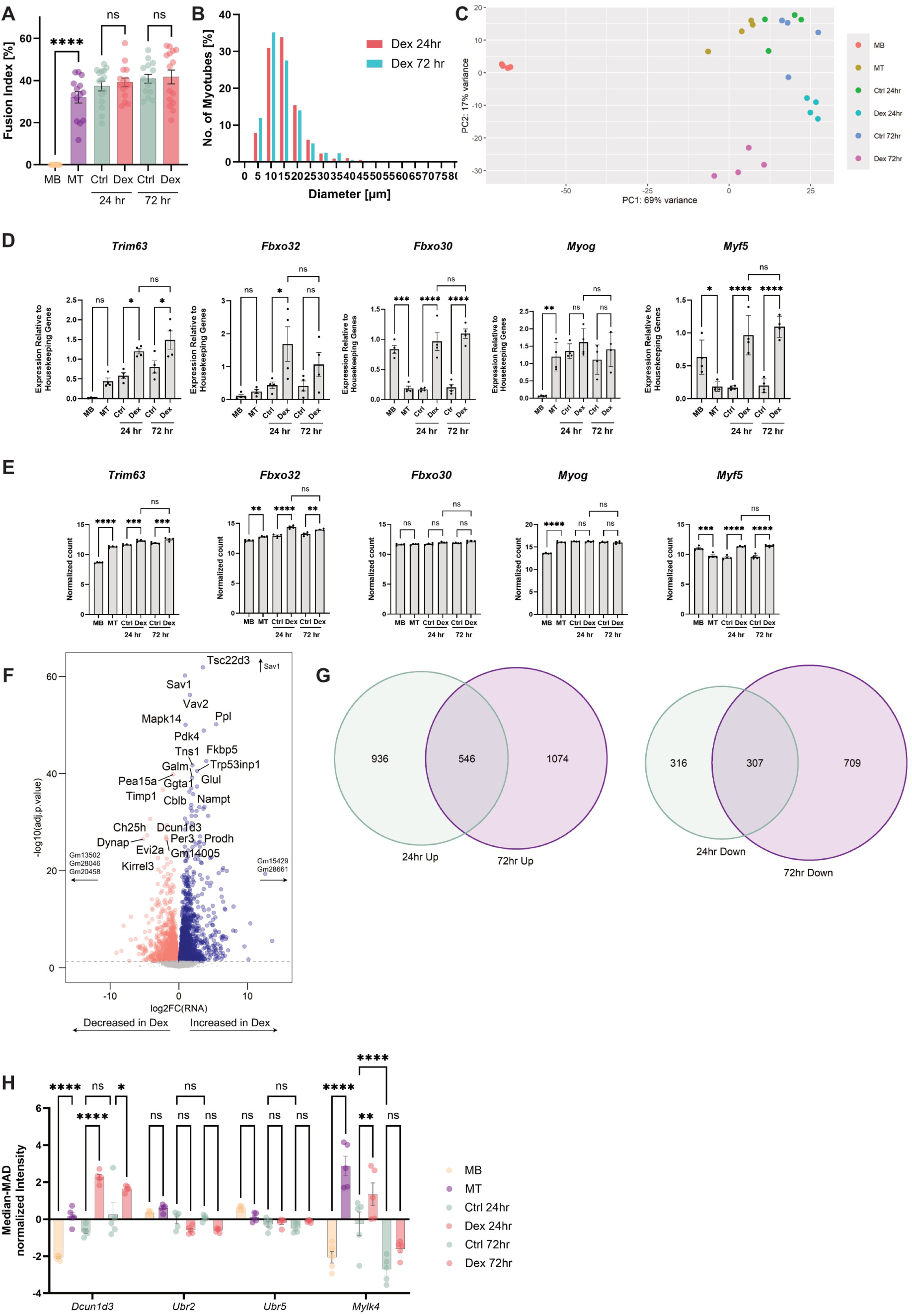
A) Fusion index of C2C12 cells at collection time points. Significance was tested using one-way ANOVA to compare means. n=3. ****<0.0001. B) Frequency distribution of MyHC positive C2C12 diameter measurements between Dex 24 hr and 72 hr. Values are indicated as % of total. C) PCA plot of bulk mRNA-seq using raw count intensity with the DESeq2 package. D) and E) RT-qPCR measurements (D) of indicated genes using the same cDNA samples submitted for sequencing, plotted in TPM (E). n=4 per time point. Significance was tested using one-way ANOVA and Tukey’s post-hoc test. Non-significant results have been omitted for legibility unless deemed relevant. *<0.05, **<0.01, ***<0.001, ****<0.0001. F) Volcano plot of transcripts at Dex <> Crl 24 hr. Grey dashed line indicates adj.p-value =0.05. G) Venn diagram of significantly up/downregulated genes in Dex <> Ctrl 24 hr and 72 hr. H) TMT-MS measured protein levels of genes undergoing DEG and DAS. Values have been median-MAD normalized and tested for significance using two-way ANOVA. *<0.05, **<0.01, ****<0.0001.

**Supplementary Figure 2.**
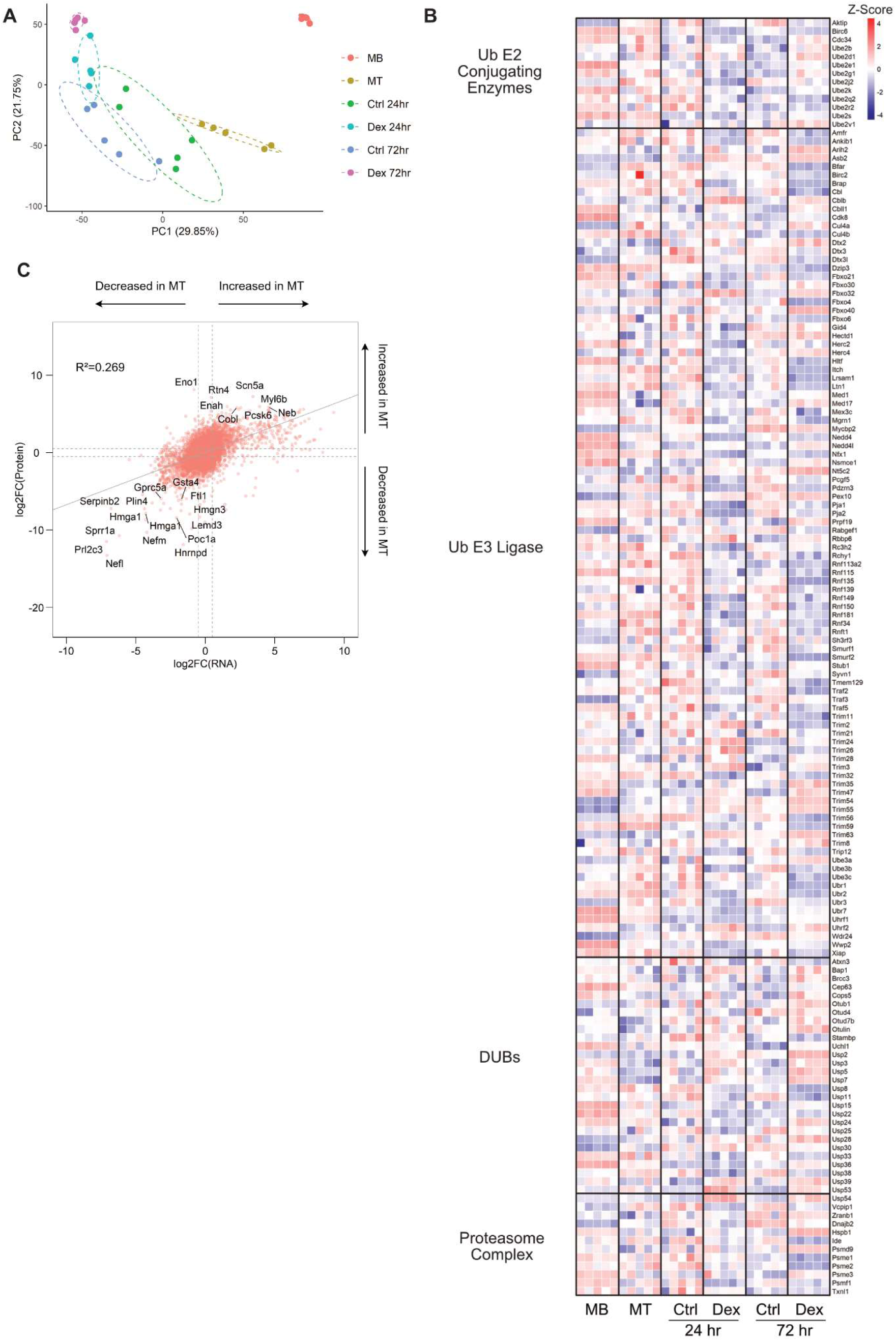
A) PCA plot of TMT-MS reporter intensity. B) Z-score heat map of TMT-MS data highlighting ubiquitin (Ub) E1∼E3 enzymes, DUBs and components of the proteasome complex as searched on the MGI database, with adj.p-value ≤0.05 in comparisons between Dex<> Ctrl 24 hr or 72 hr or Dex 24 hr <> Dex 72hr. None of the Ub E1 enzymes passed the significance filter. C) Log2FoldChanges of RNA (x-axis) plotted against those of proteins identified by TMT-MS (y-axis) for MT<>MB. R^2^ values were calculated using linear regression modelling.

**Supplementary Figure 3.**
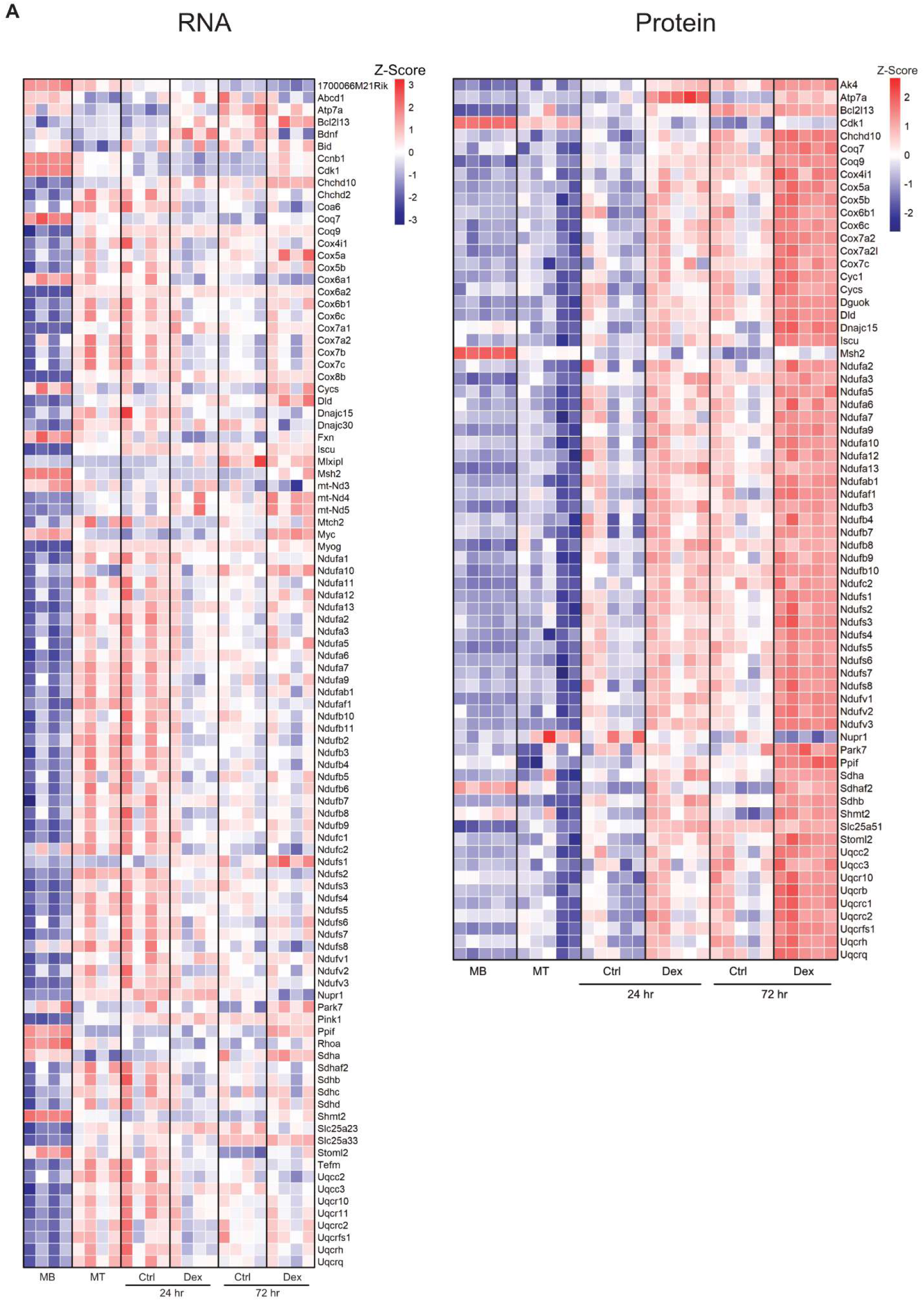
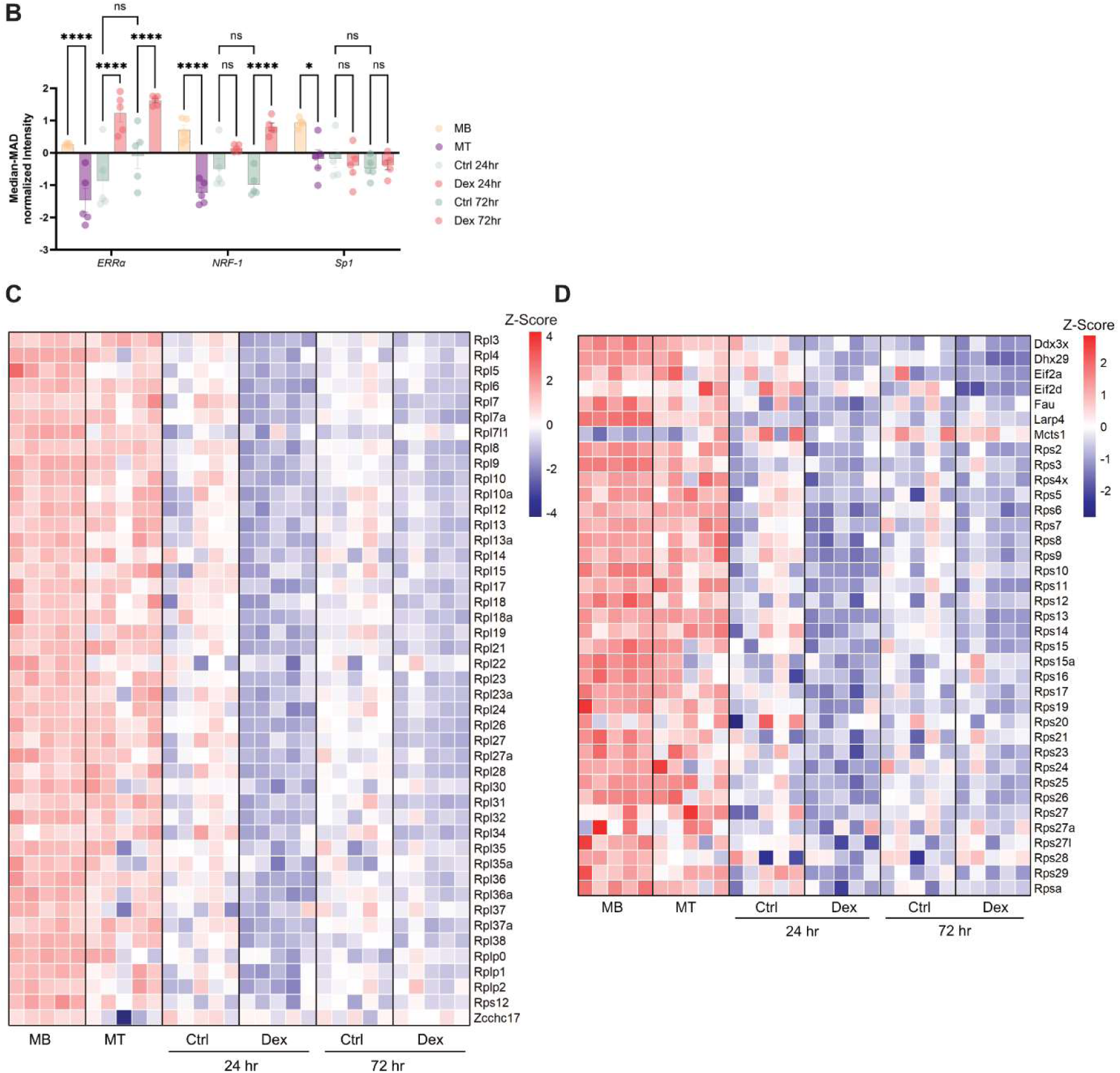
A) Z-score heat map of proteins associated with oxidative phosphorylation from RNA-seq (left) and TMT-MS data (right) which passed the adj.p-value ≤0.05 filter in comparisons of Dex <> Ctrl at 24 hr and 72 hr. RNA data has additional genes which were significant (adj.p≤0.05) in MT <> MB comparisons and protein has additional genes which were significant in Dex 24 hr vs Dex 72 hr comparisons. B) Median-MAD normalized intensity of transcription factors responsible for mitochondrial gene expression. TMT-MS data were filtered for the following transcription factors: NRF-1, NRF-2, PPARα, ERRα, Sp1, PGC-1α, PGC-1β, and PRC (Scarpulla et al. 2008). Significance testing with two-way ANOVA. *<0.05, **<0.01, ***<0.001, ****<0.0001. C) and D) Z-score heat map of cytosolic large ribosomal subunits (C) and small ribosomal subunits (D from TMT-MS data.

**Supplementary Figure 4.**
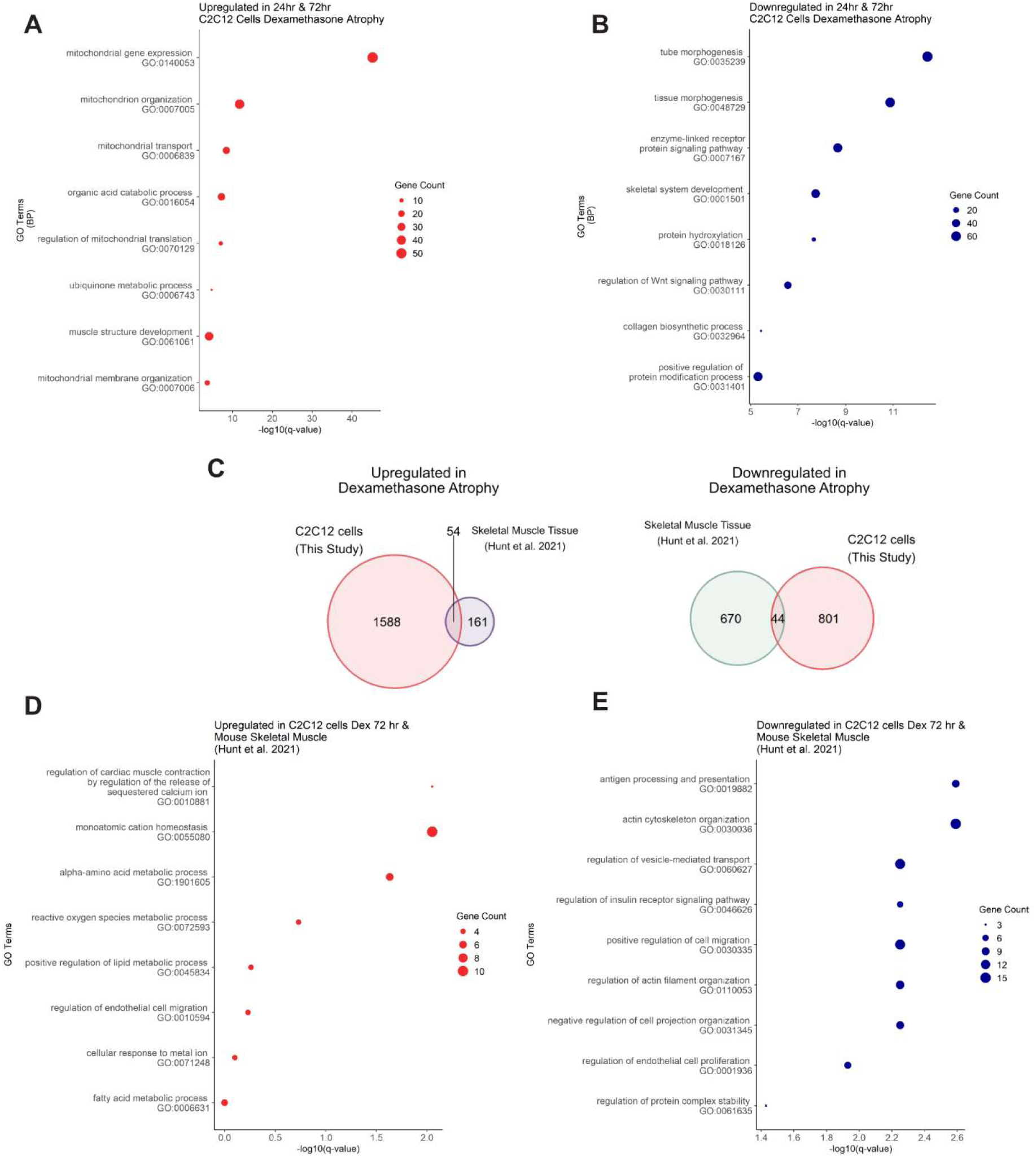
A) Venn diagram of significantly changing proteins (adj.p-value≤0.05) from TMT-MS data between this study, at the 24 hr time point and Hunt et al. (2021) dex data. B) GO analysis of up (red, left) or down (blue, right)-regulated proteins specific to C2C12 dex atrophy model from this study. C) GO analysis of up (red, left) or down (blue, right)-regulated proteinss specific to dex atrophy model from Hunt et al. (2021).

## Supplementary Tables

Supplementary Table 1: GSEA of RNA-Seq data for MT over MB and Dex over Ctrl 72 hr

Supplementary Table 2: Genes undergoing differential expression and DAS

## The Paper Explained

### Problem

Sarcomeric protein degradation and signaling pathway activation during skeletal muscle atrophy show high variability depending on mode of atrophy and model used. Comparison of mouse dataset and commonly used differentiated murine immortalized myoblasts using the same atrophy stimulus and similar screening technique is missing.

### Results

We found robust recapitulation of differentiation hallmarks at the mRNA and protein level, as well as atrophy markers without sarcomeric protein degradation in our cellular atrophy model. We also observed significant increase in mitochondrial protein expression, decrease in cytosolic ribosomes and alternative splicing events occurring as an early response to dexamethasone treatment.

### Impact

The data highlights the benefits and limitation of the commonly used cell line model, with an additional regulatory layer occurring early in atrophy. This would benefit exploration of new therapeutic opportunities and improve designing of experiments to complement animal studies.

---

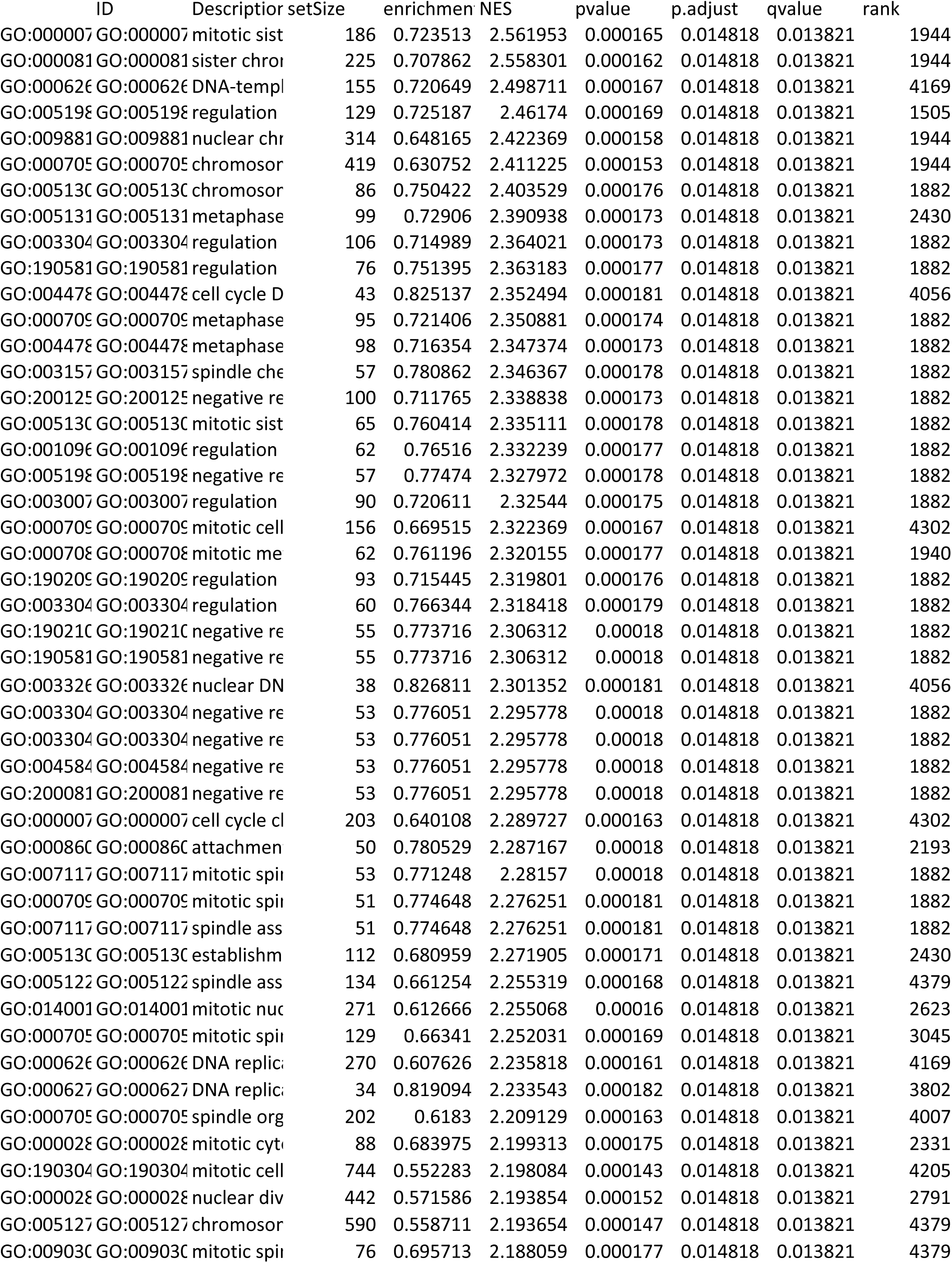
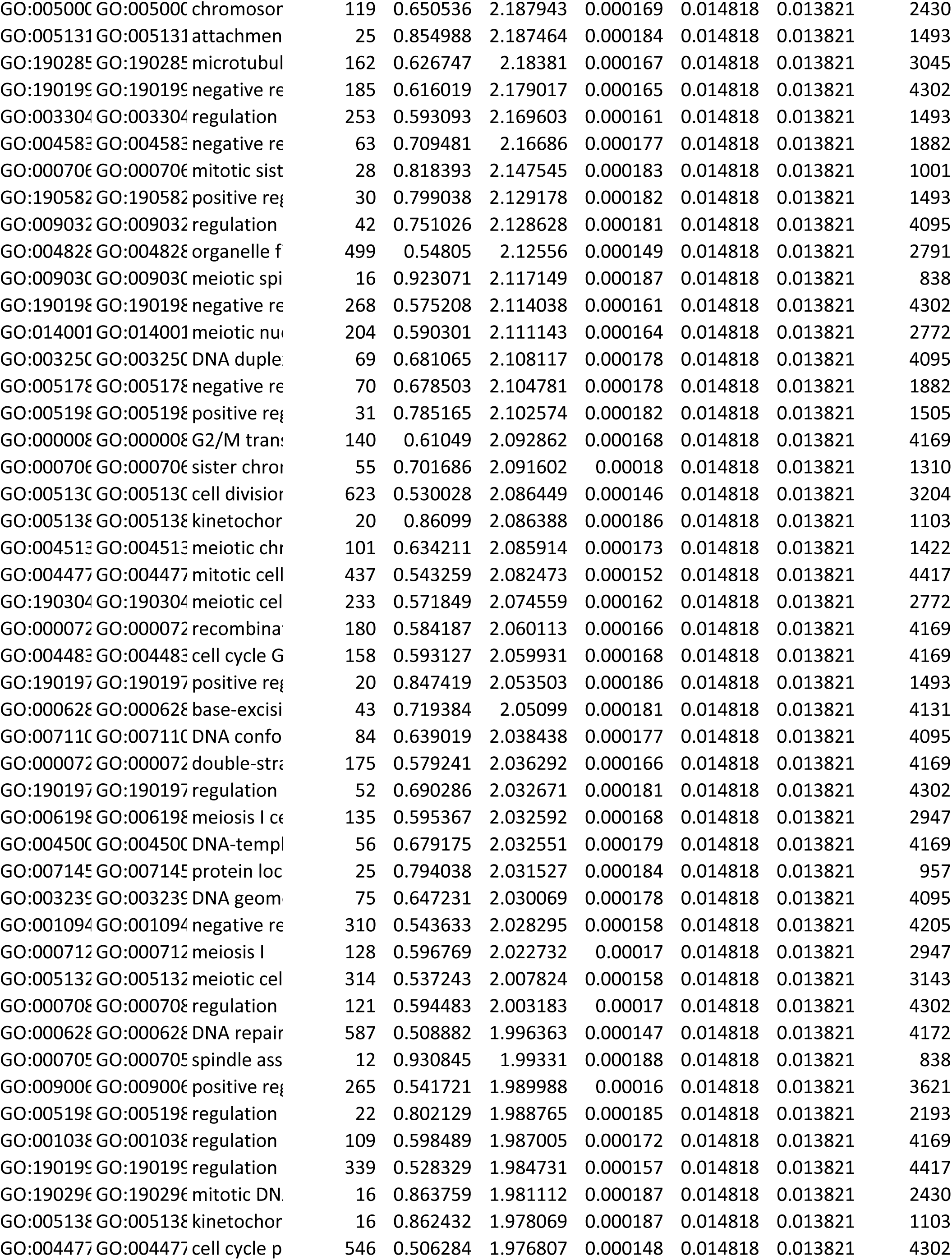
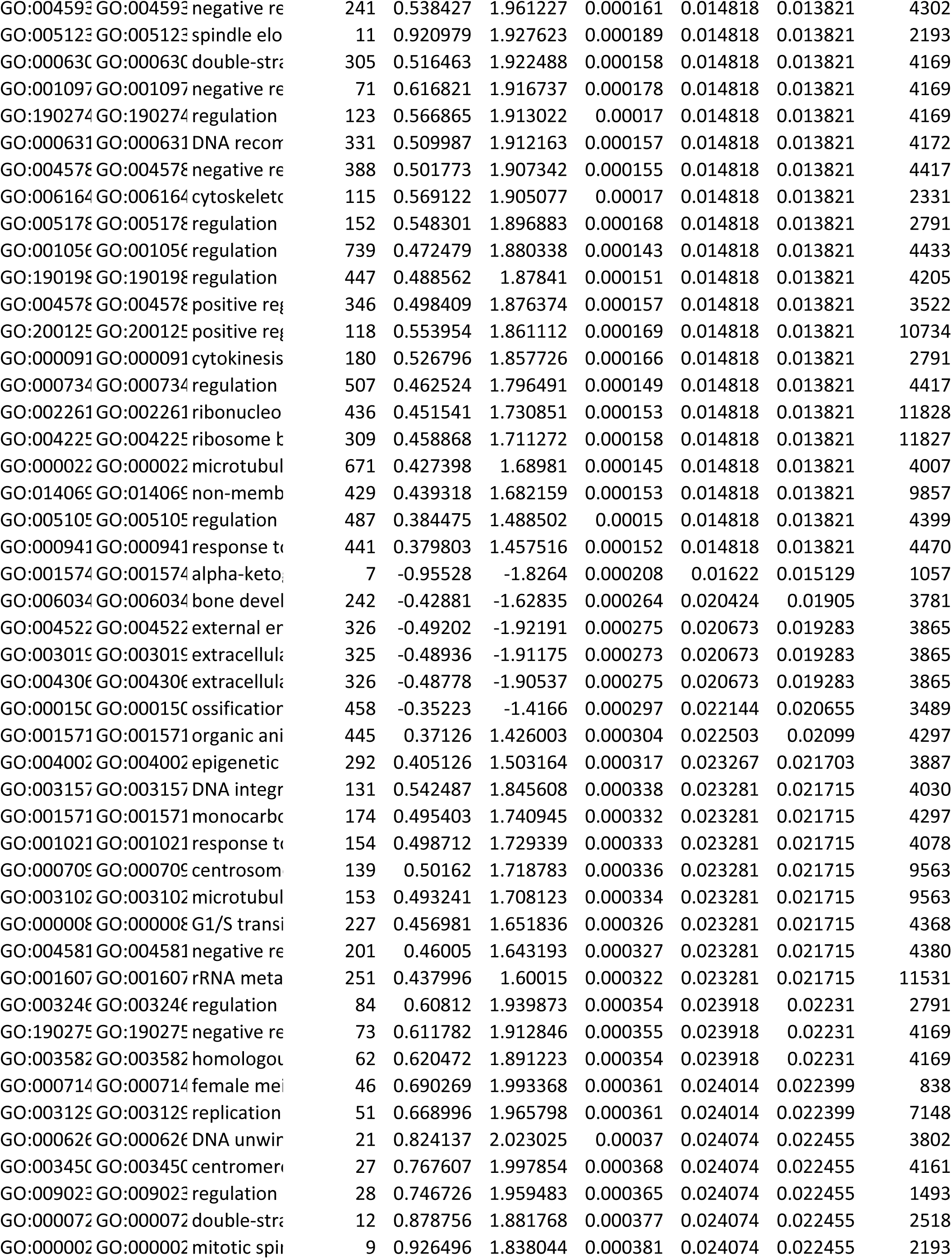
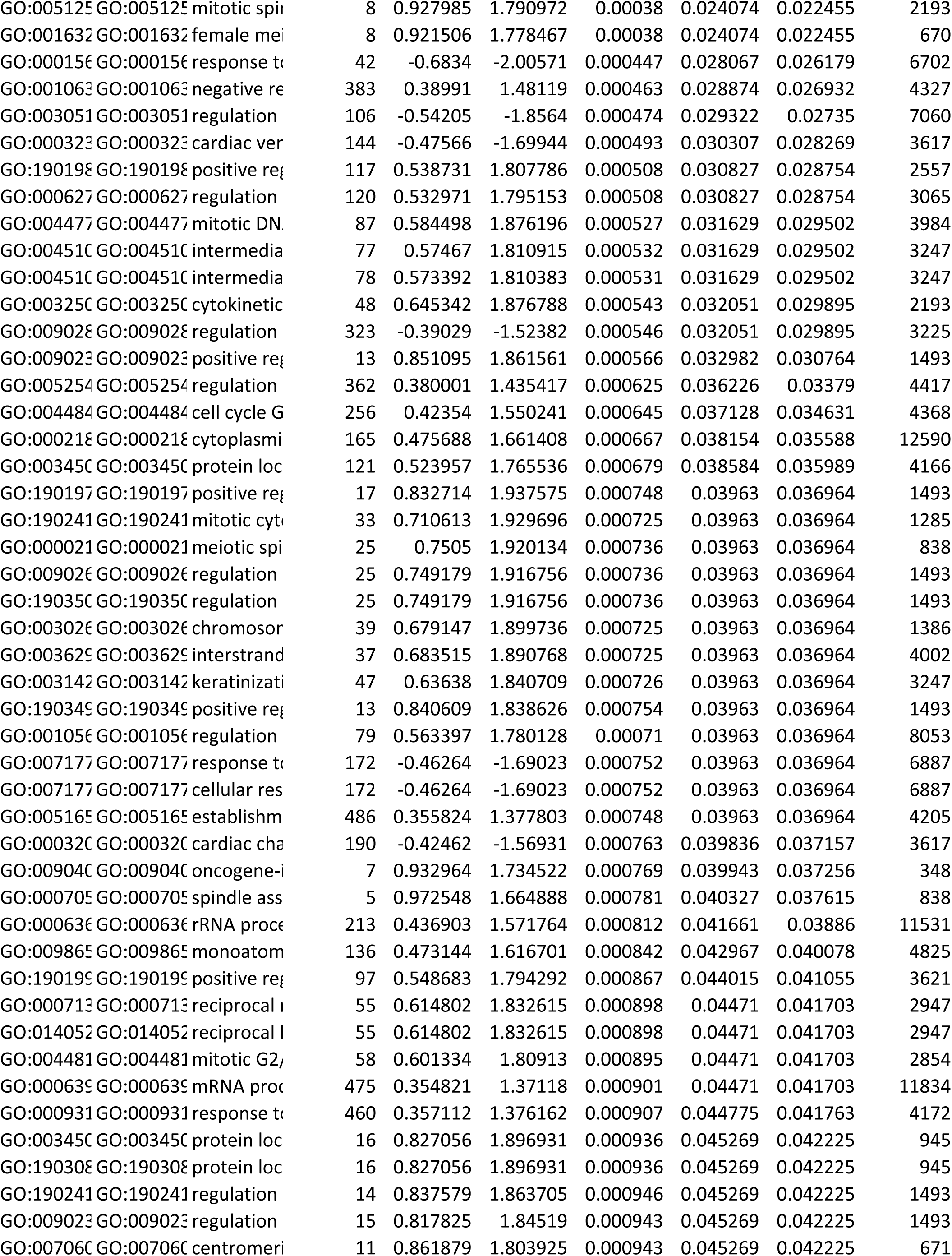
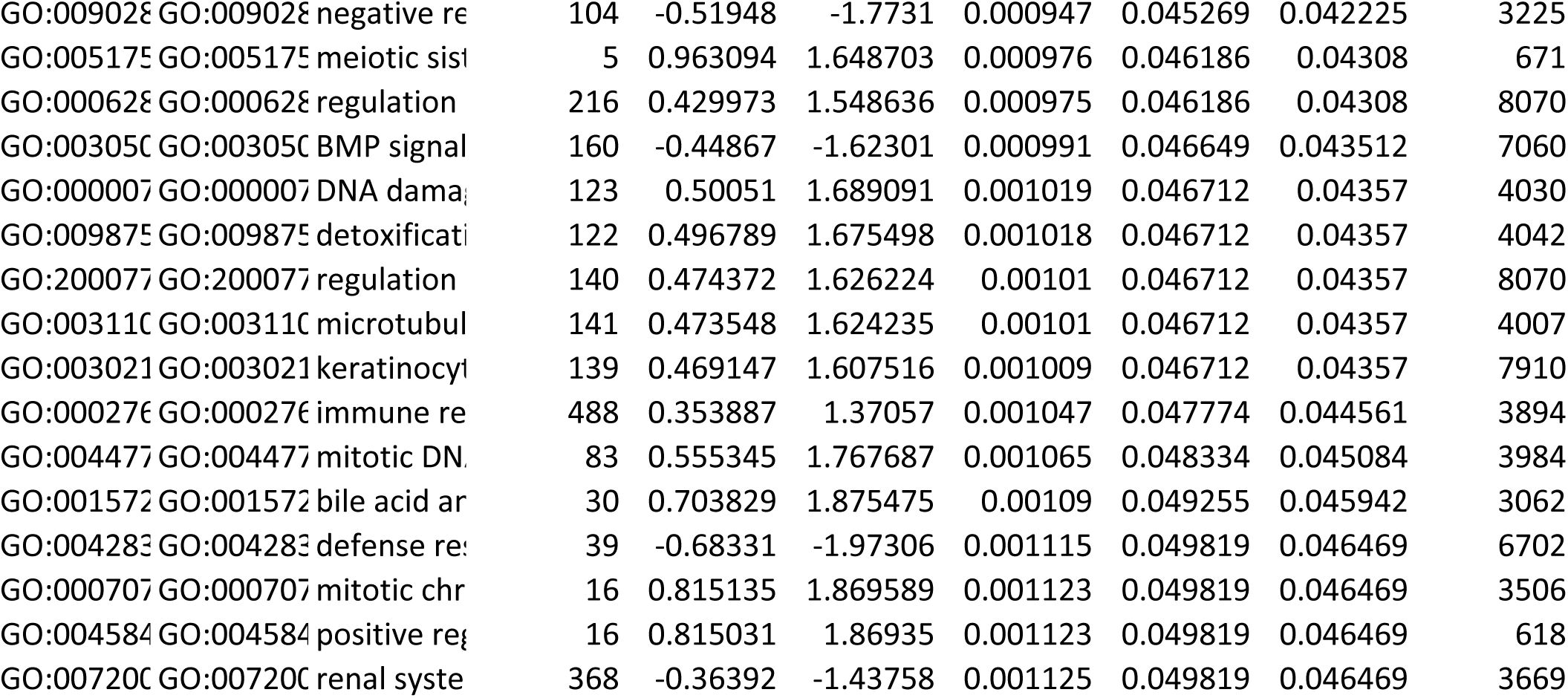

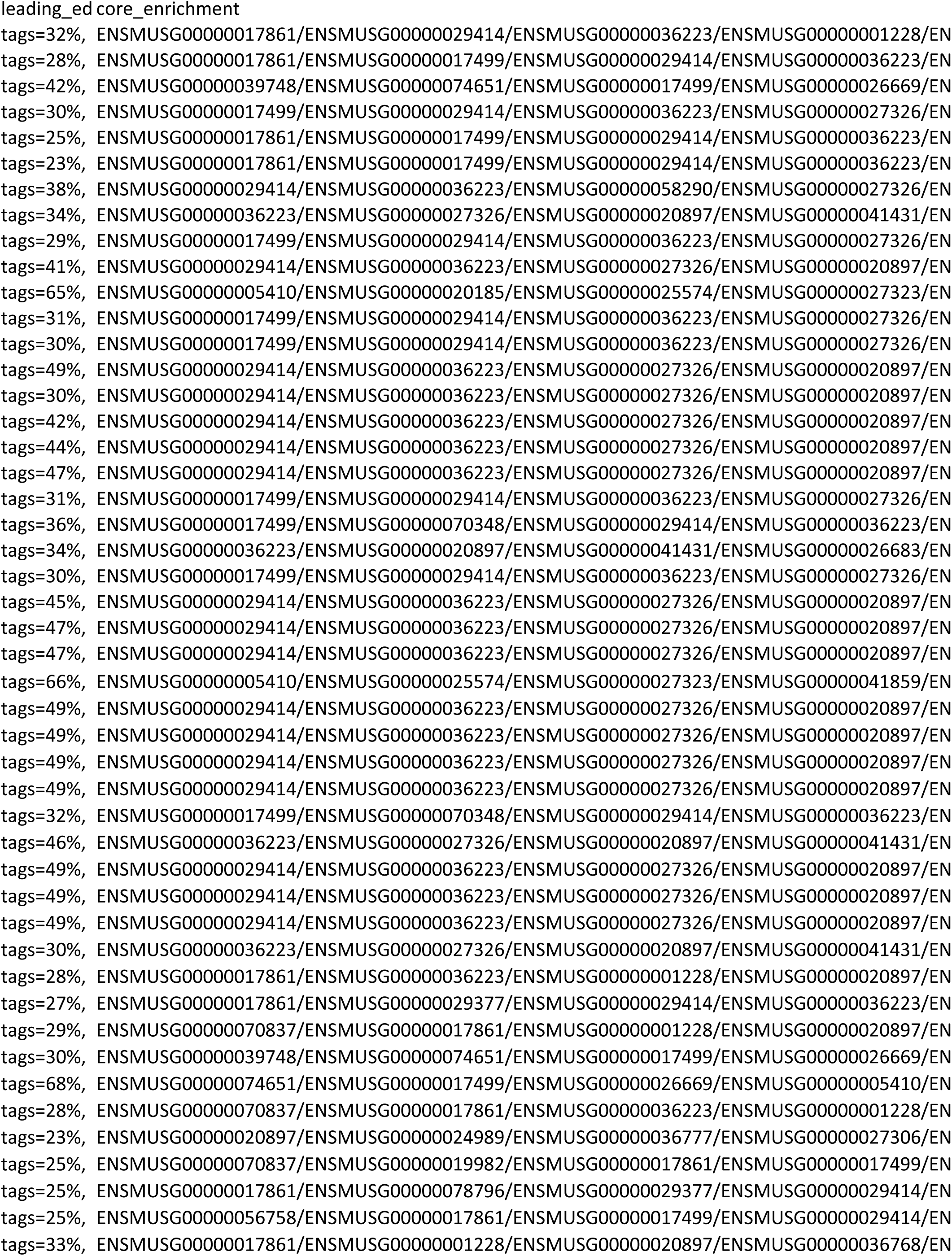
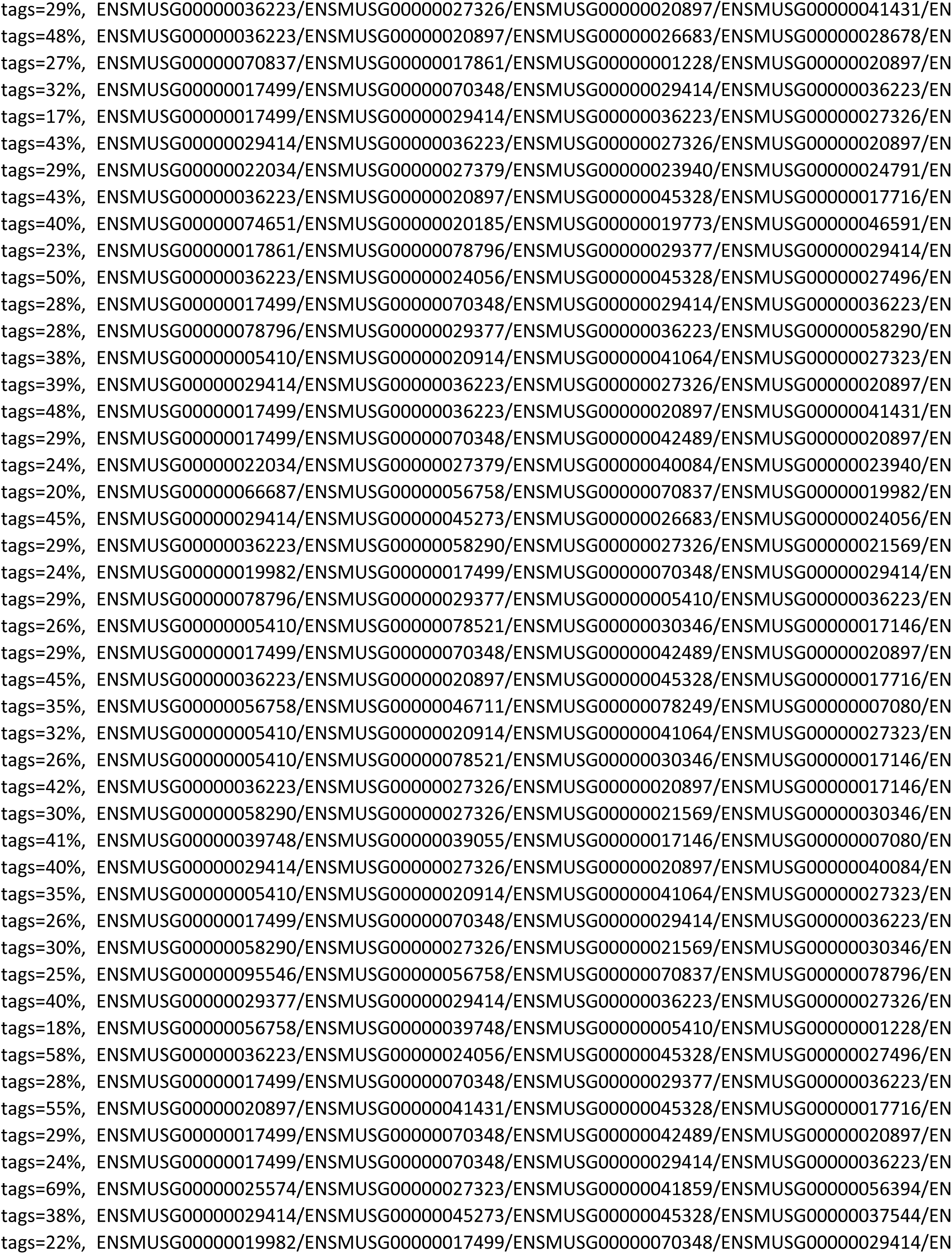
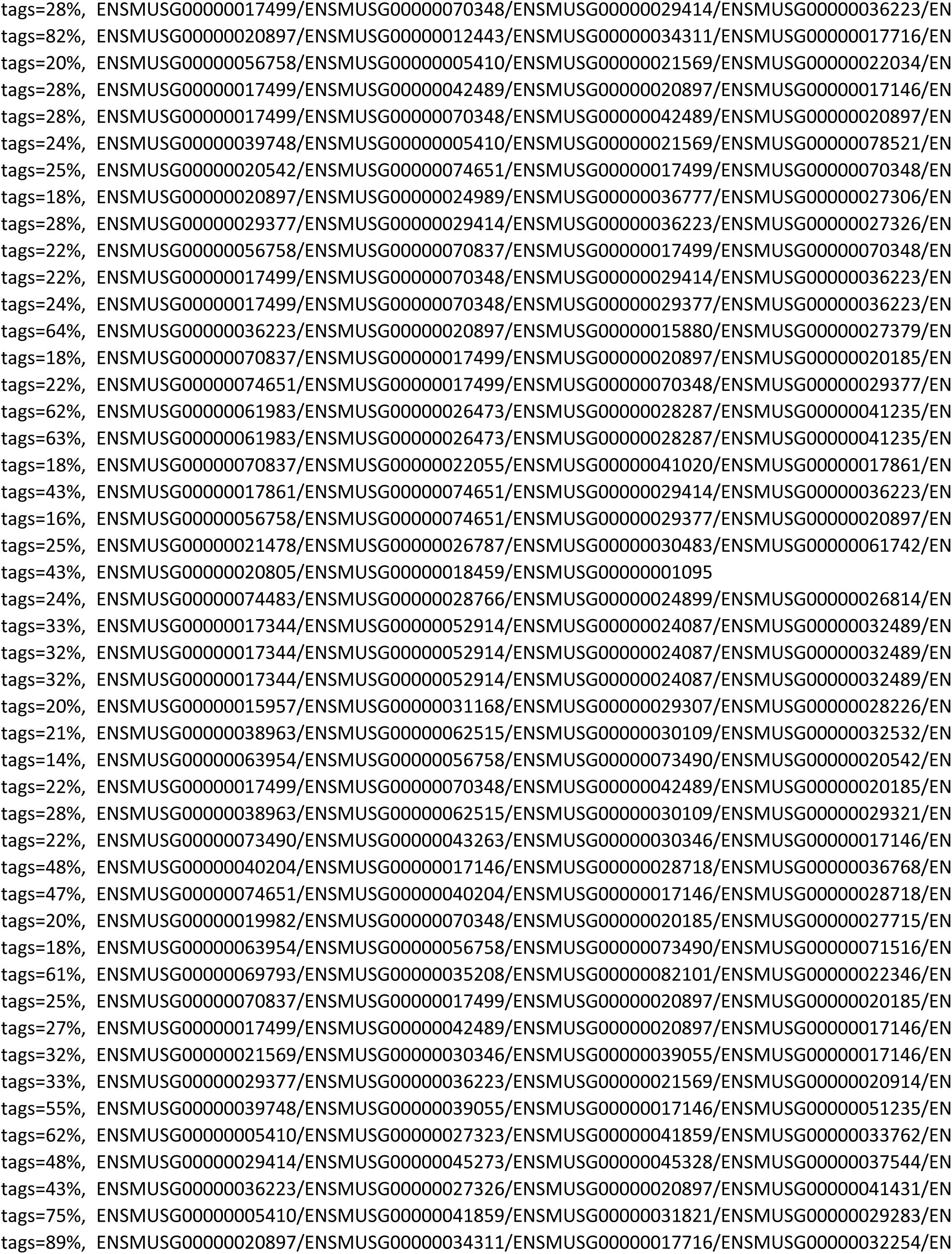
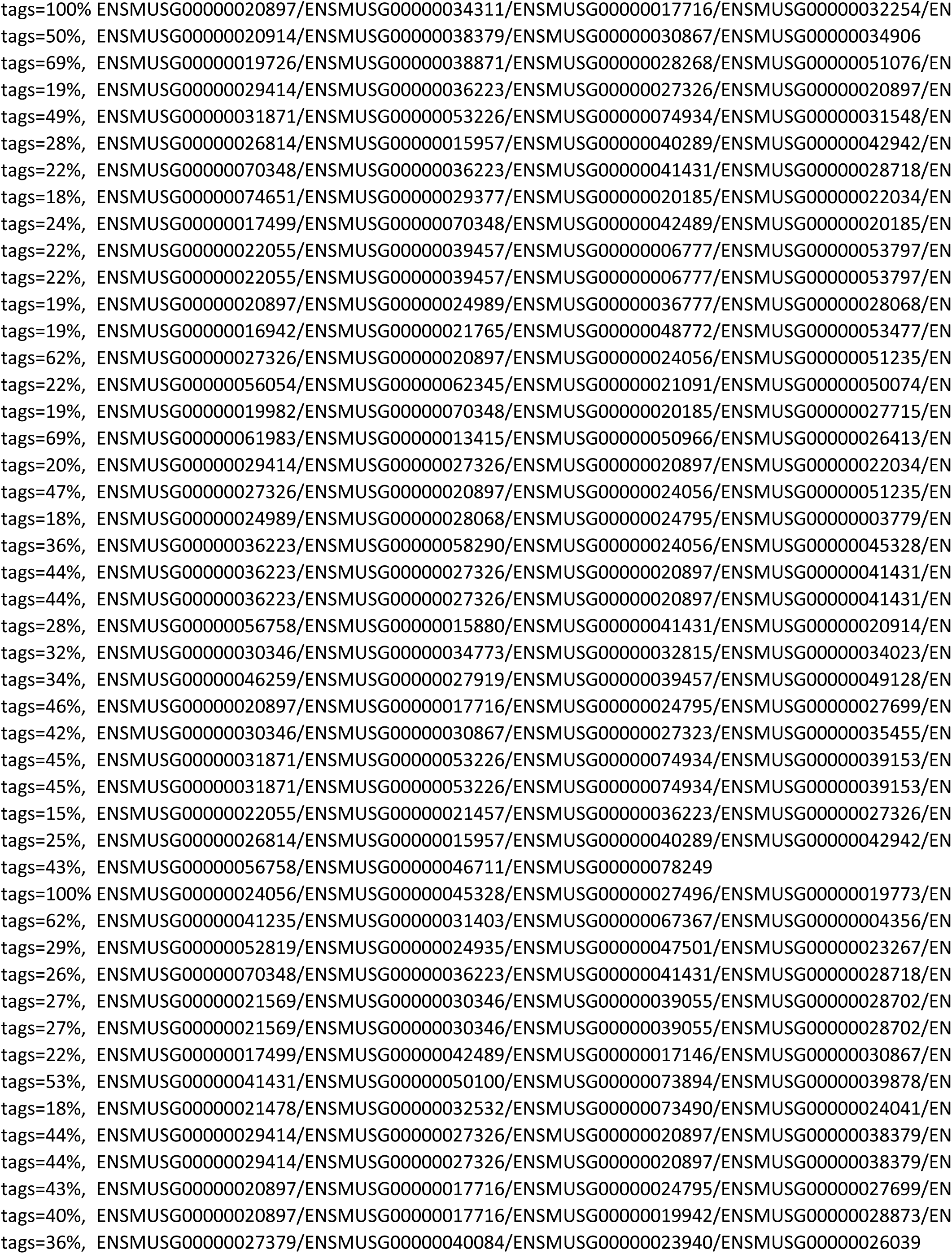
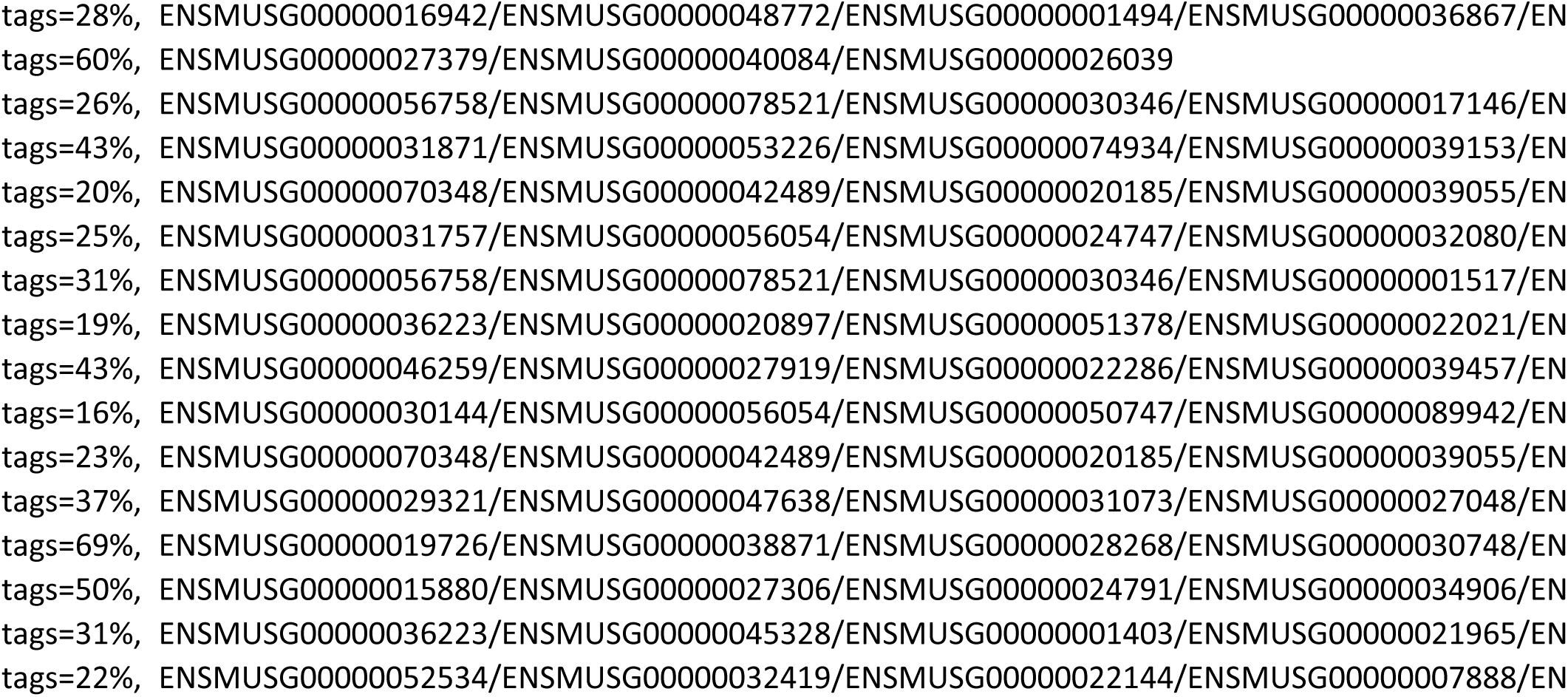
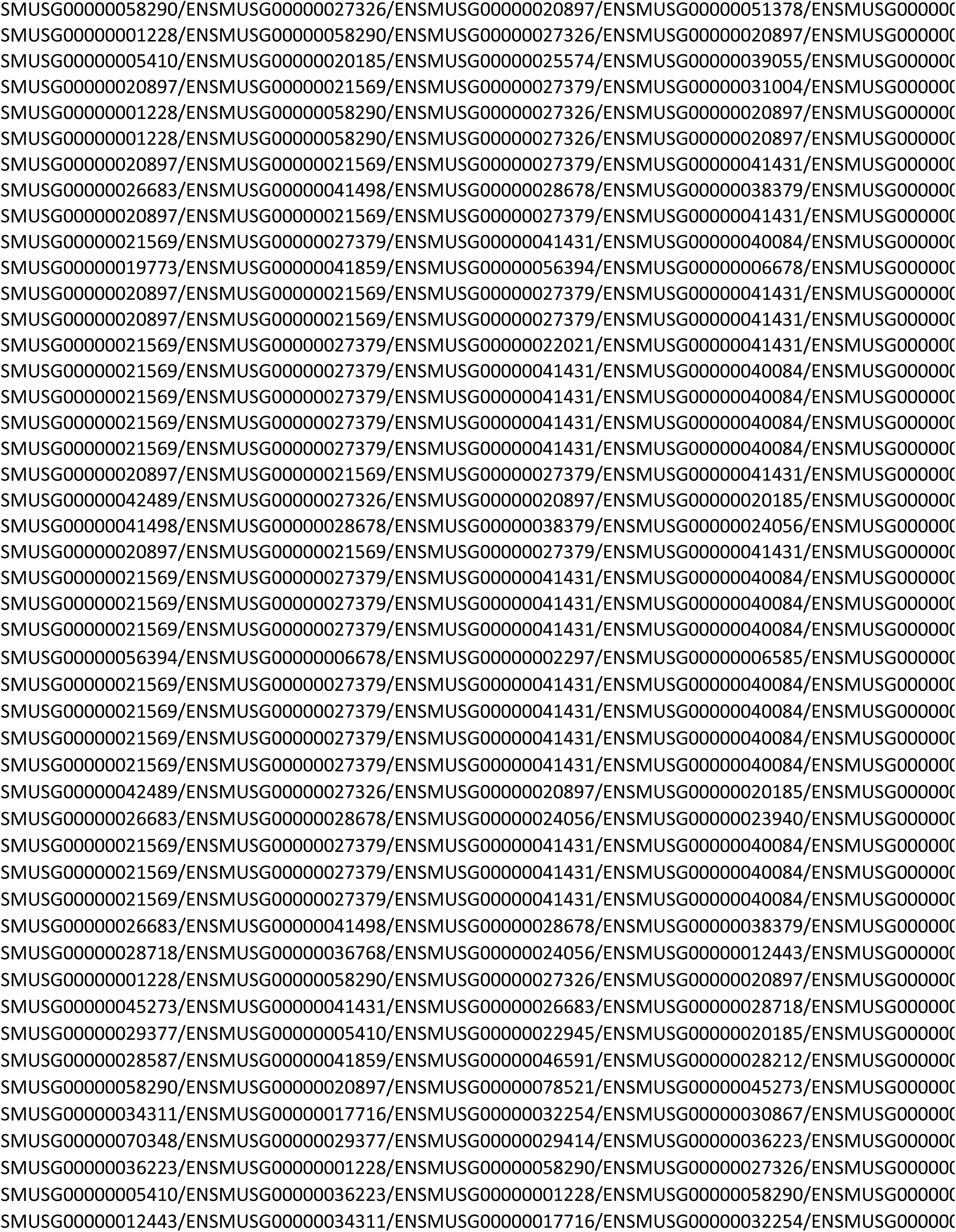
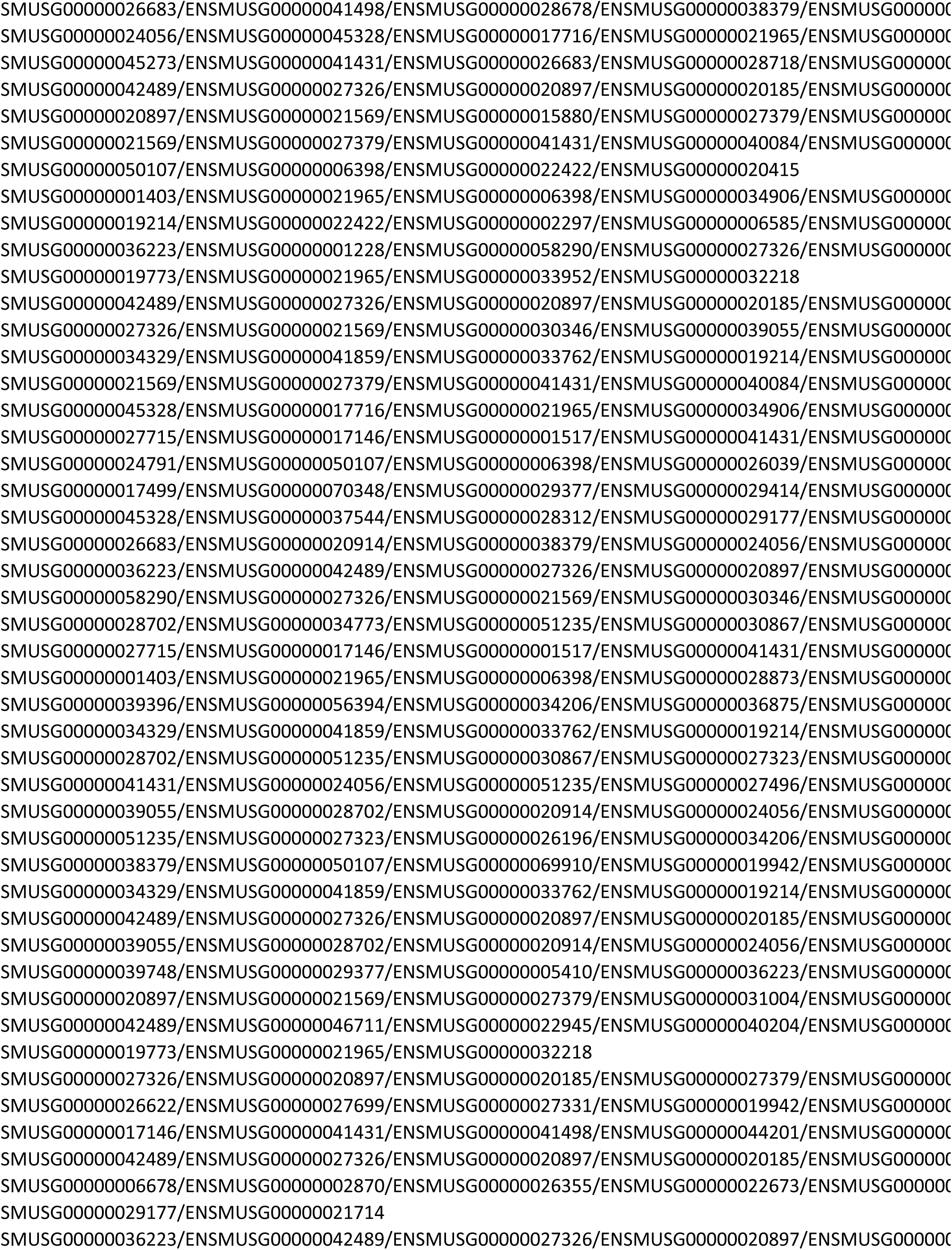
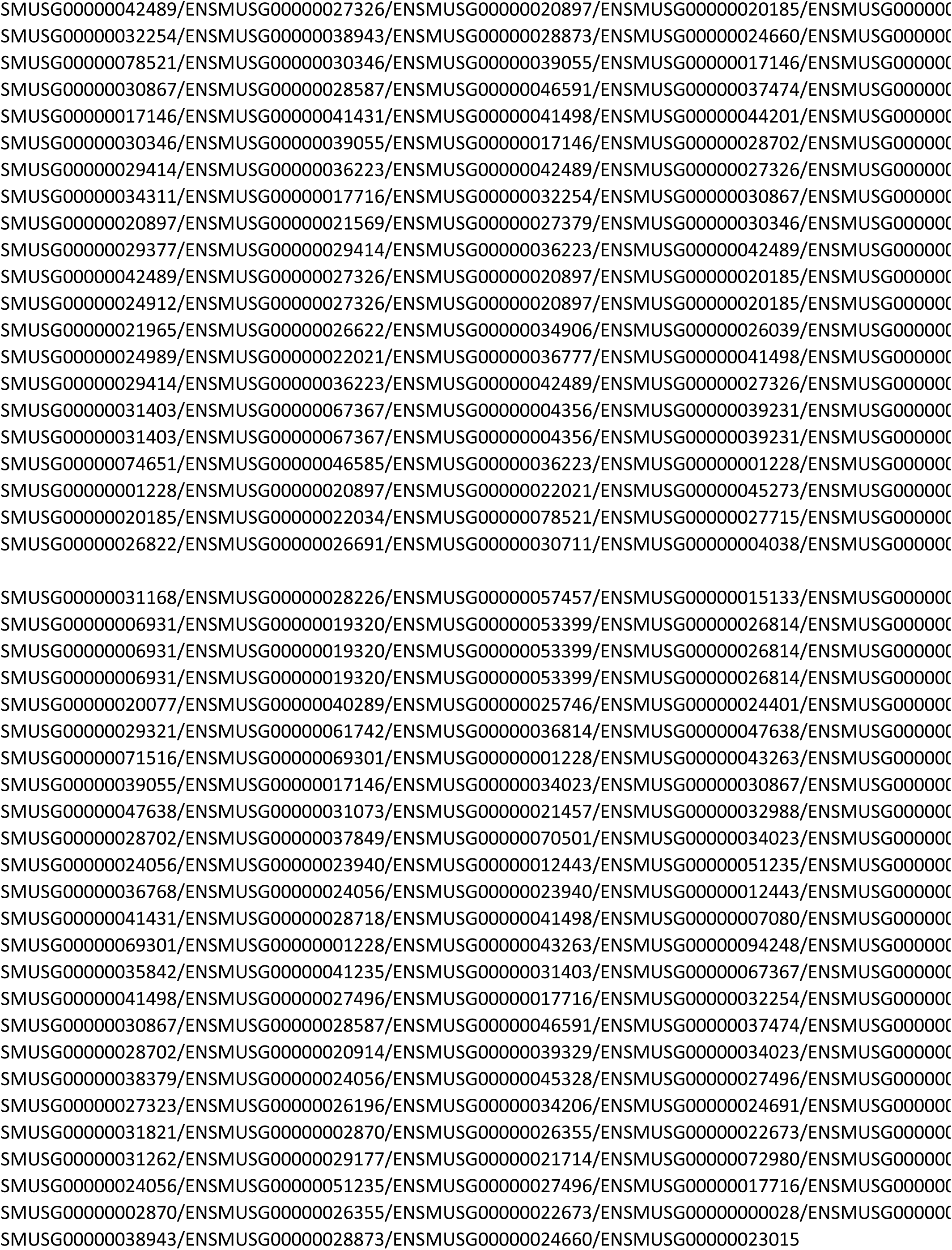
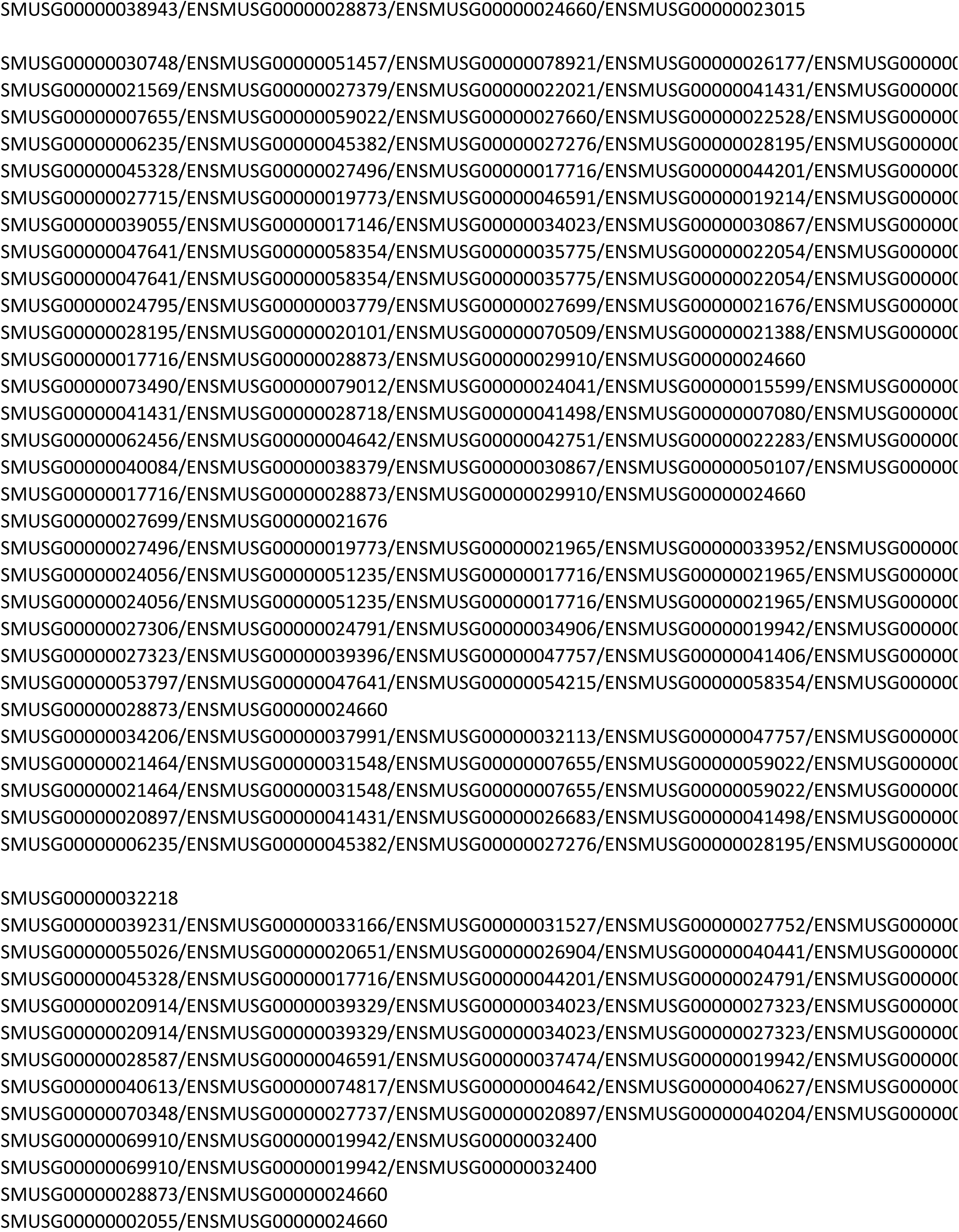
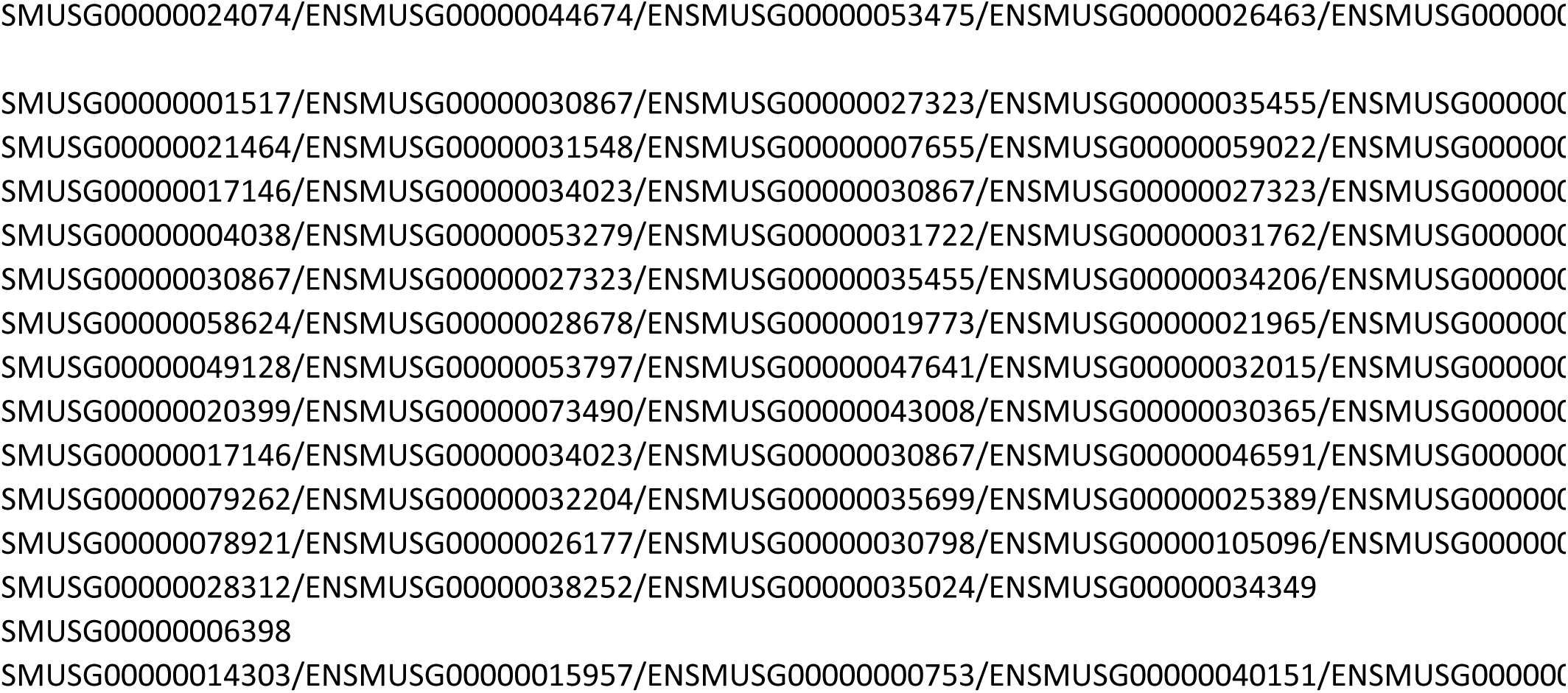
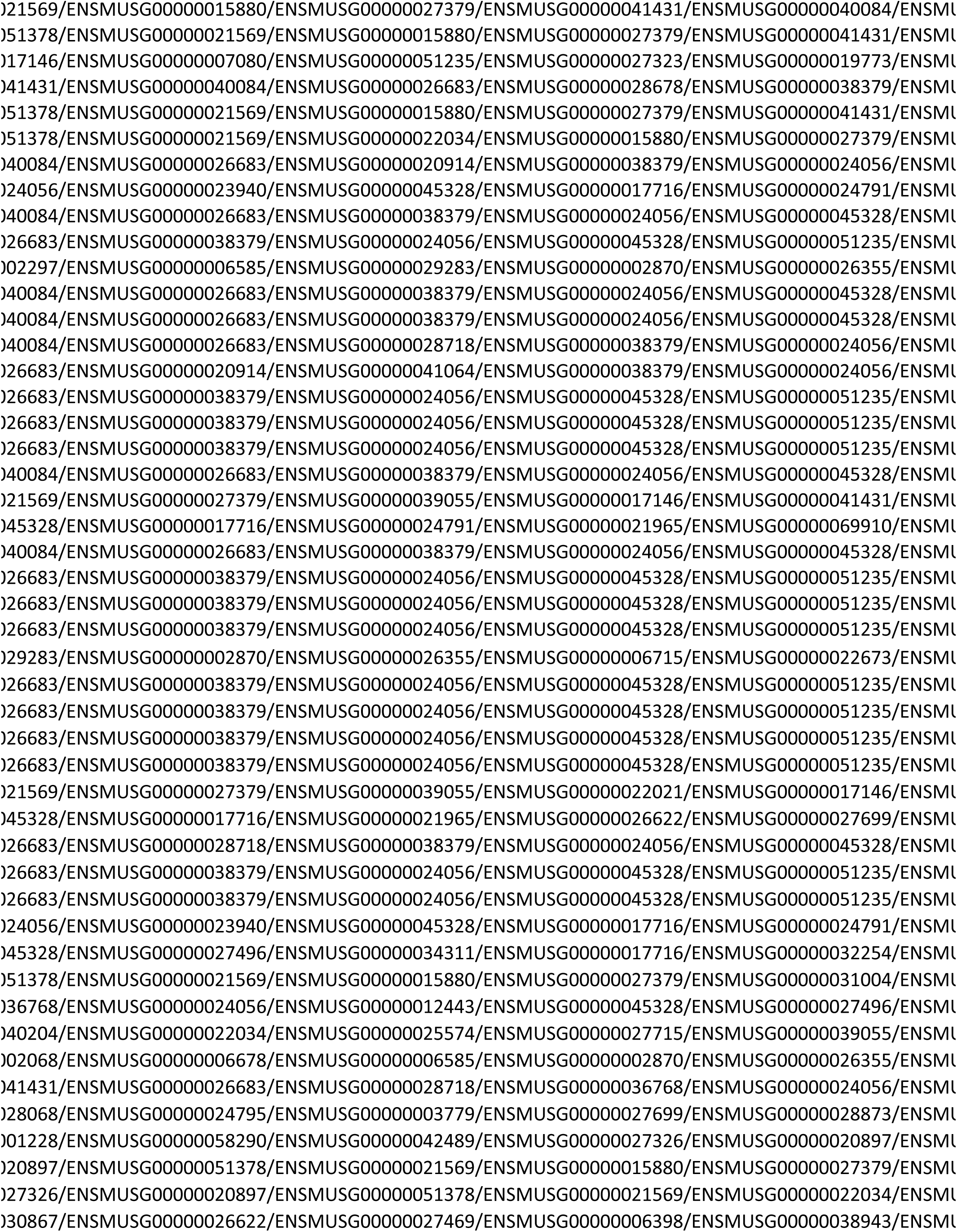
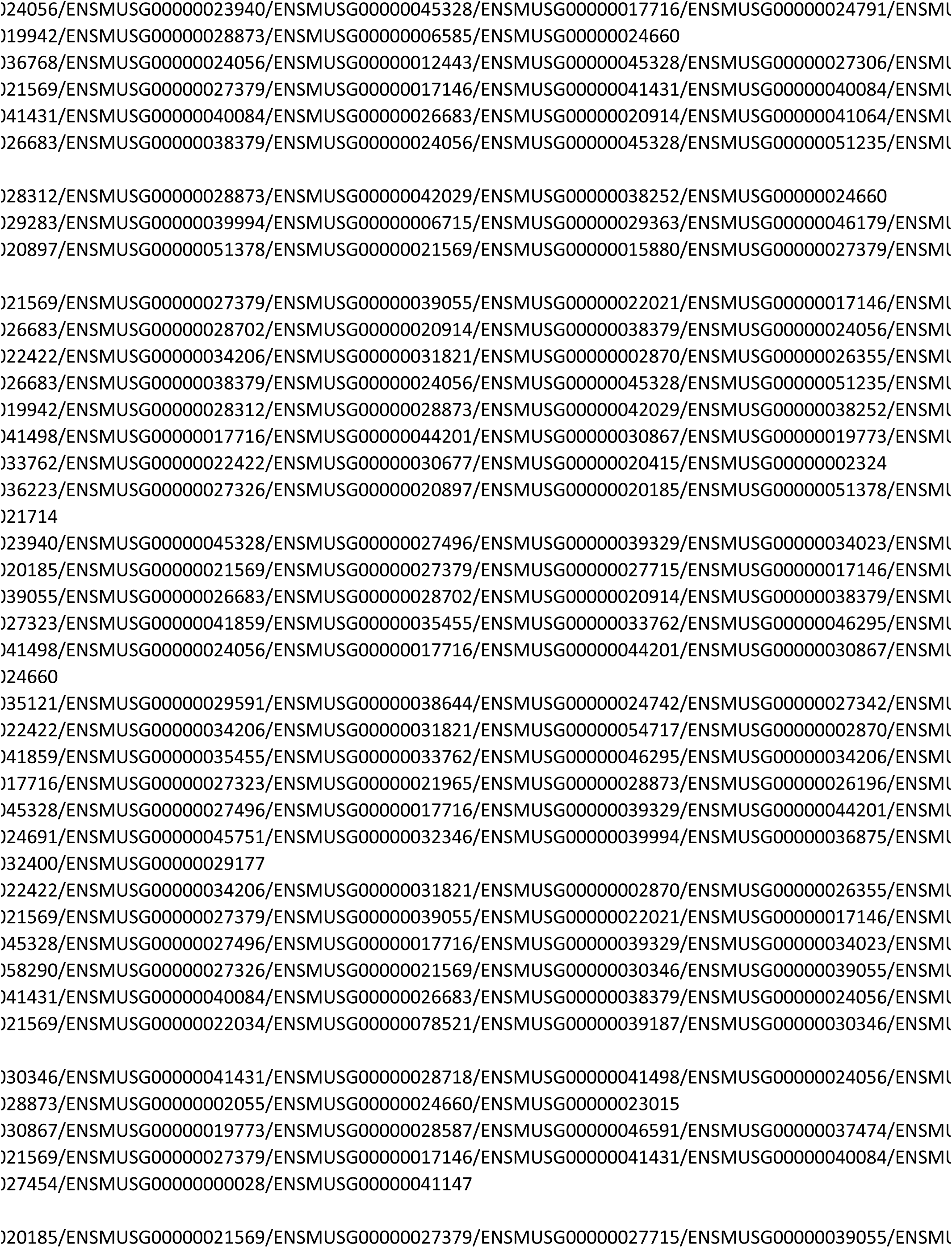
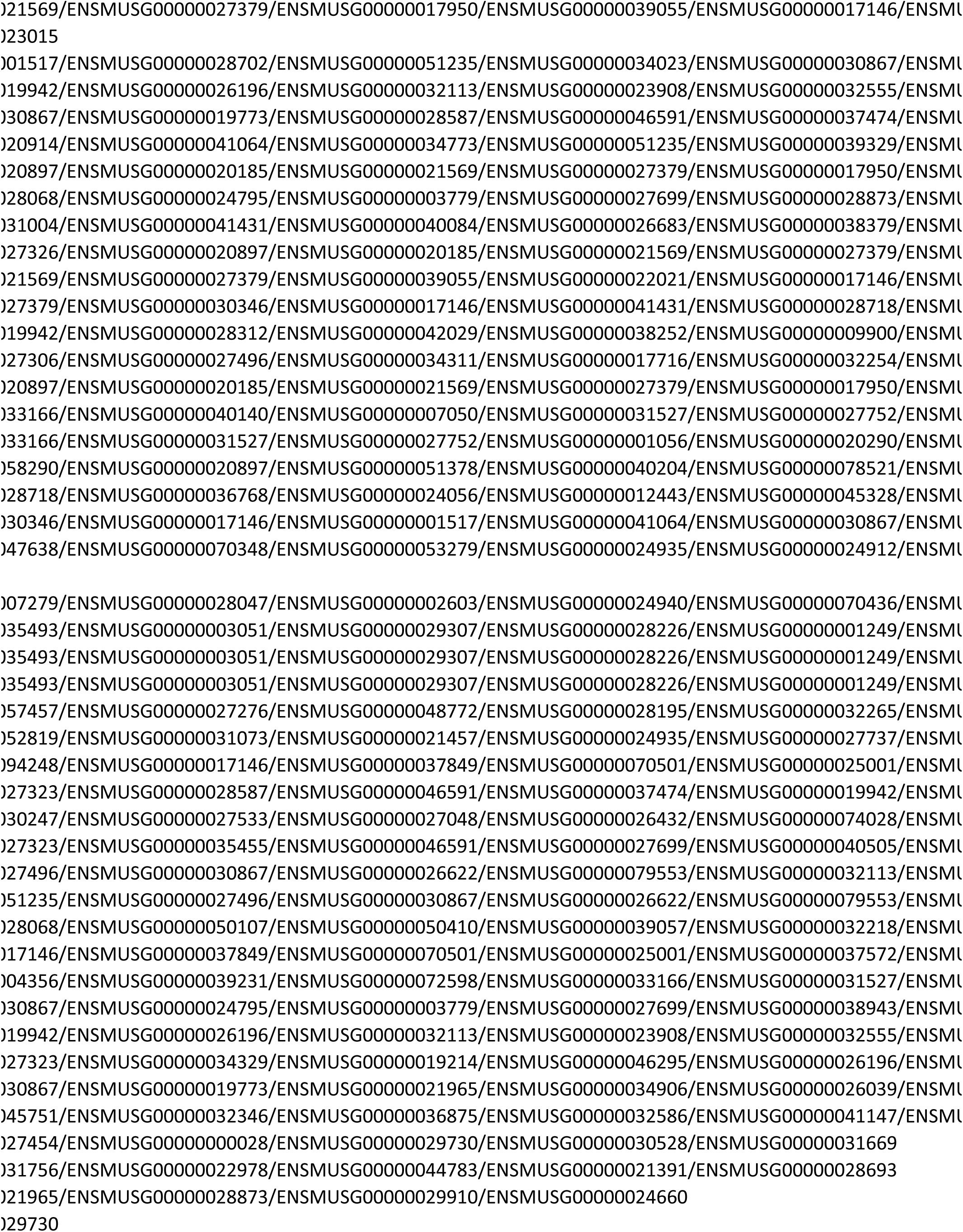
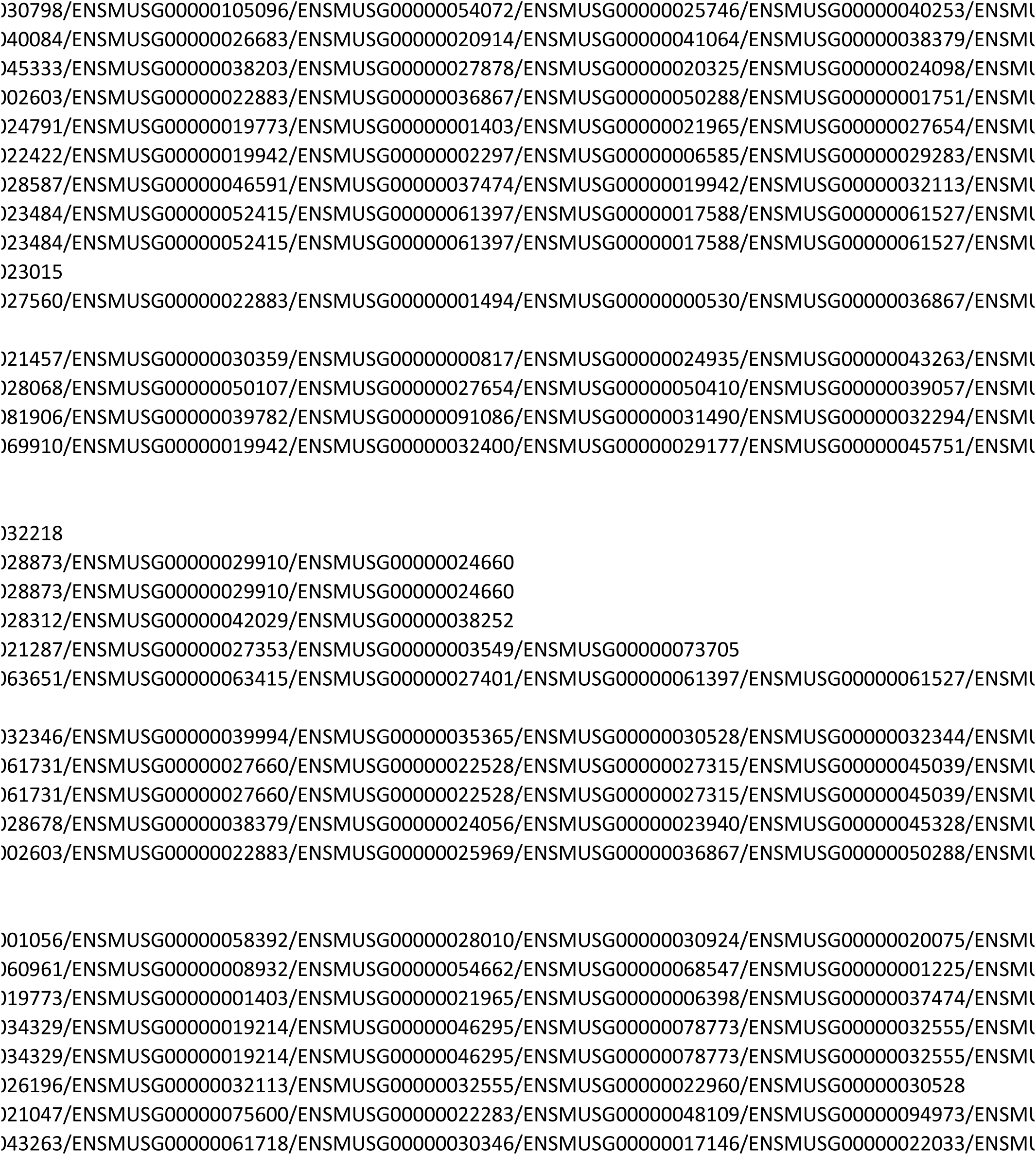
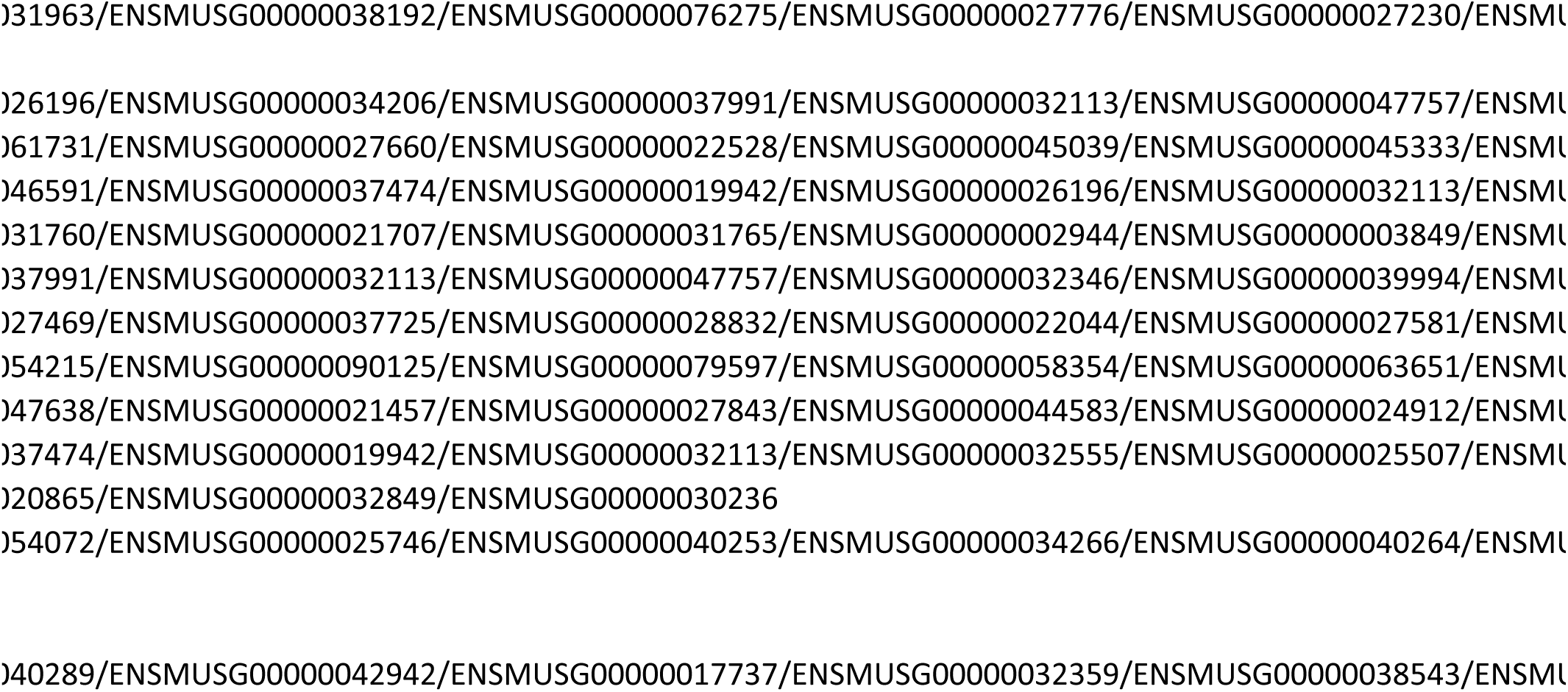
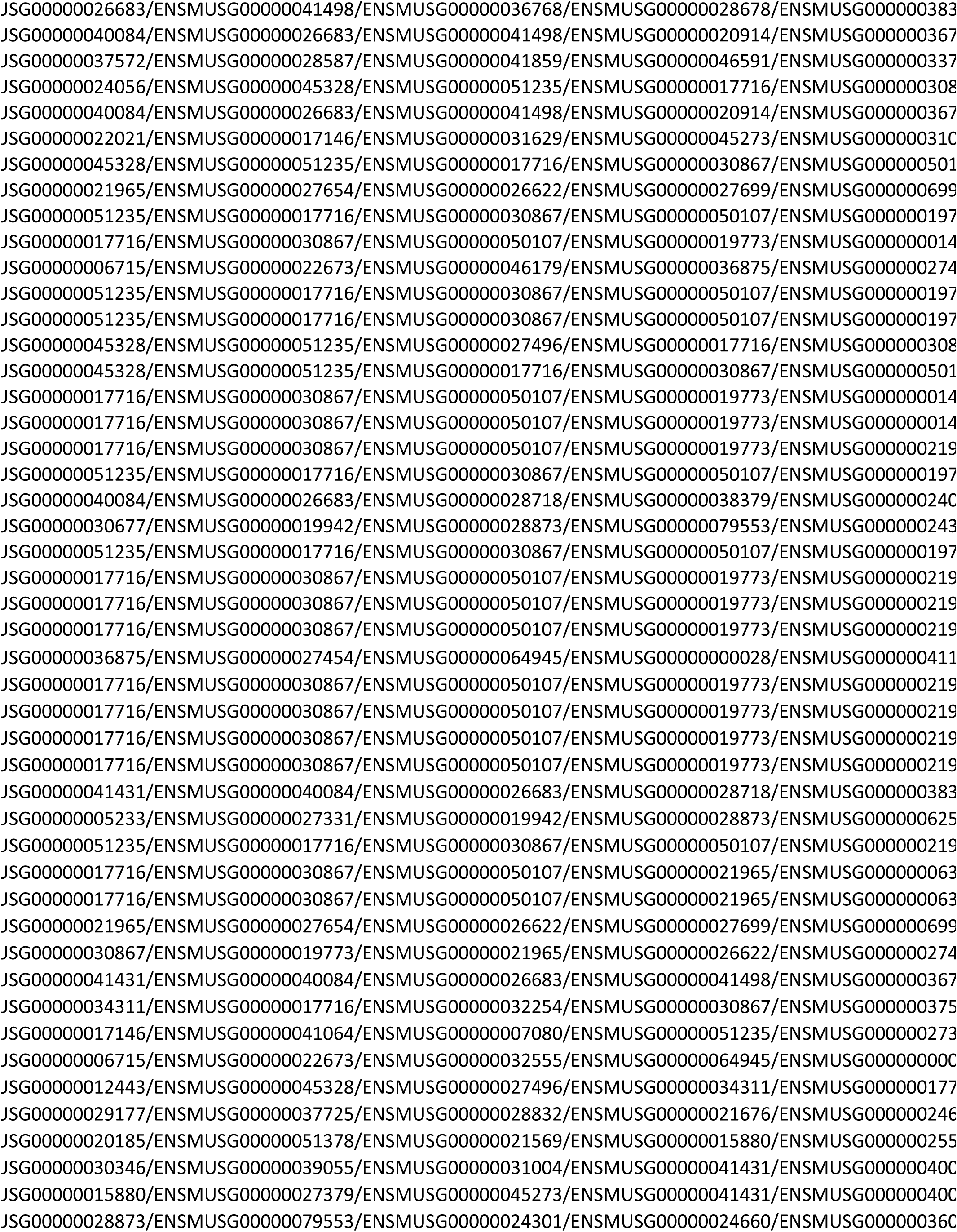
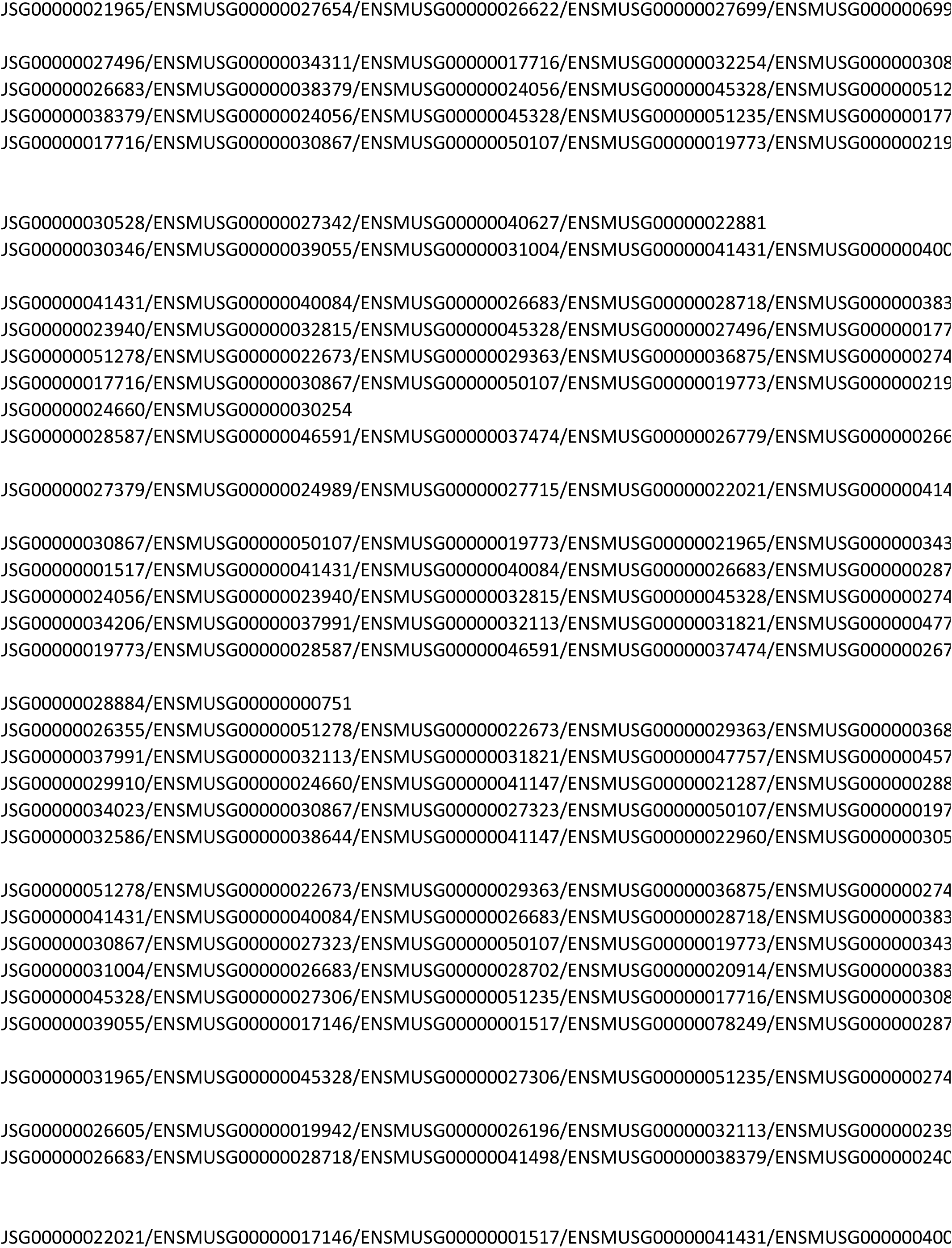
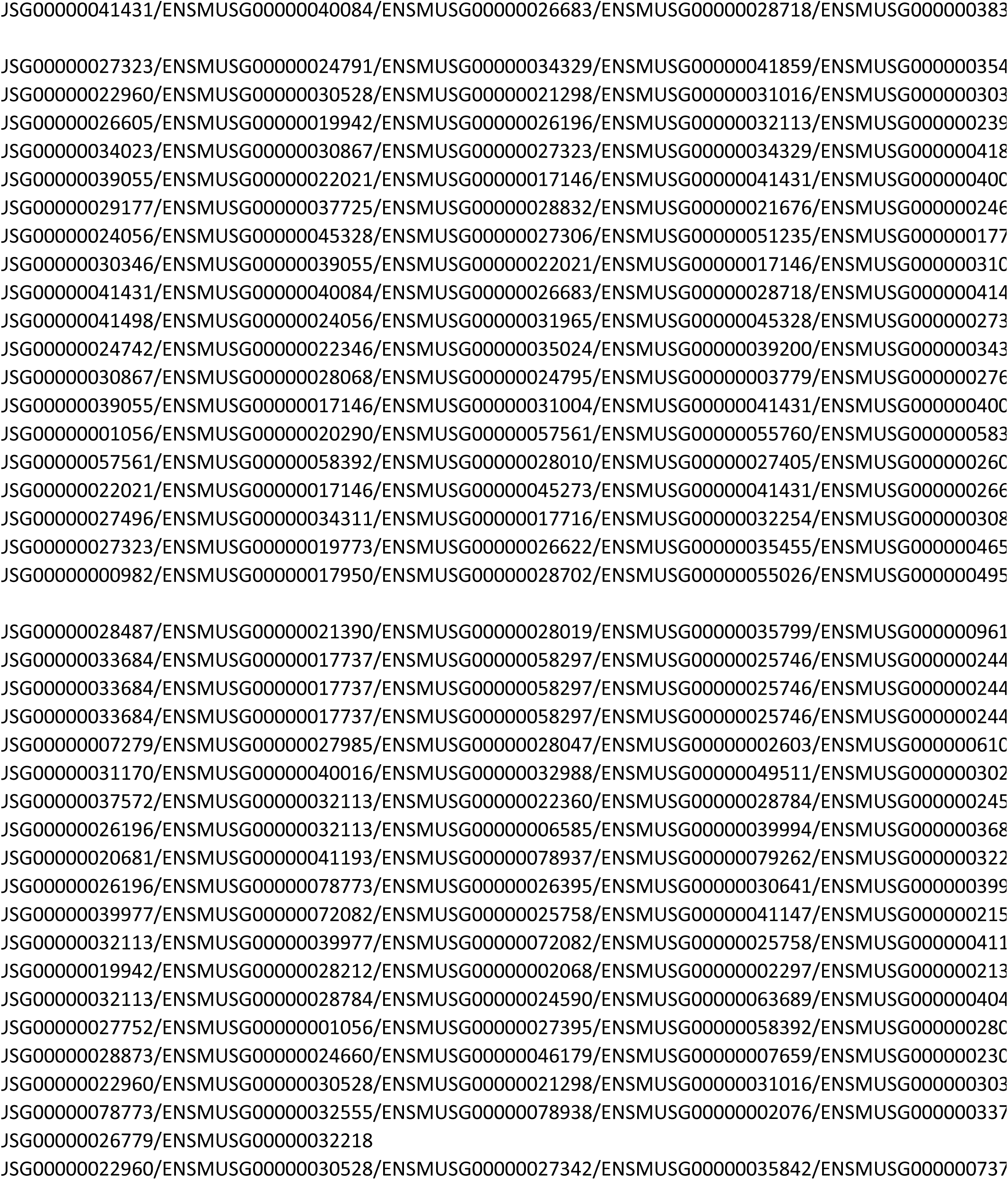
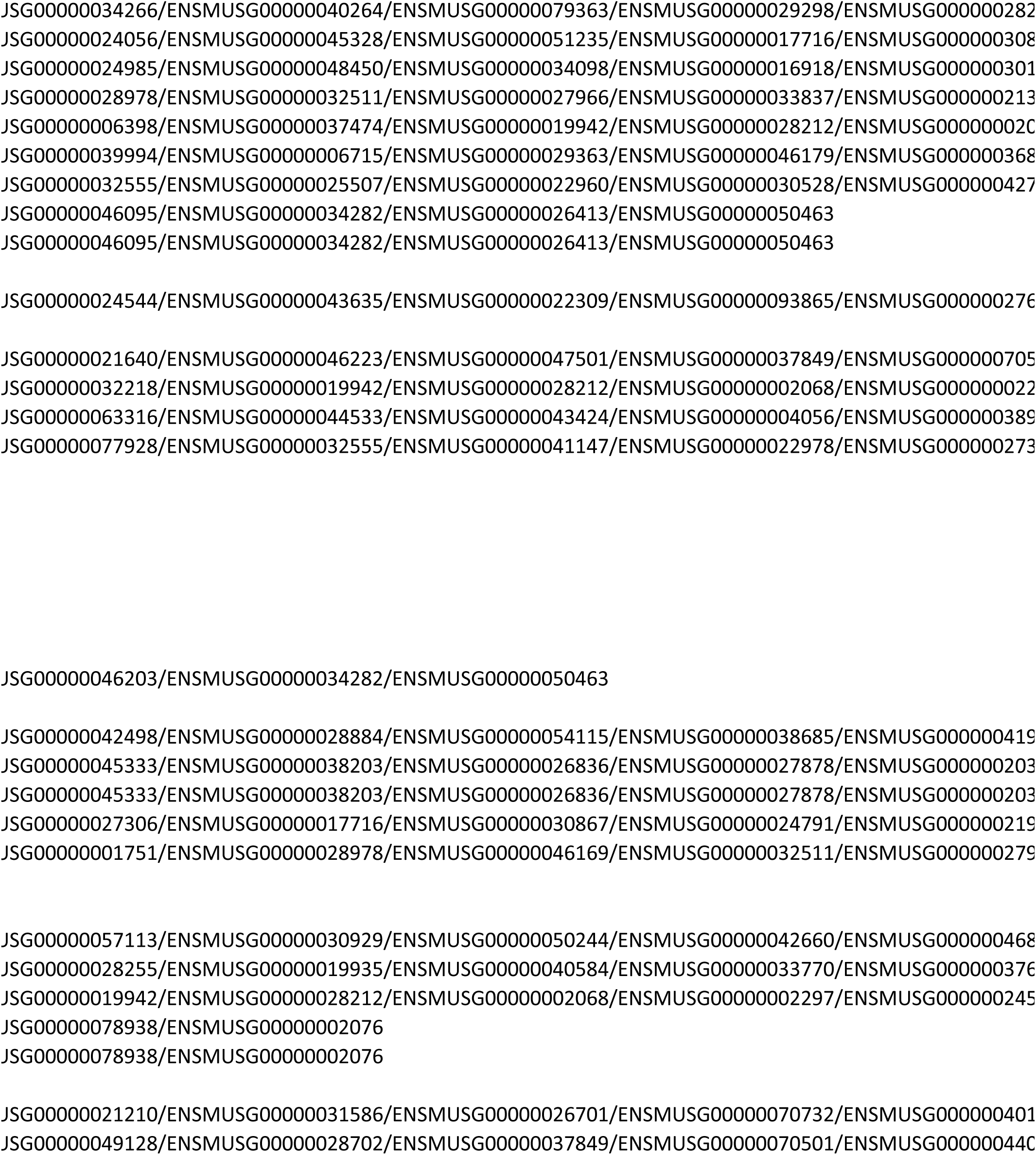
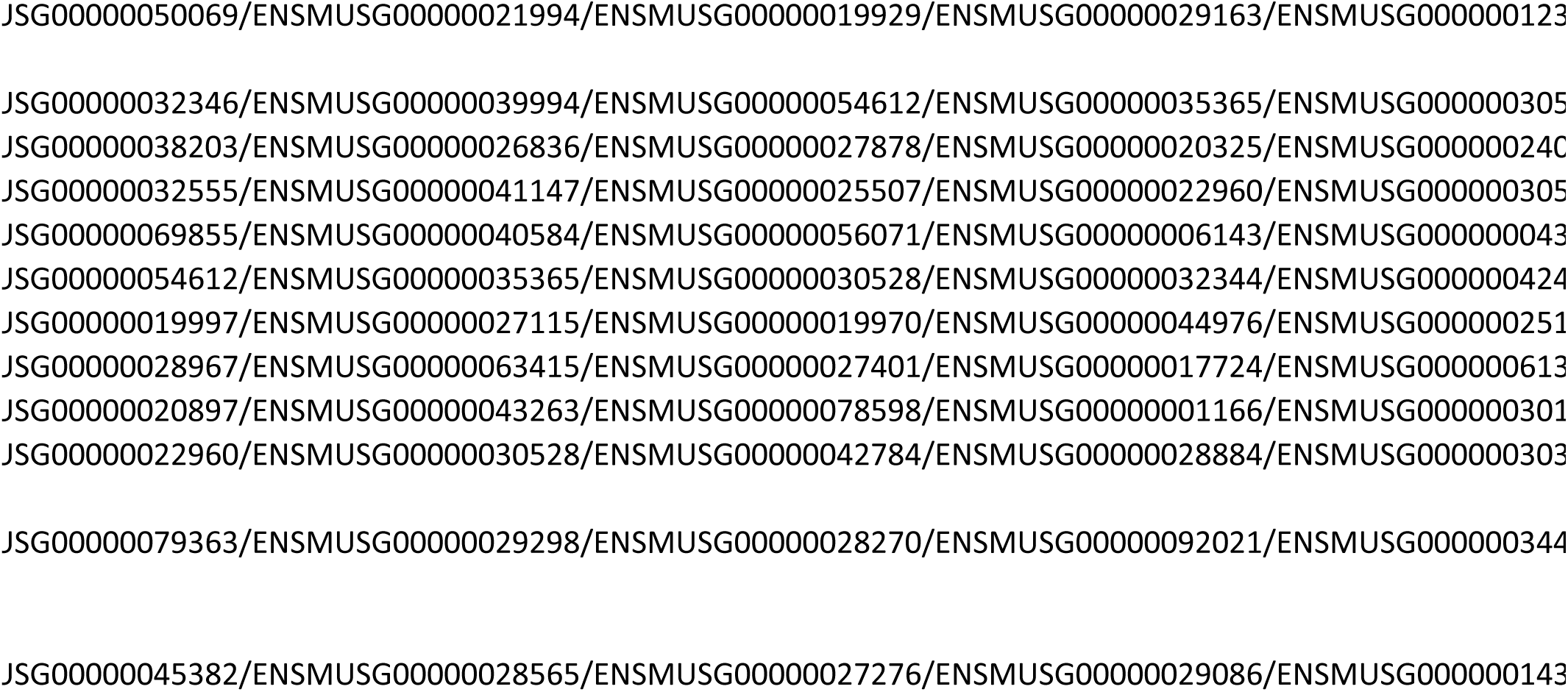
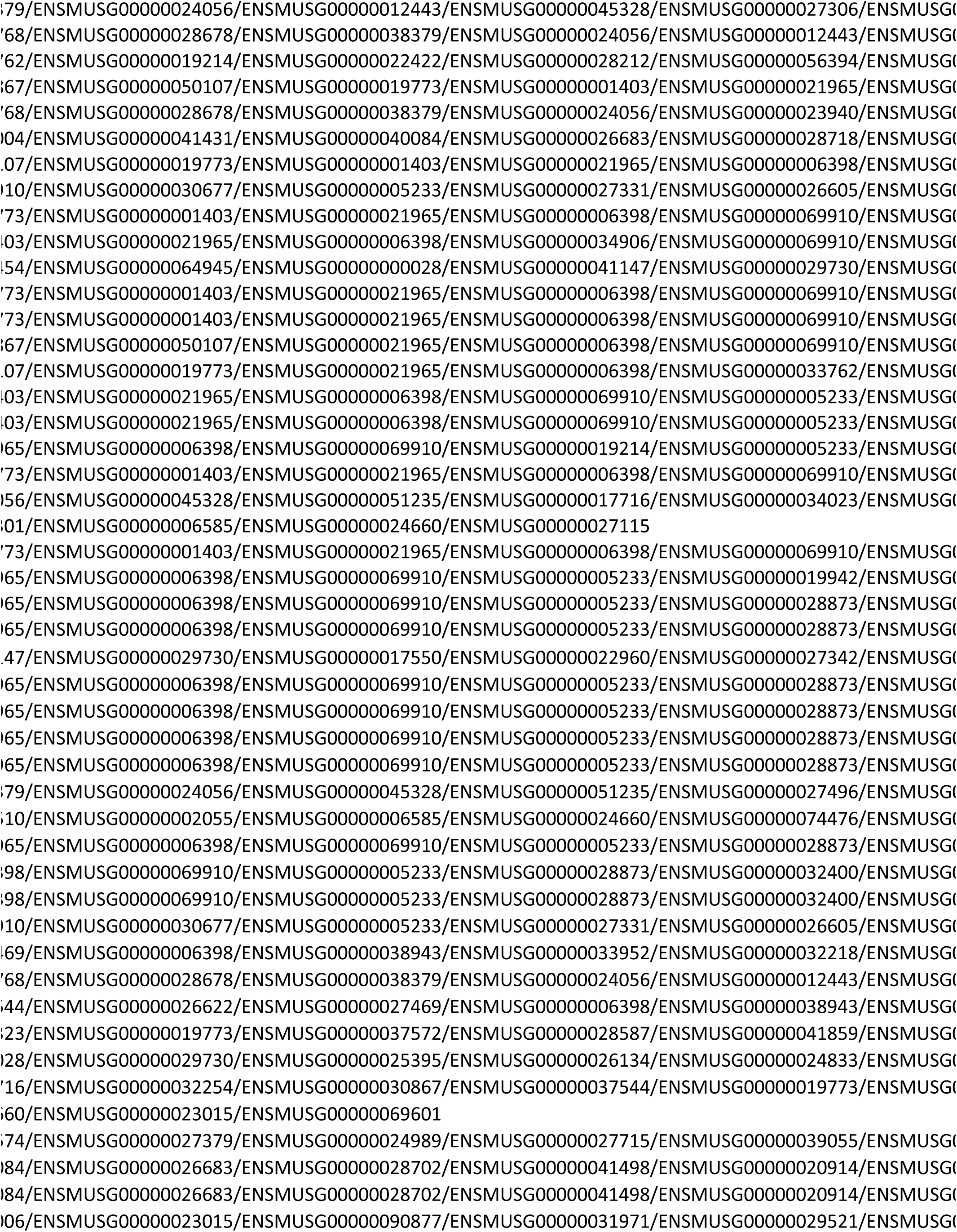
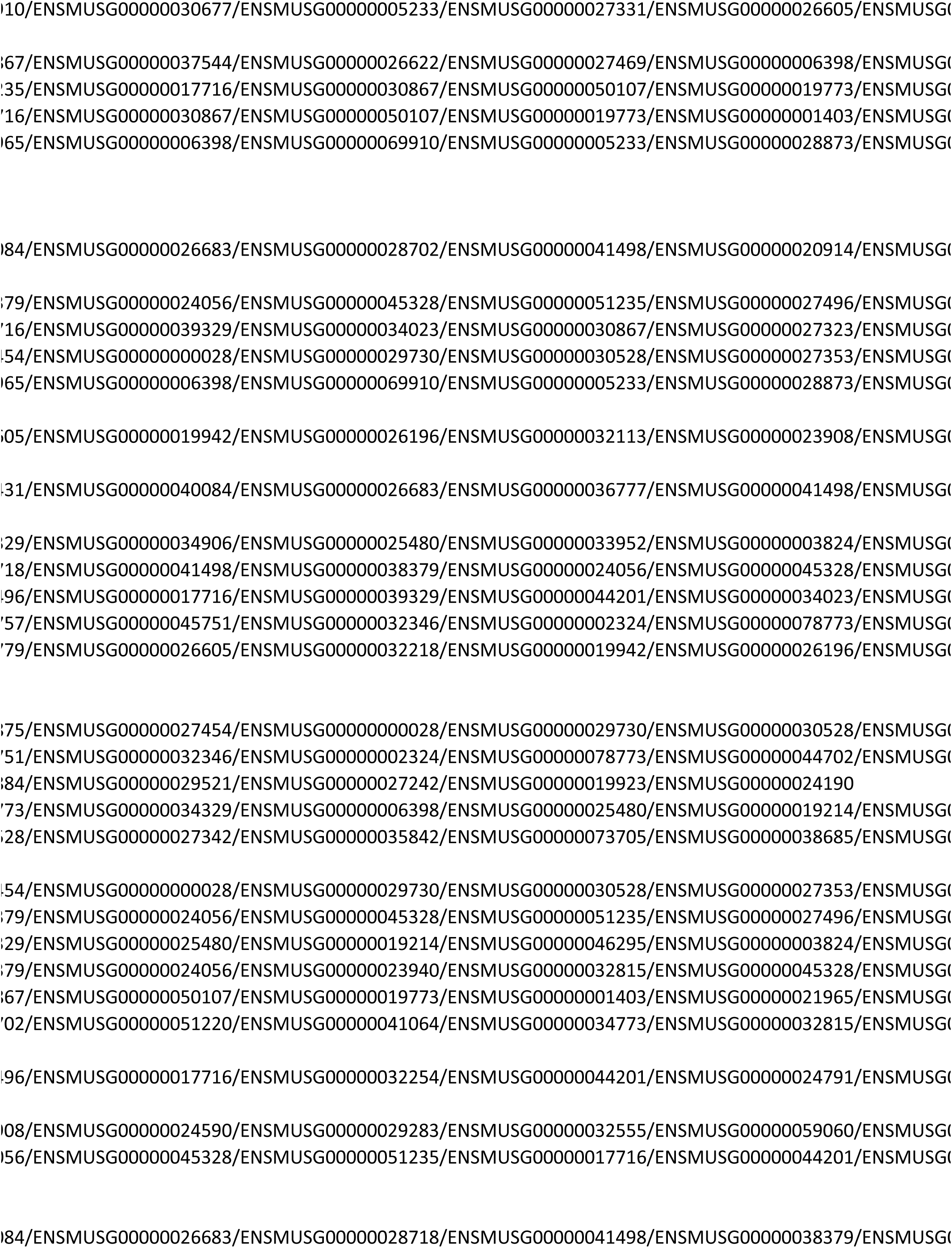
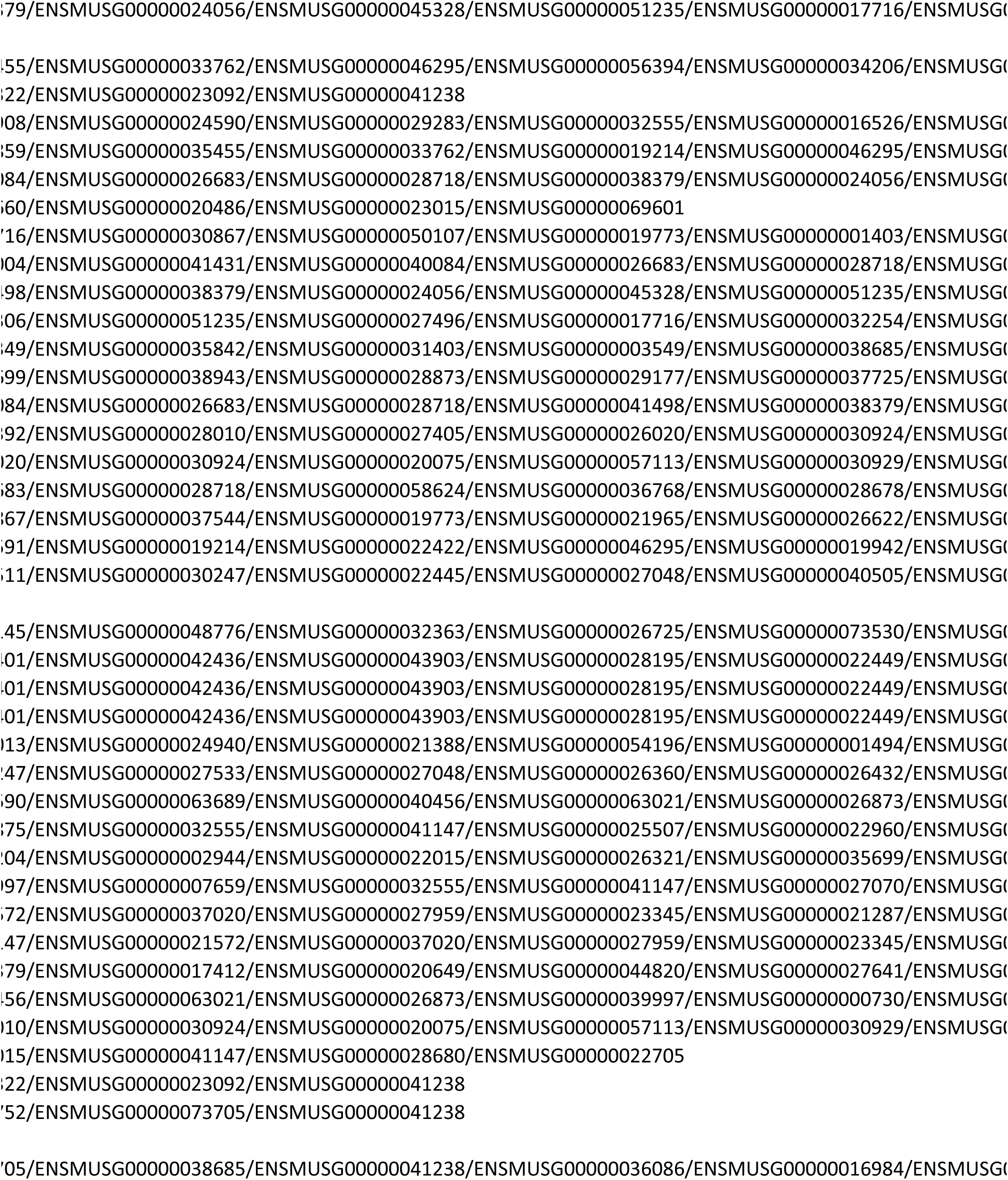
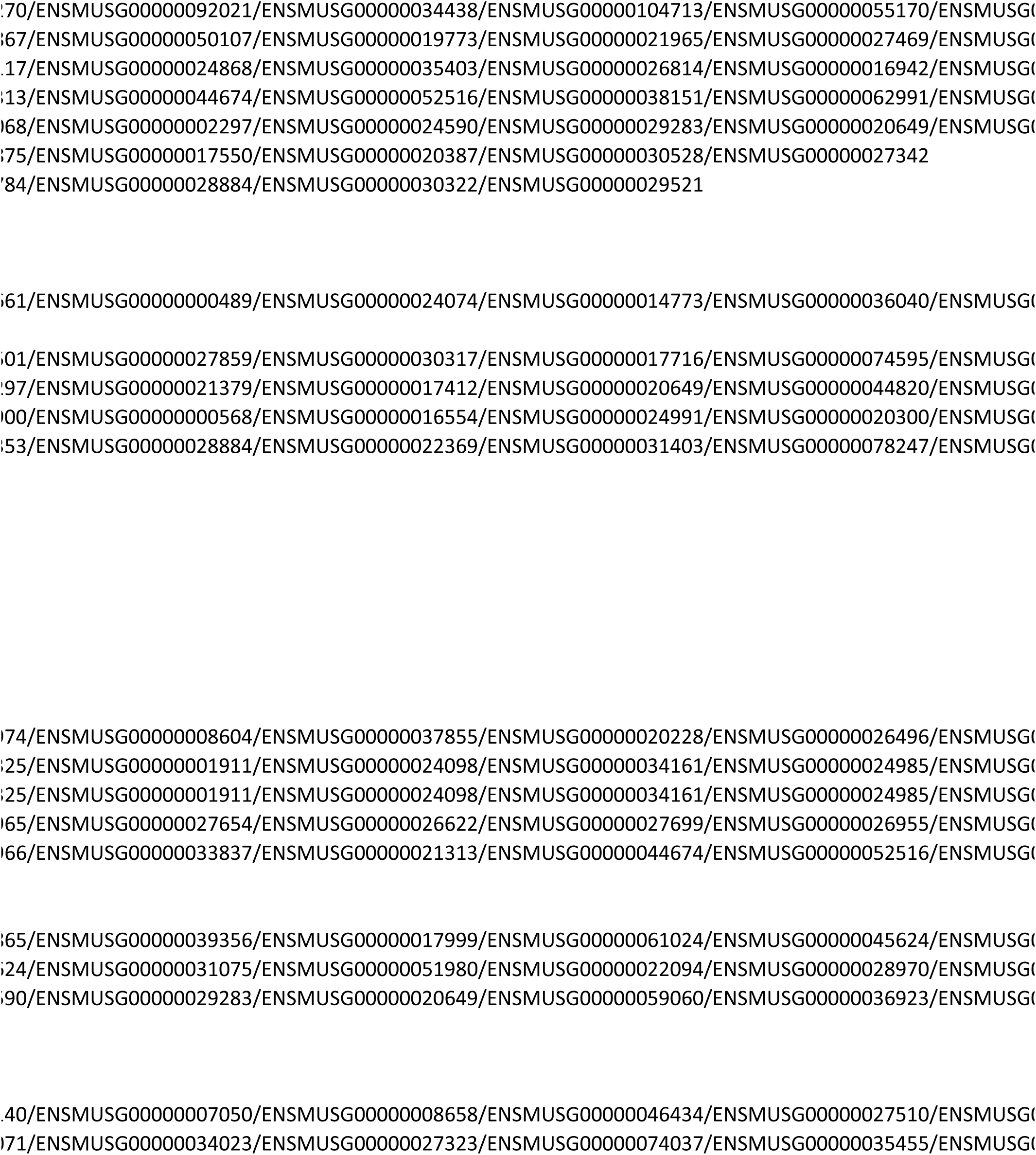
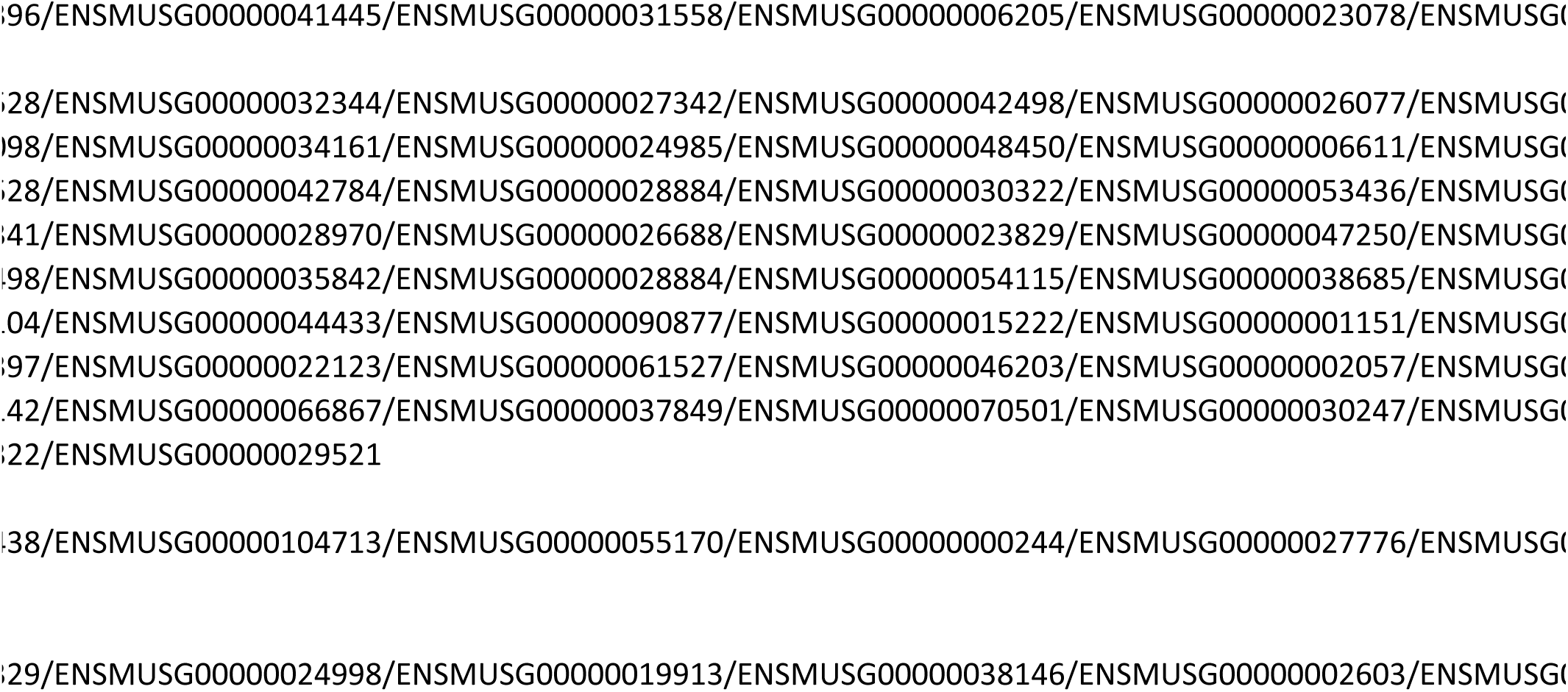
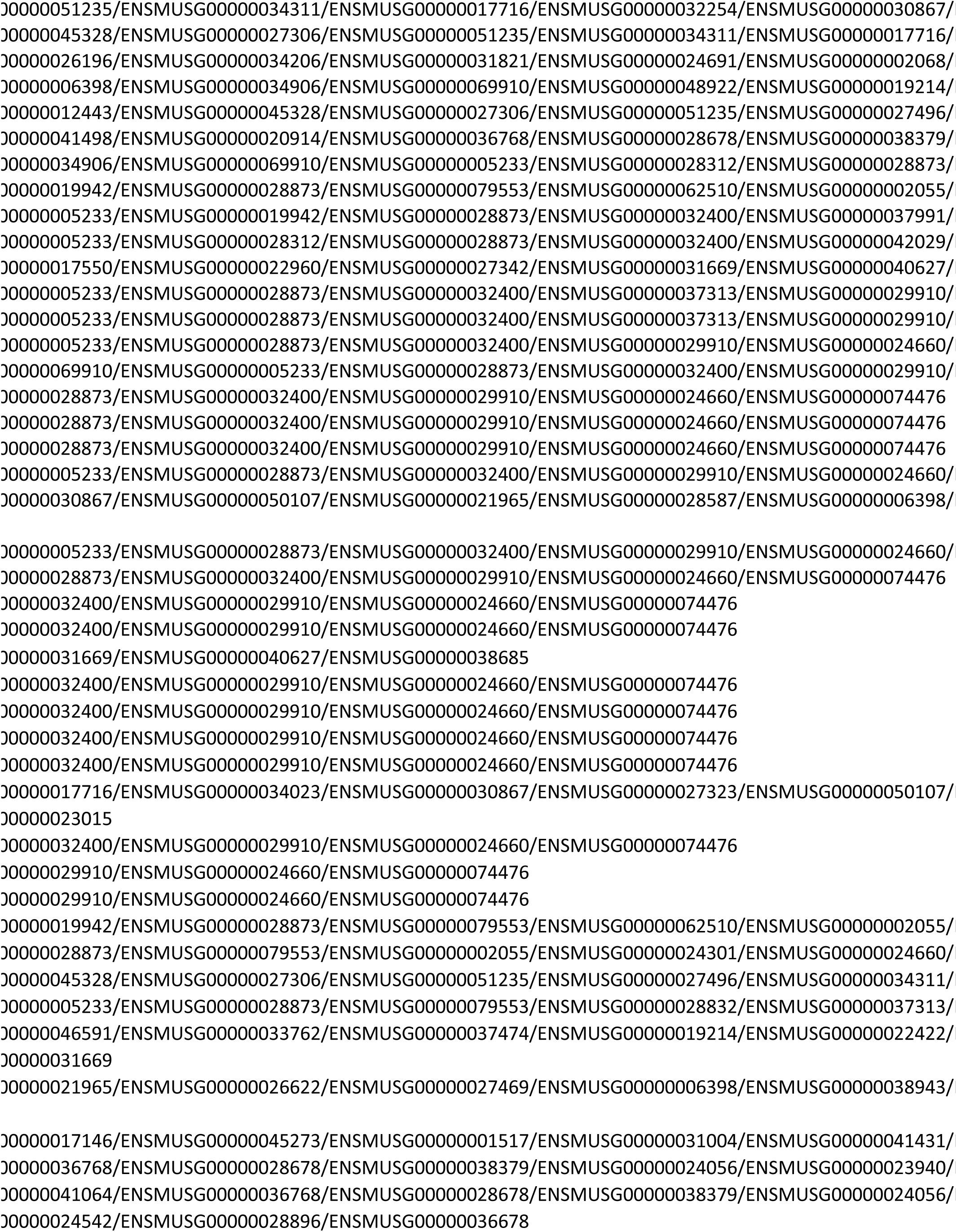
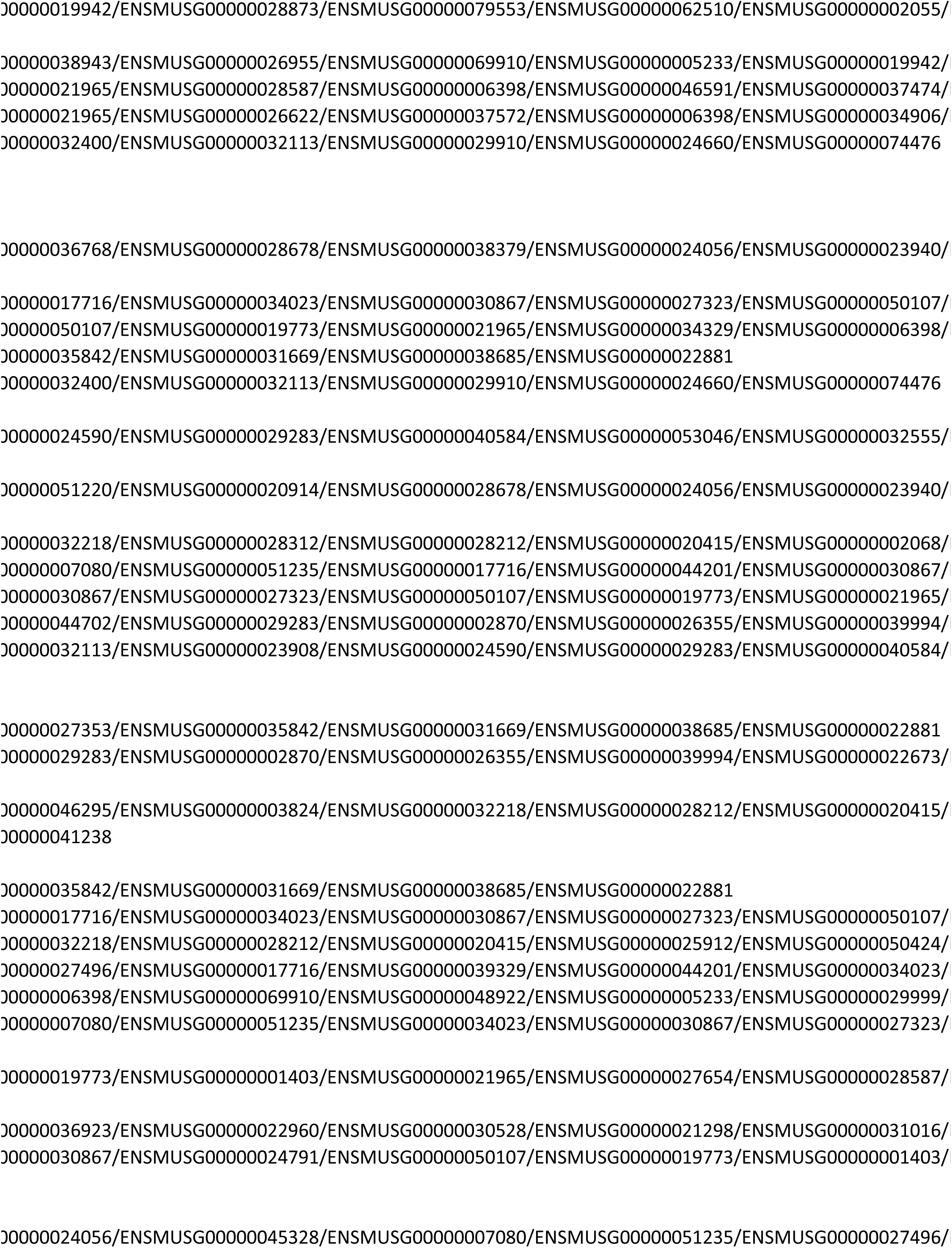
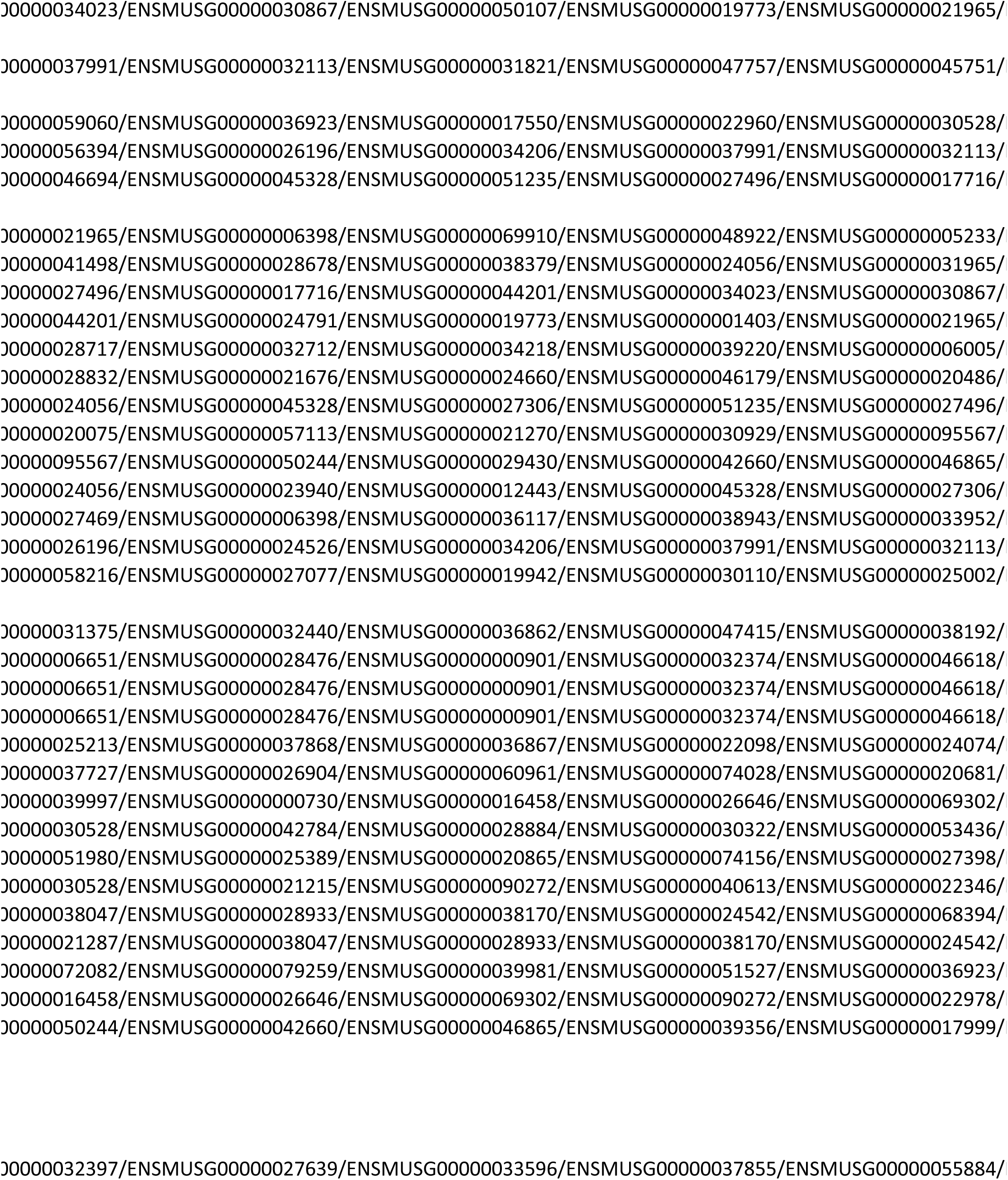
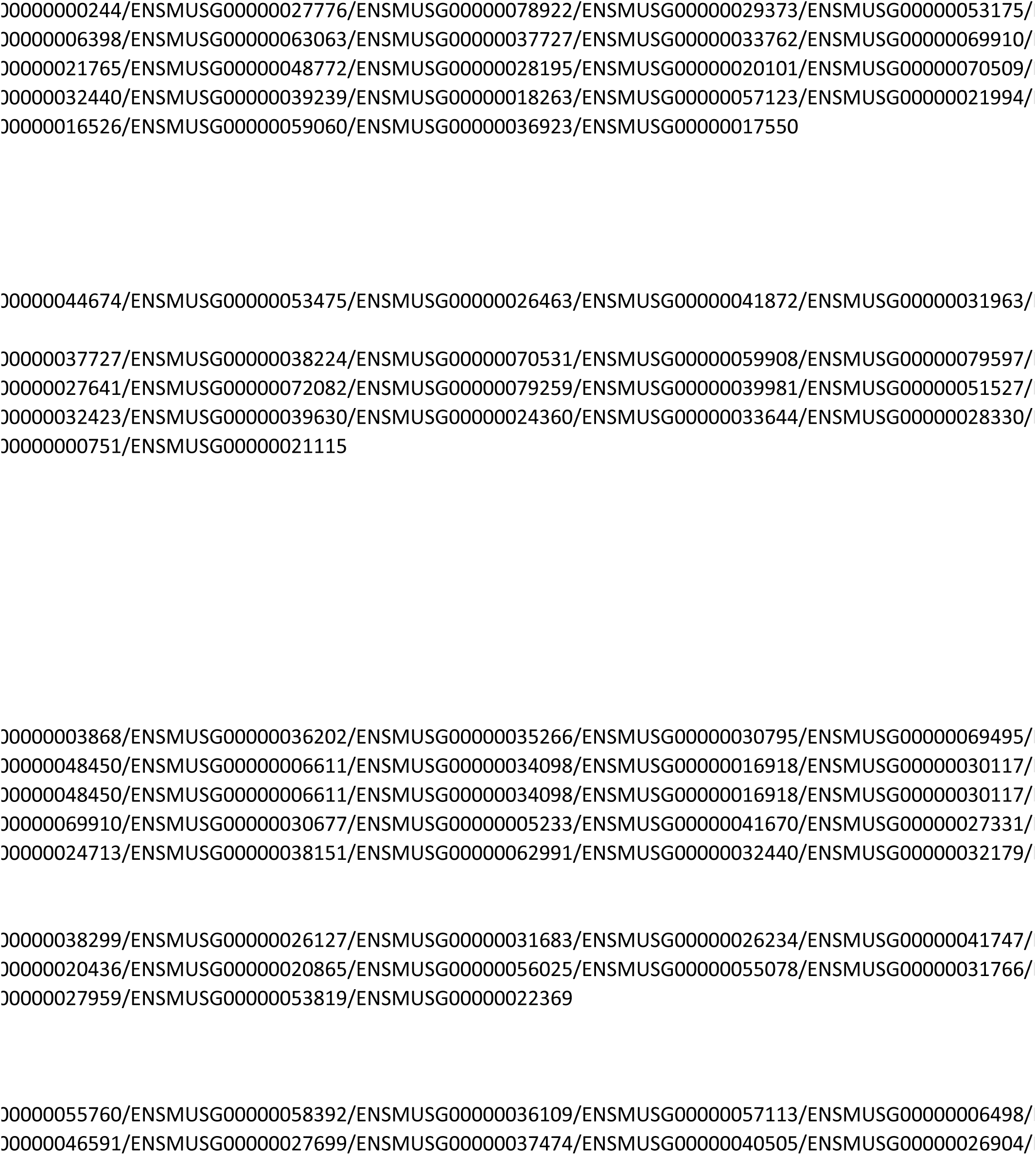
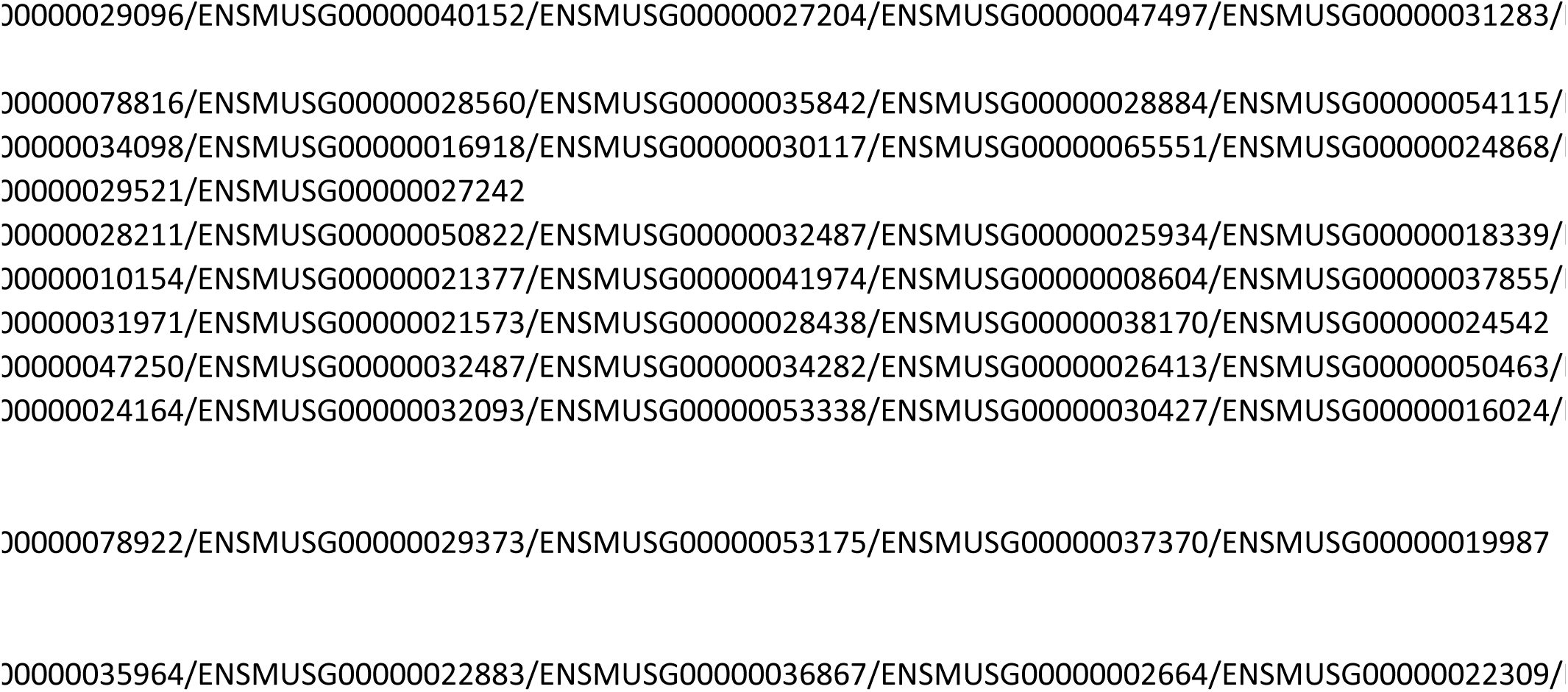
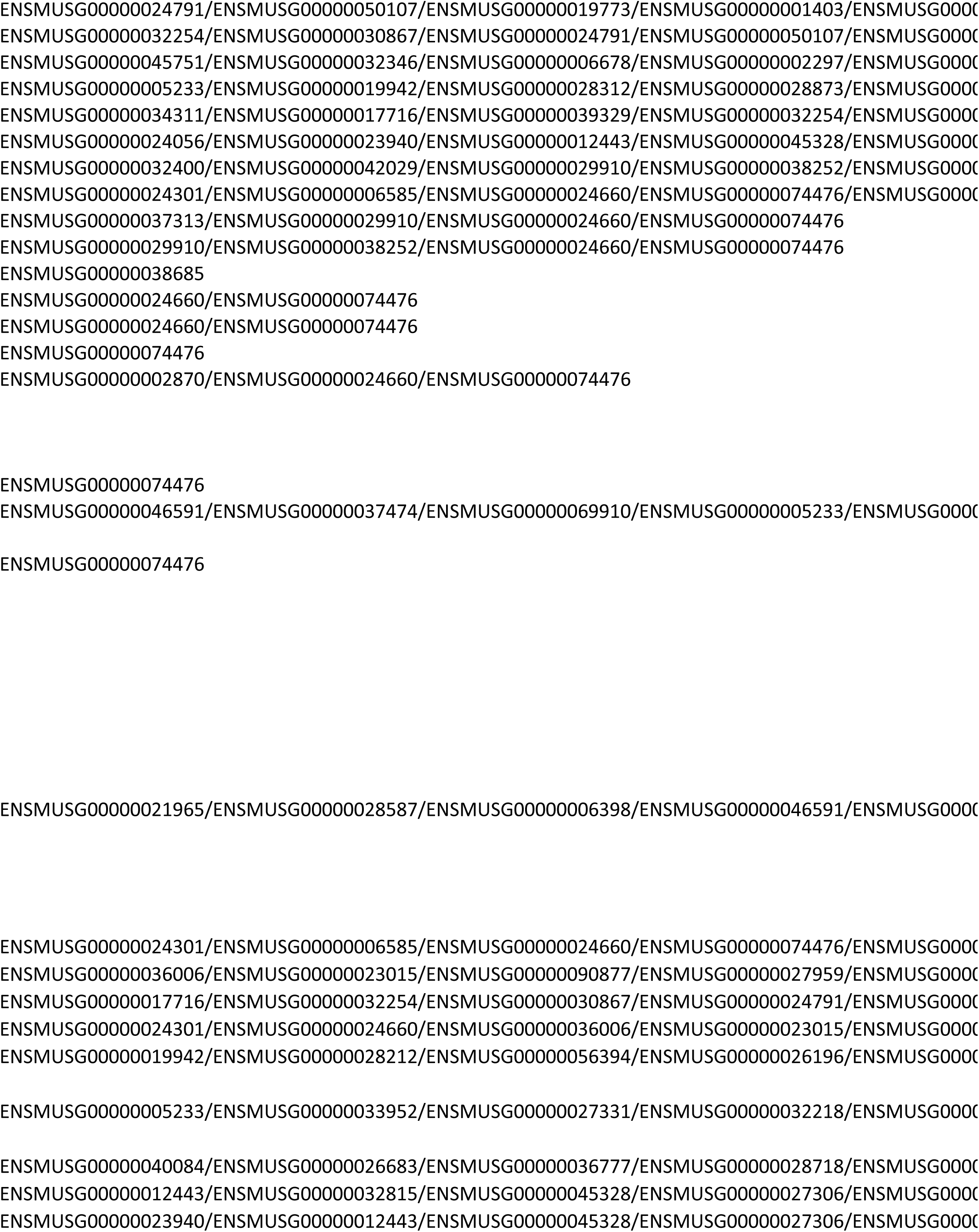
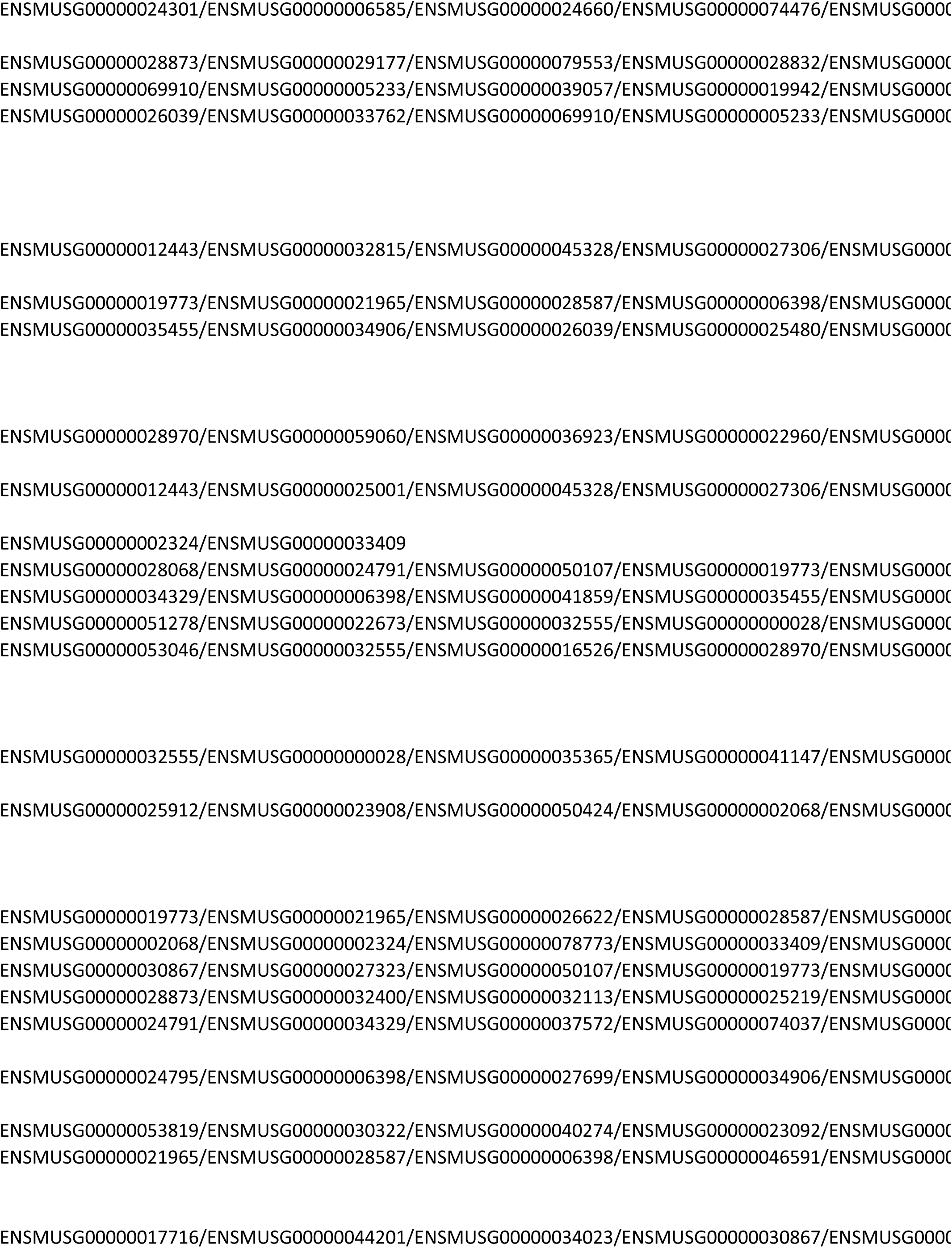
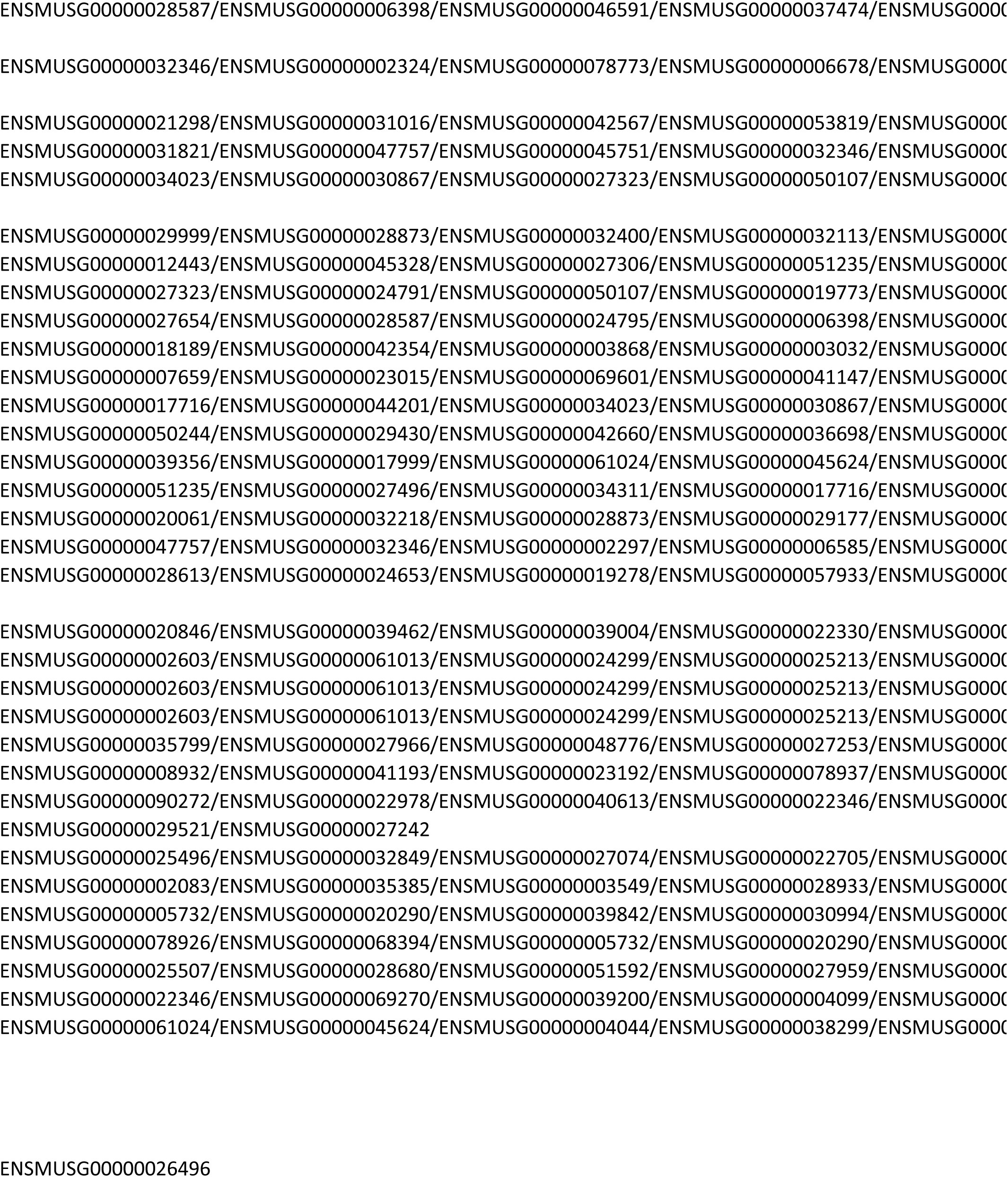
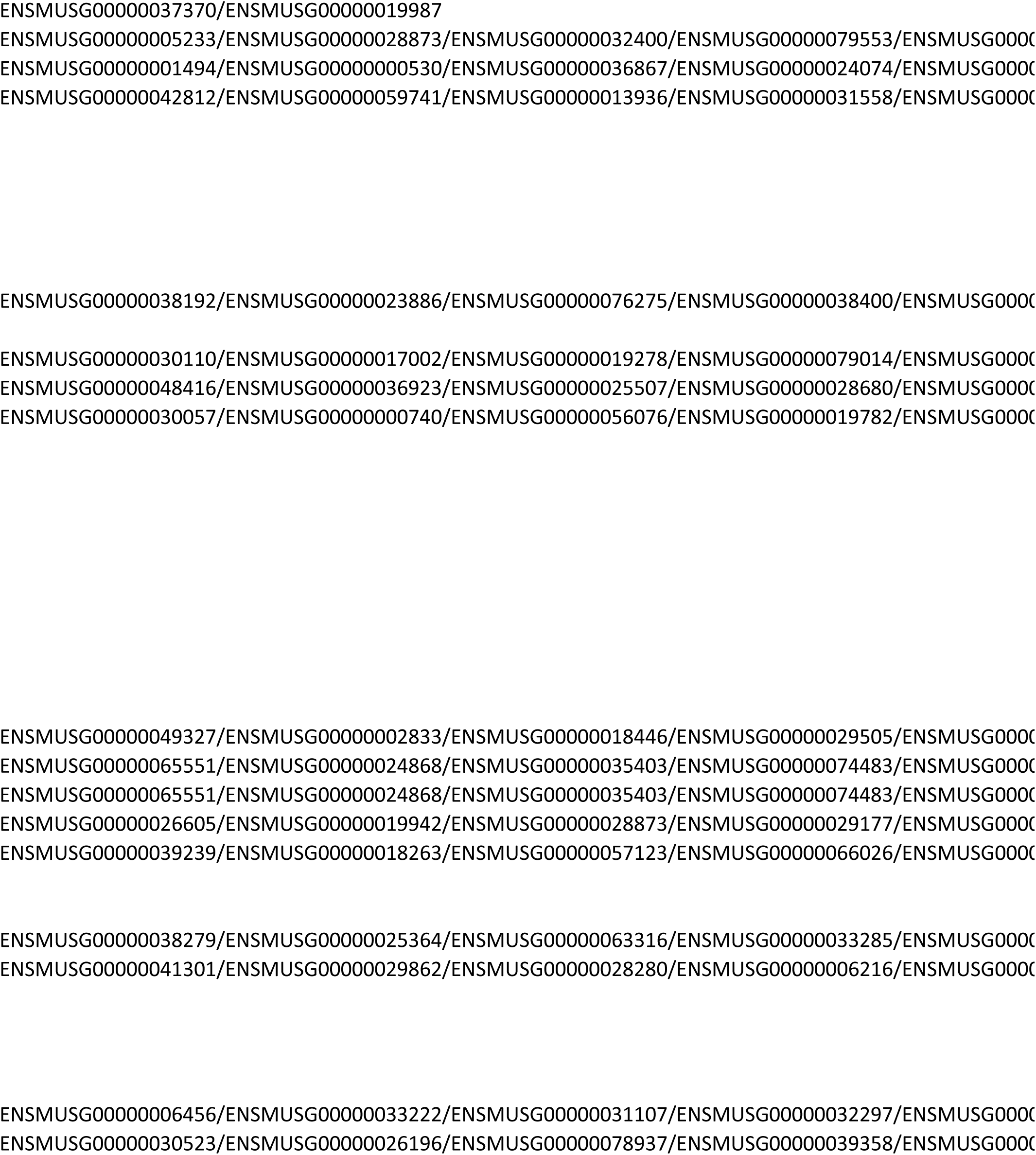
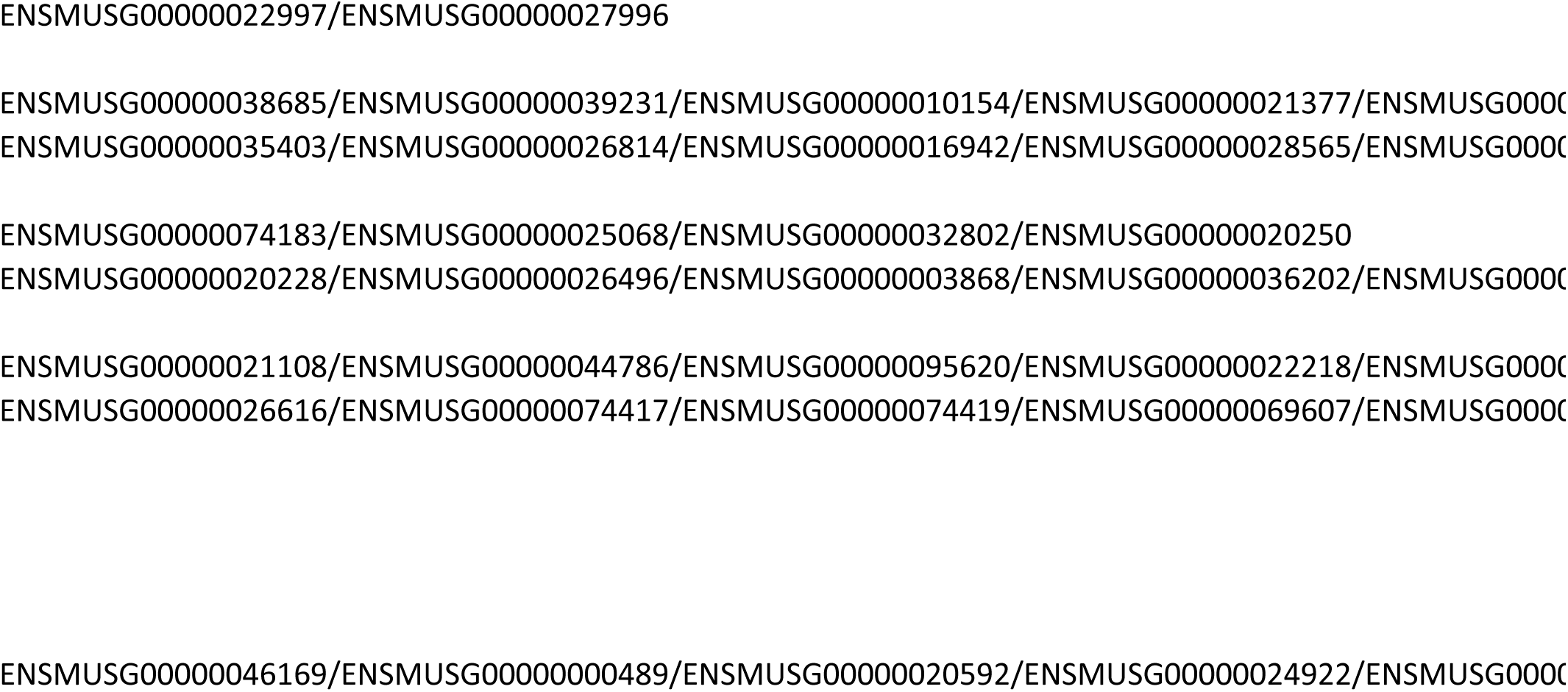
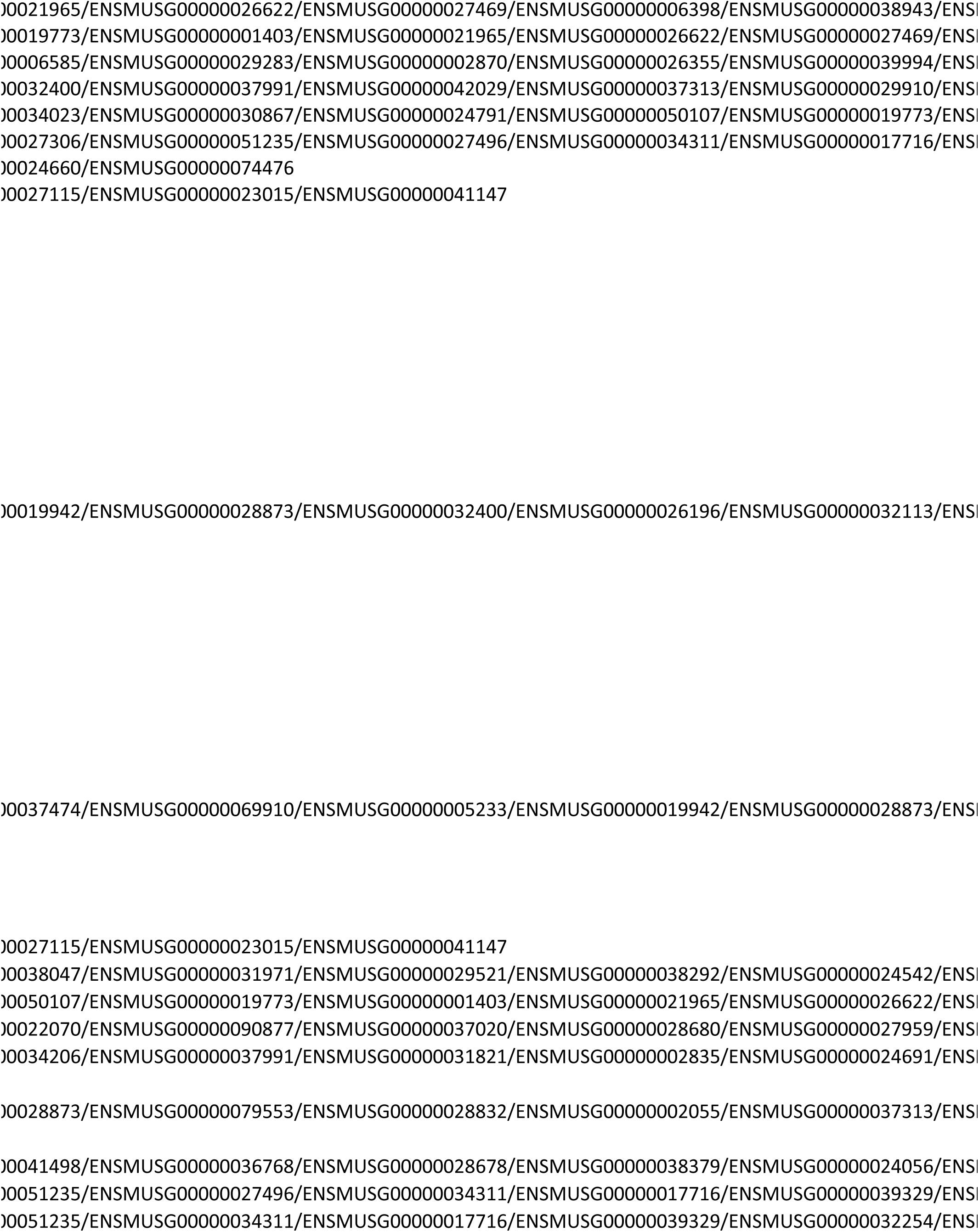
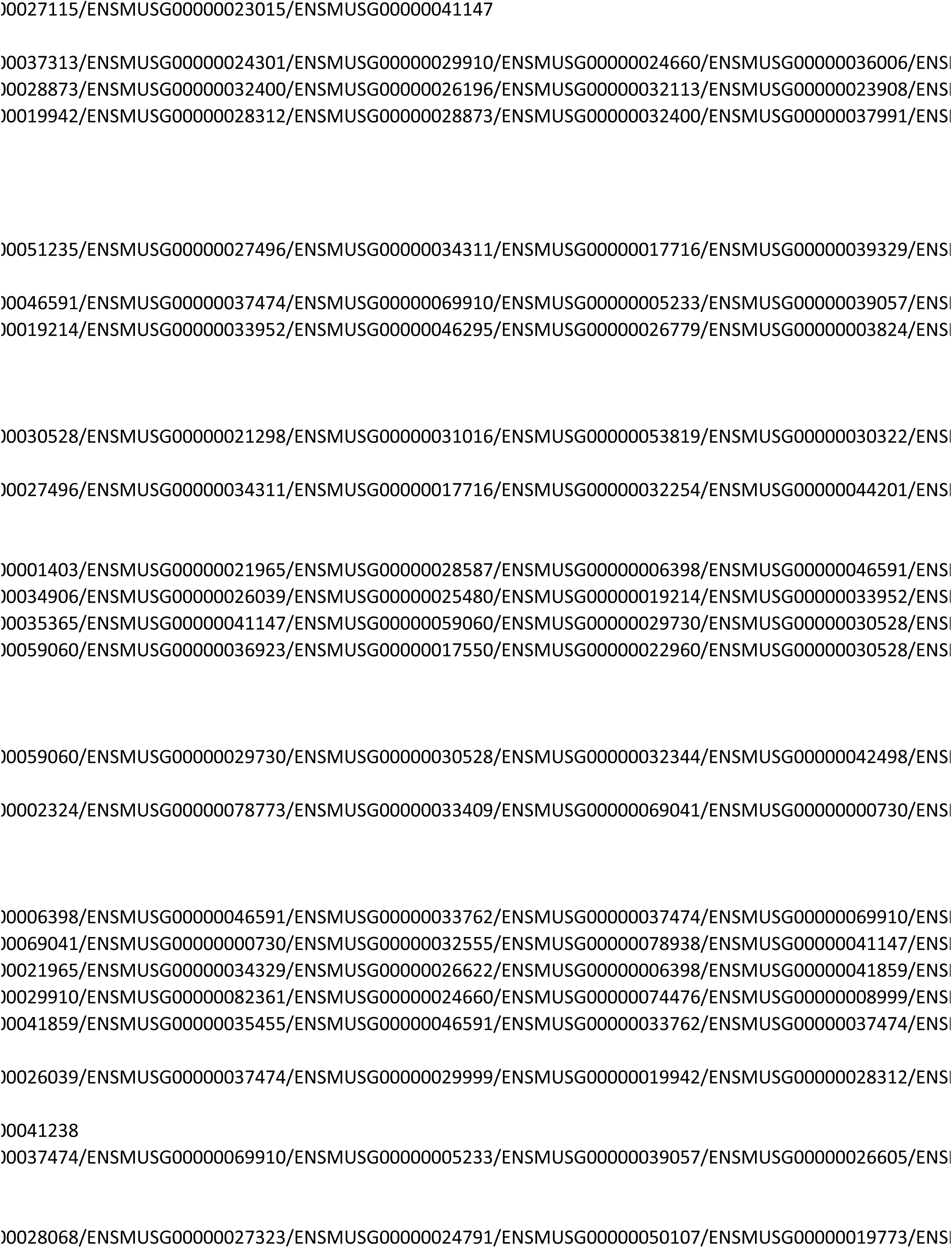
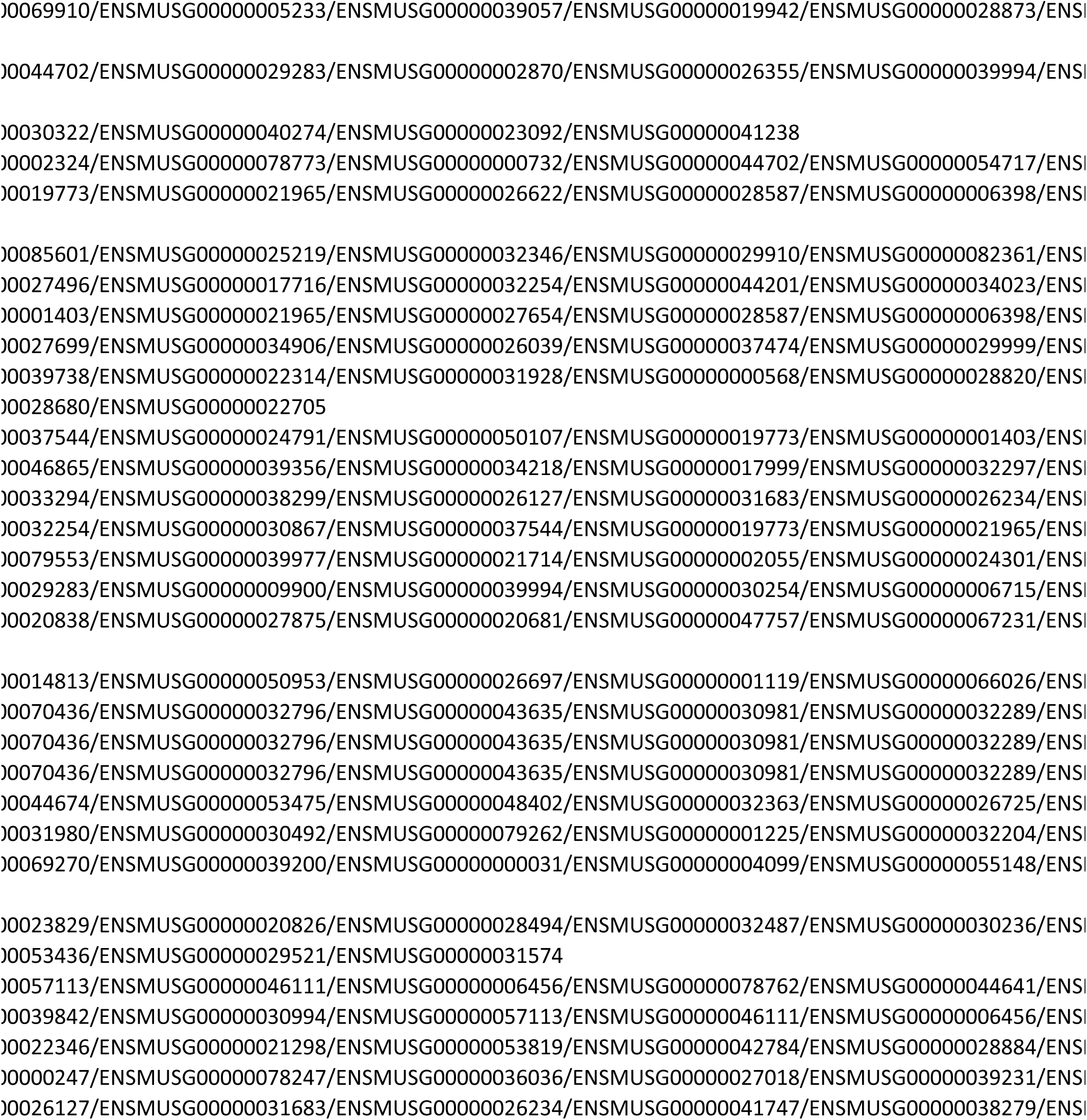
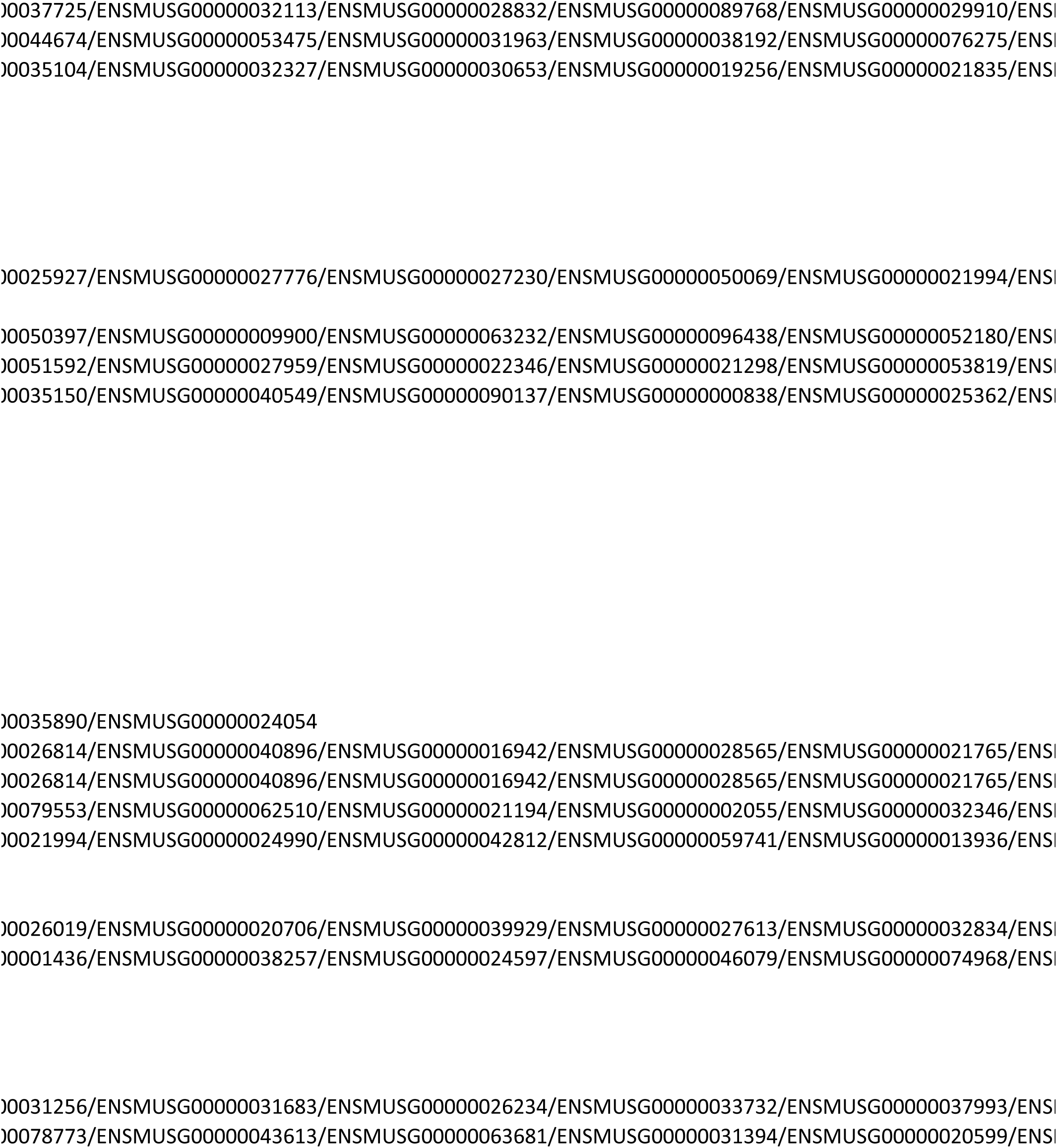
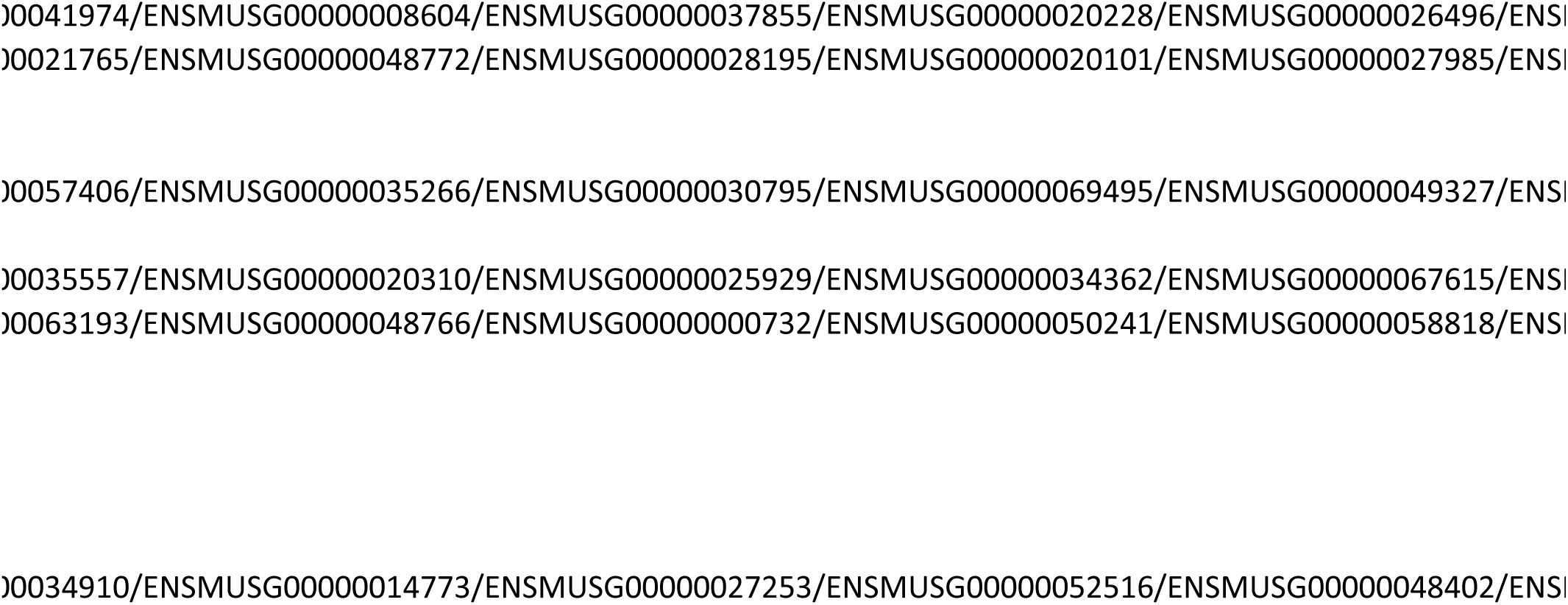
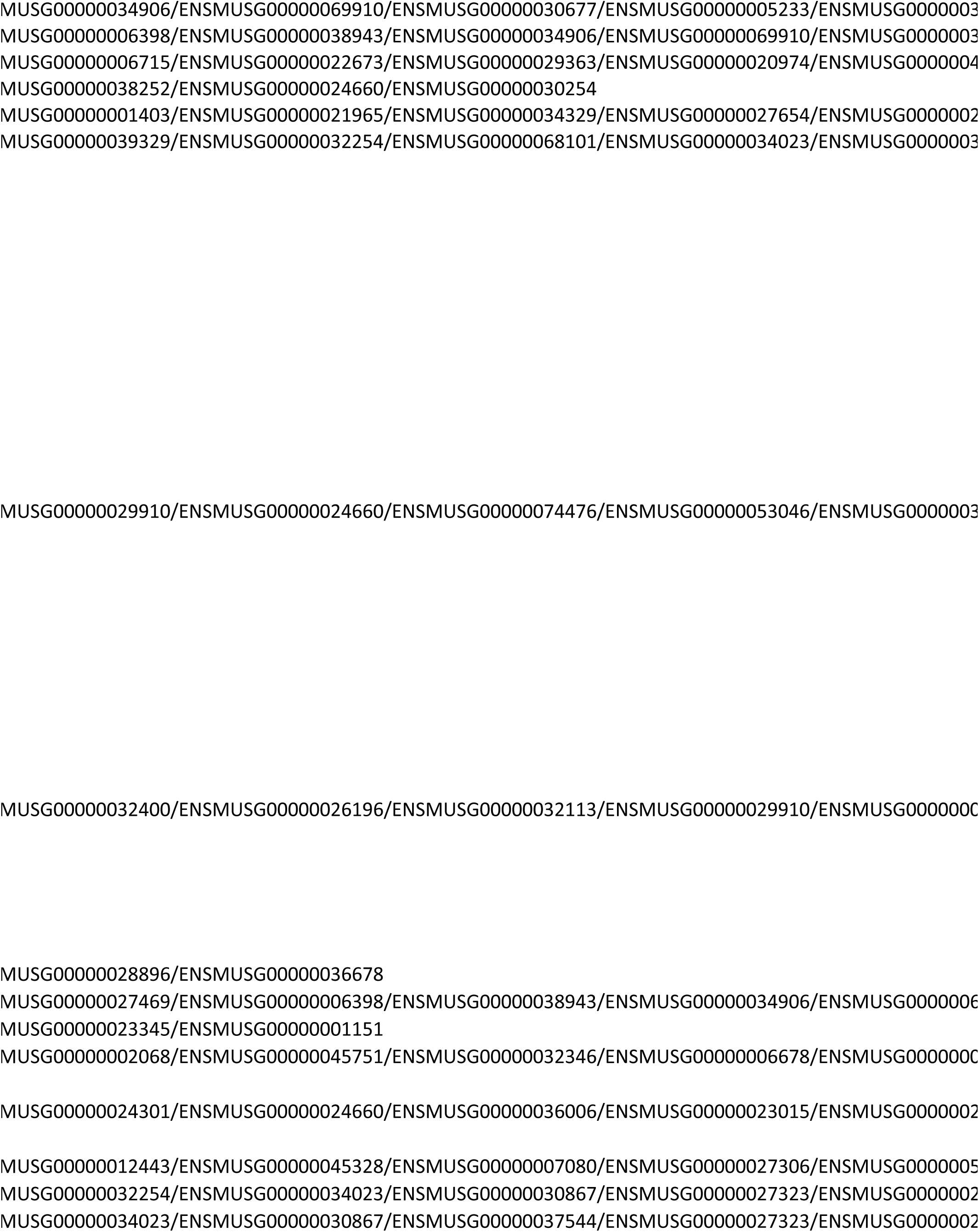
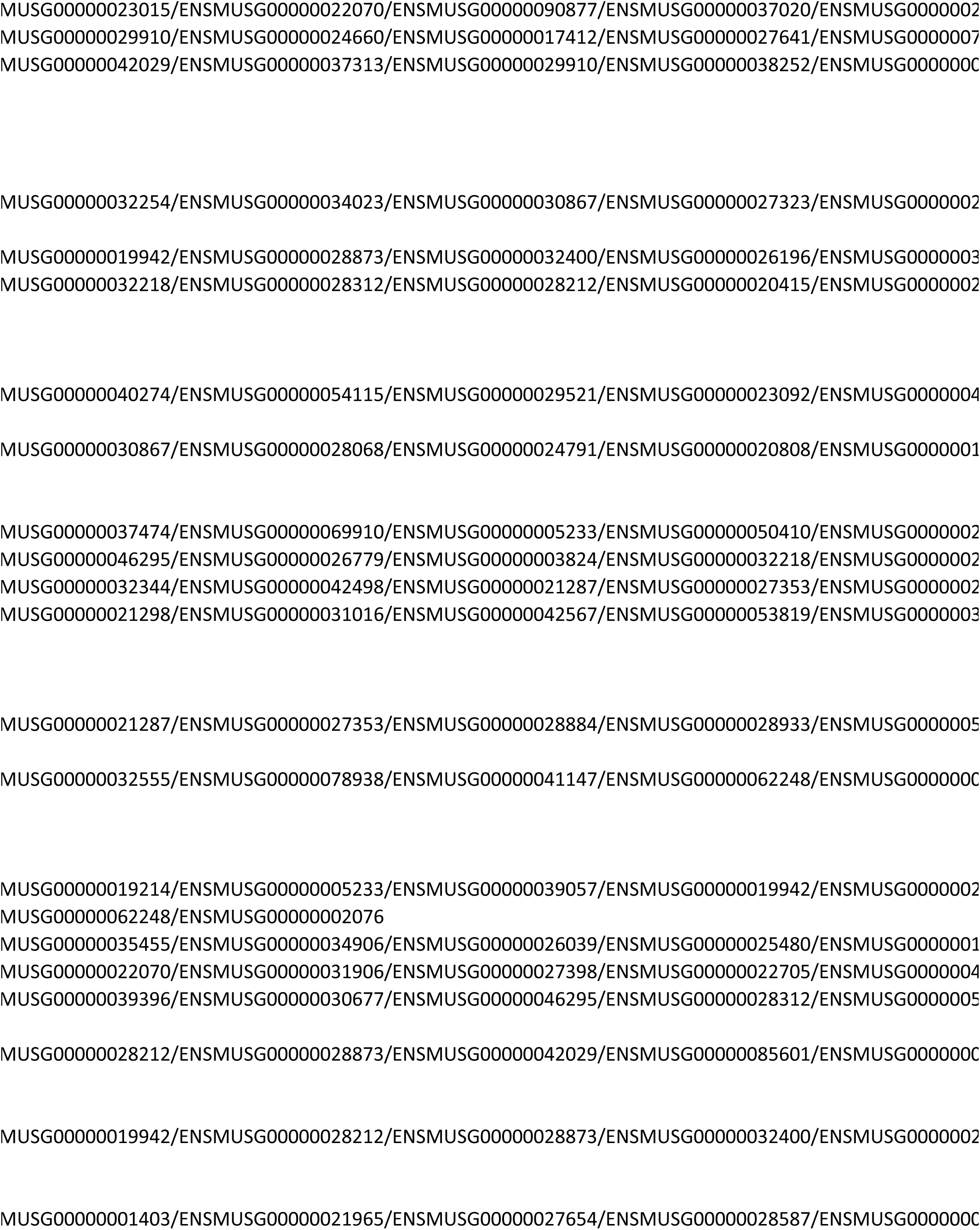
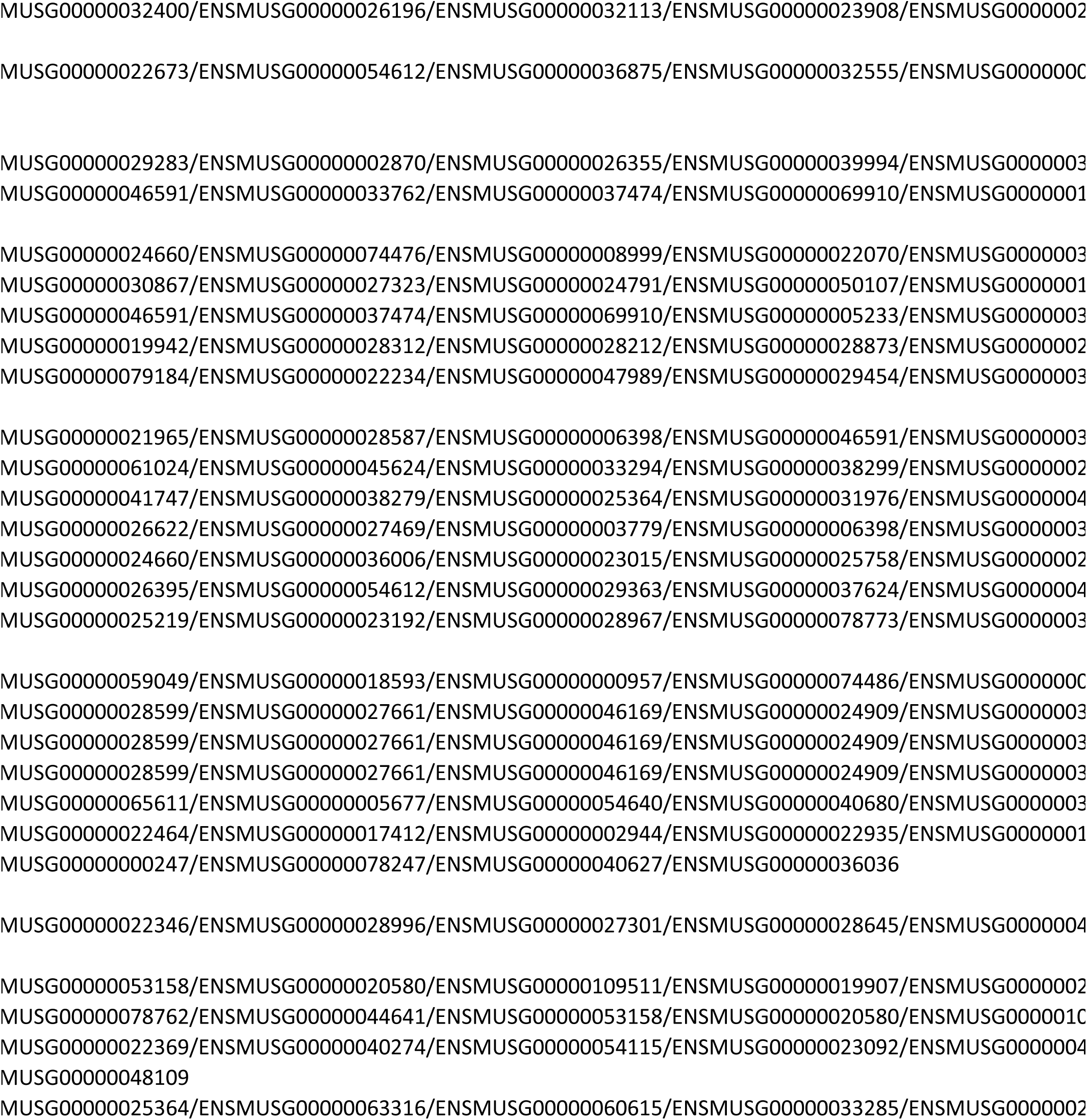
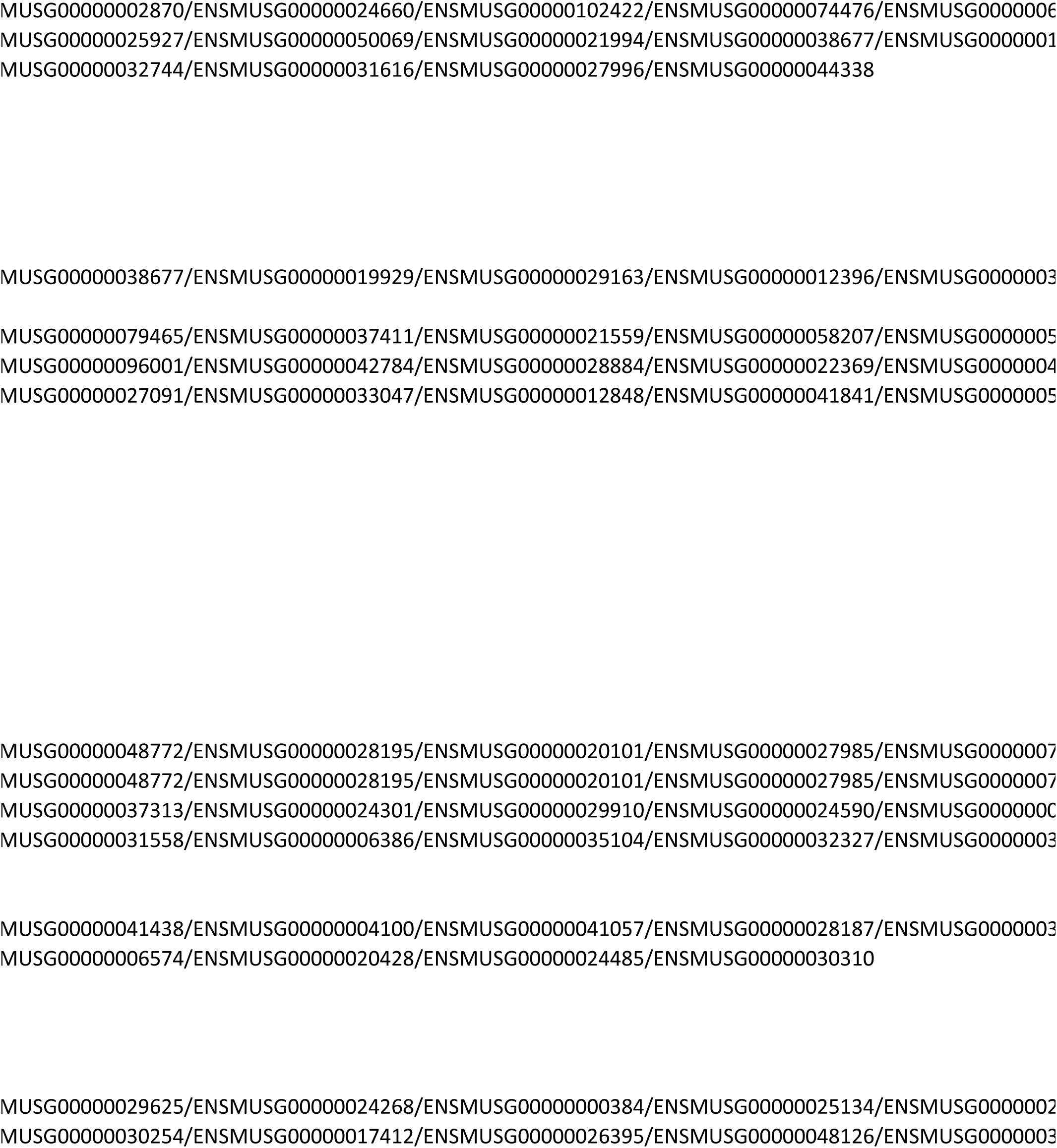
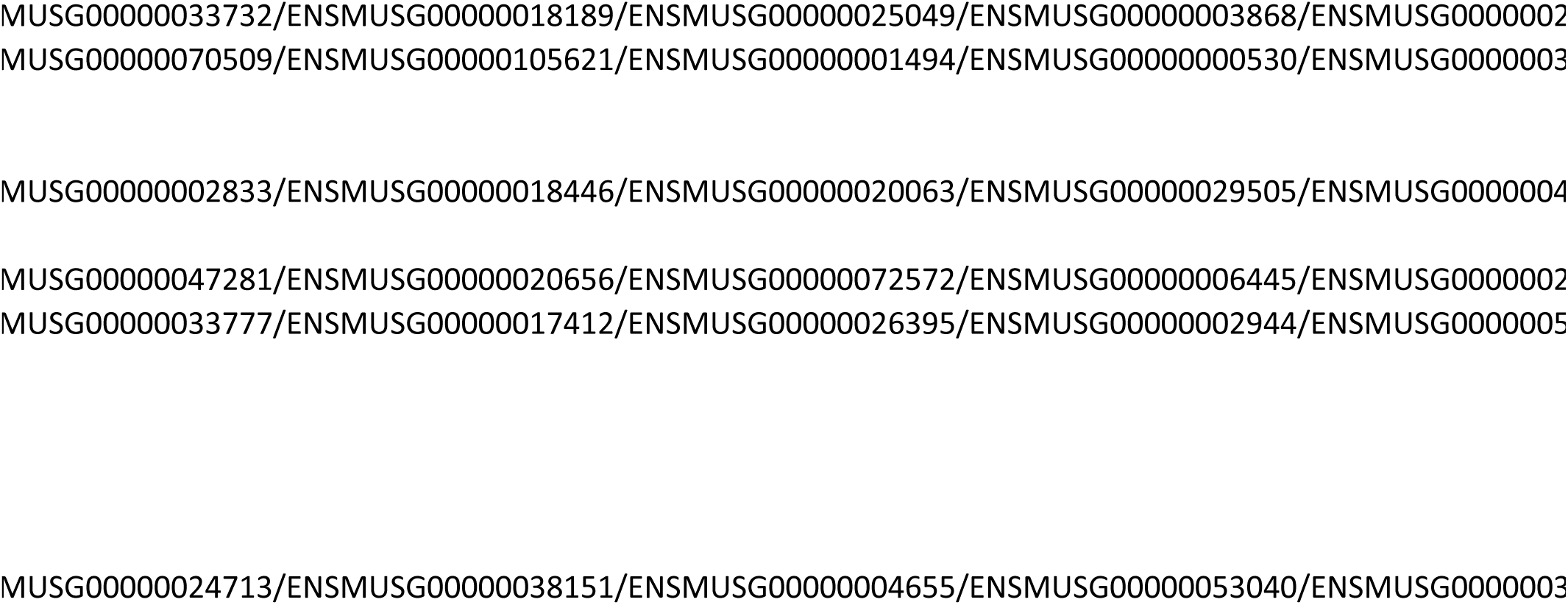
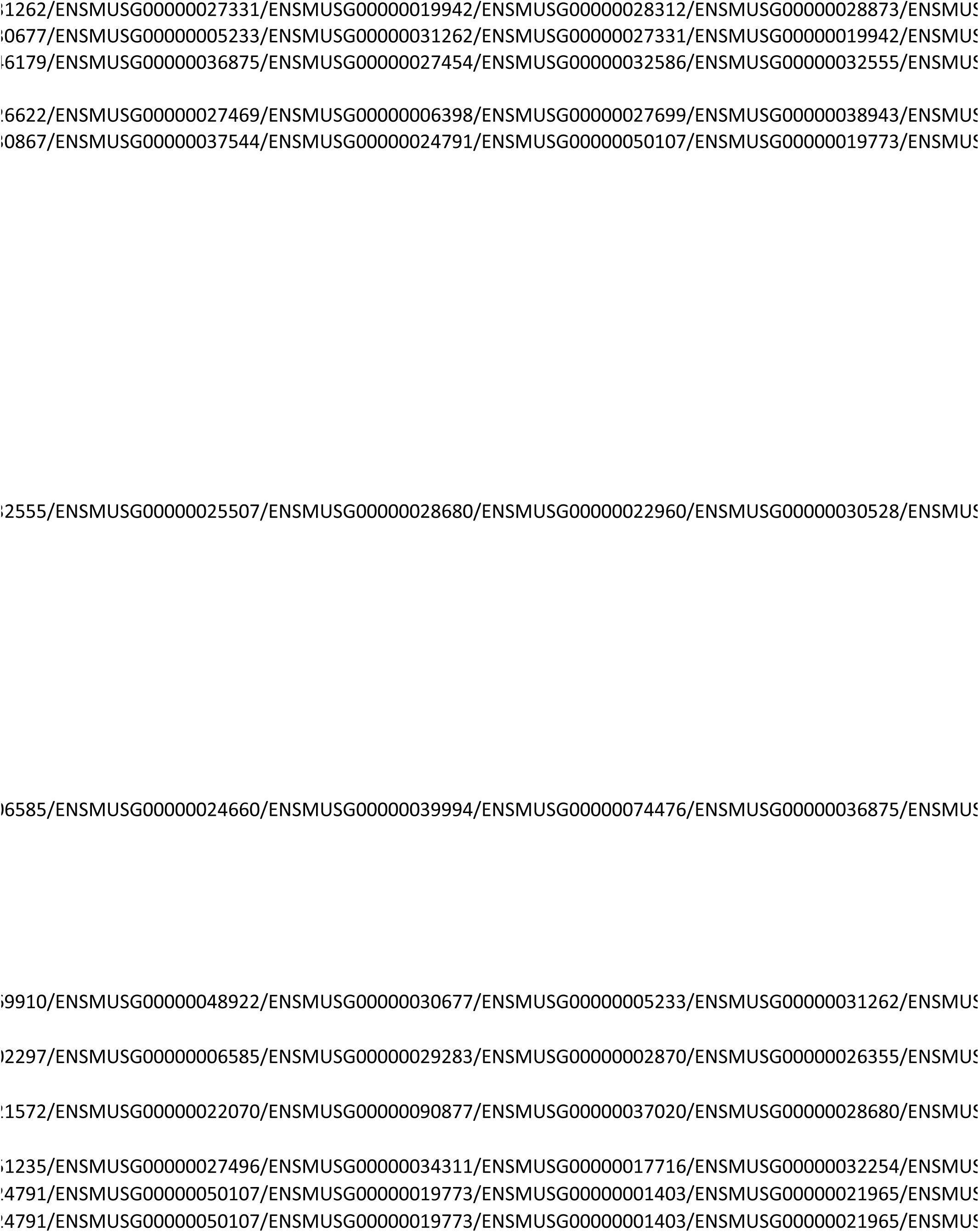
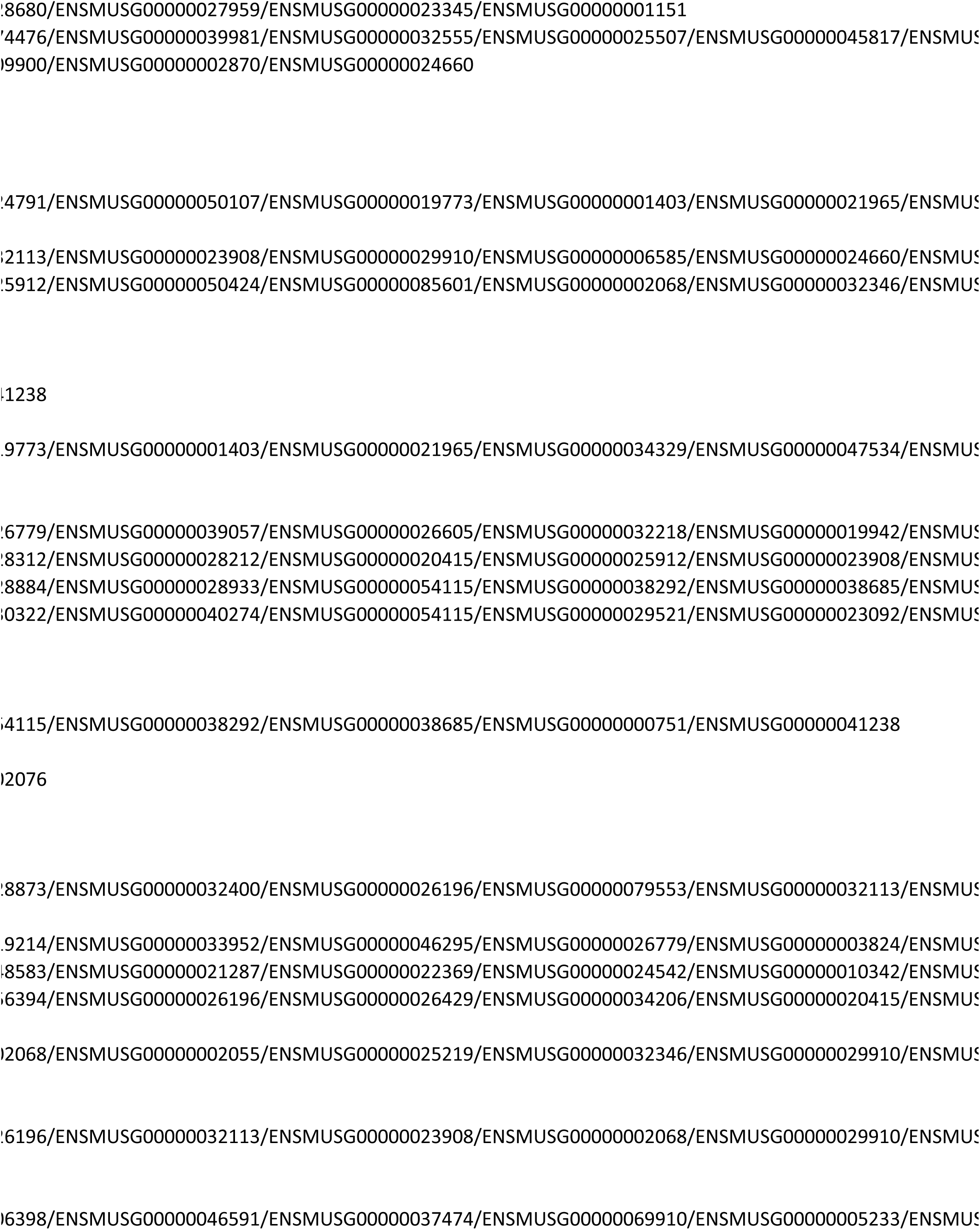
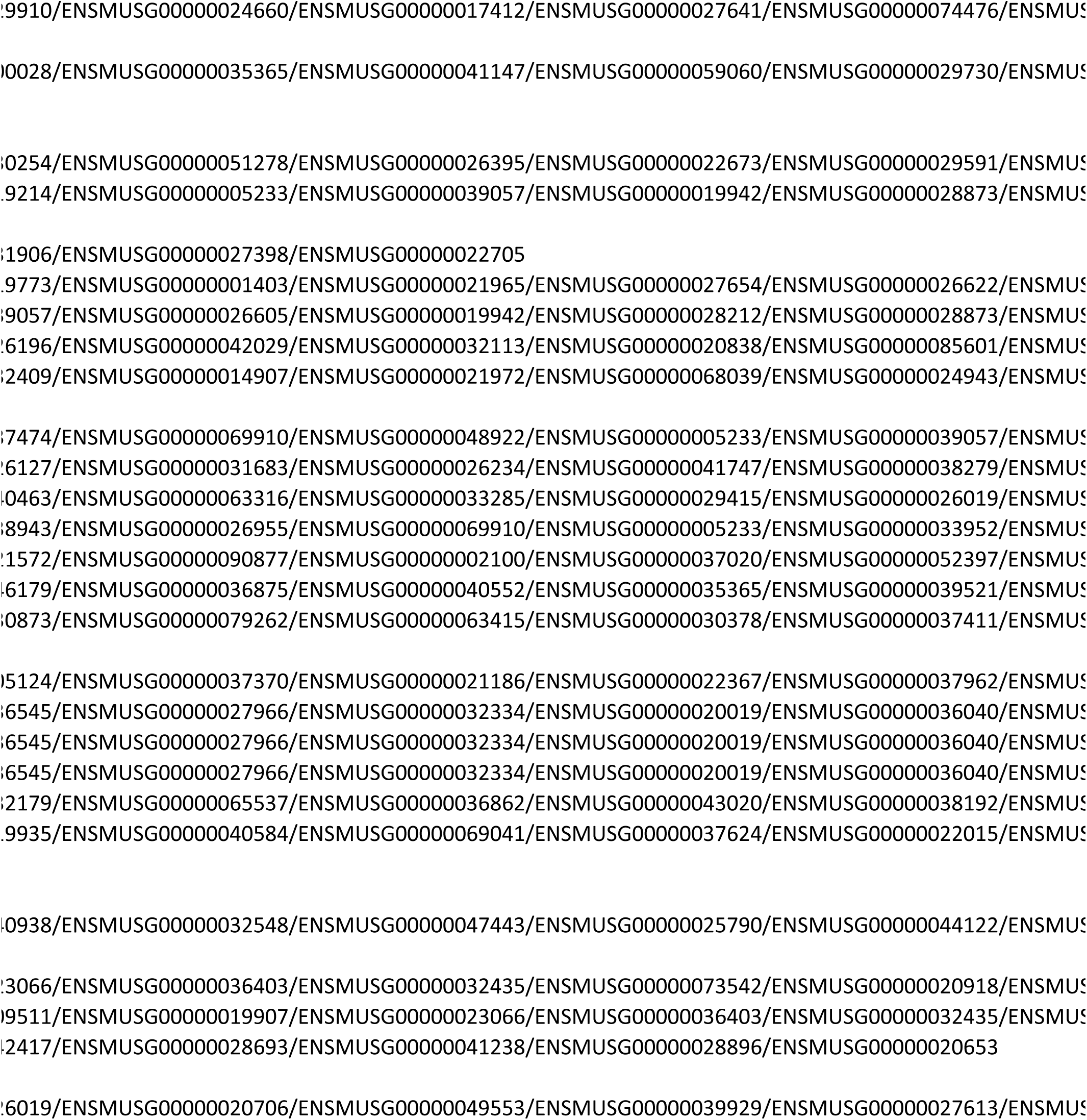
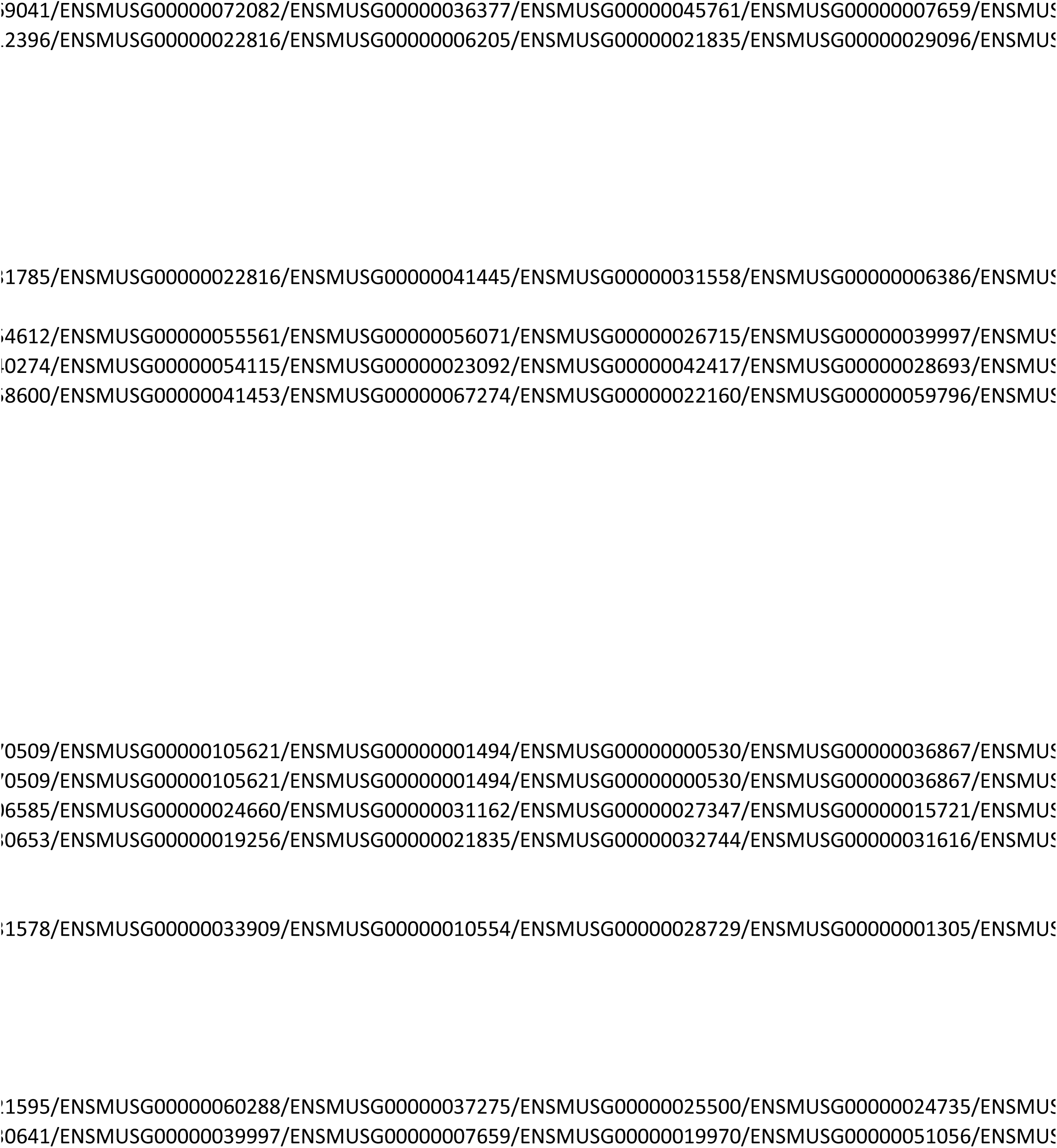
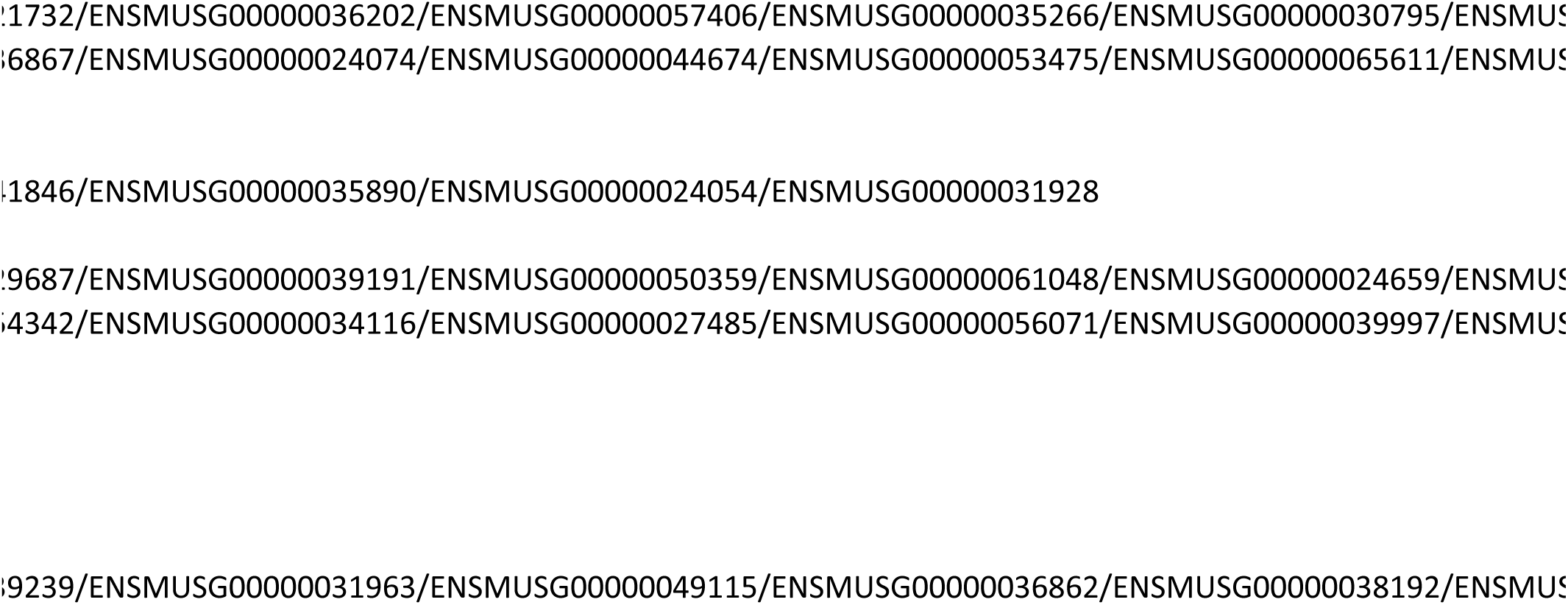
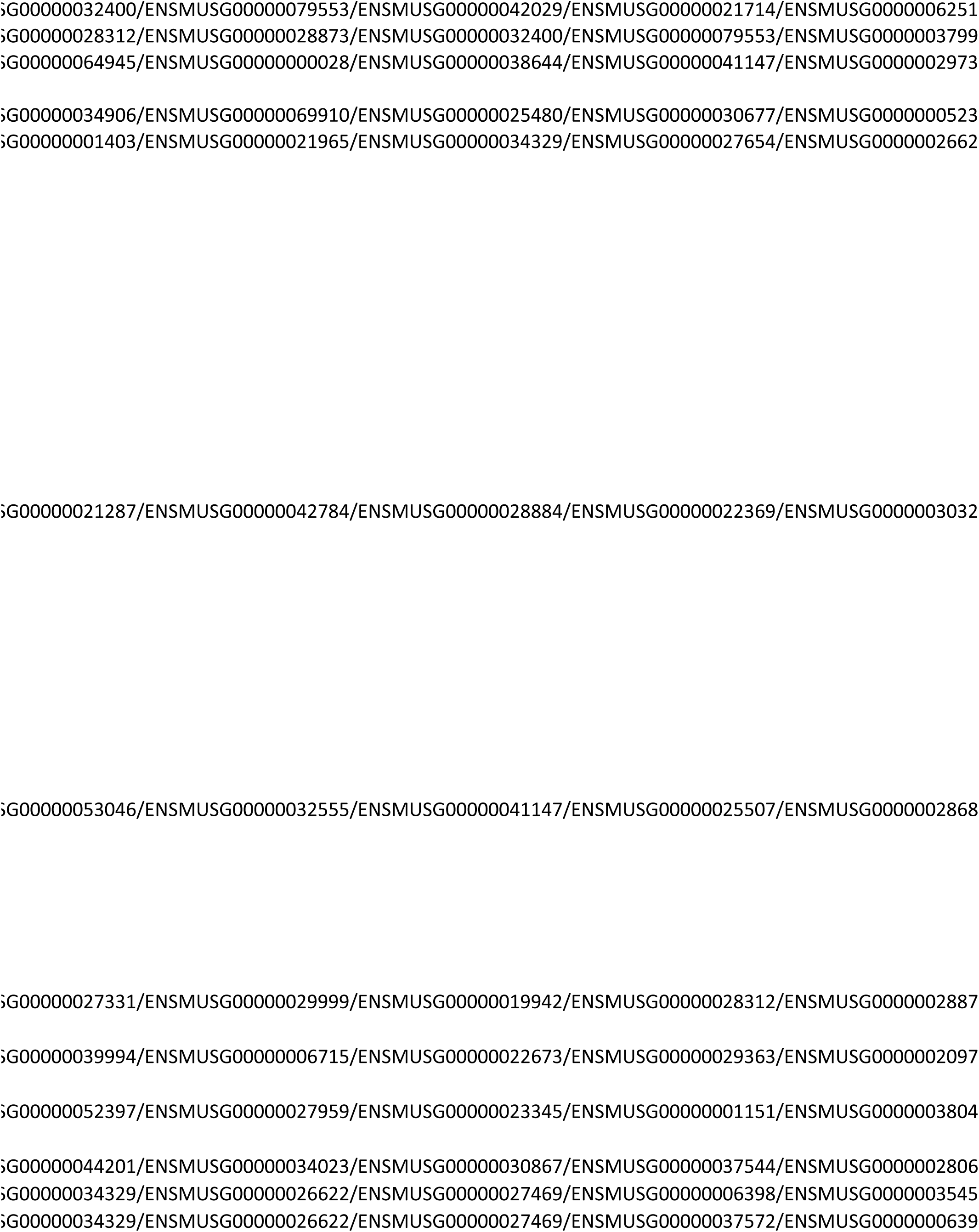
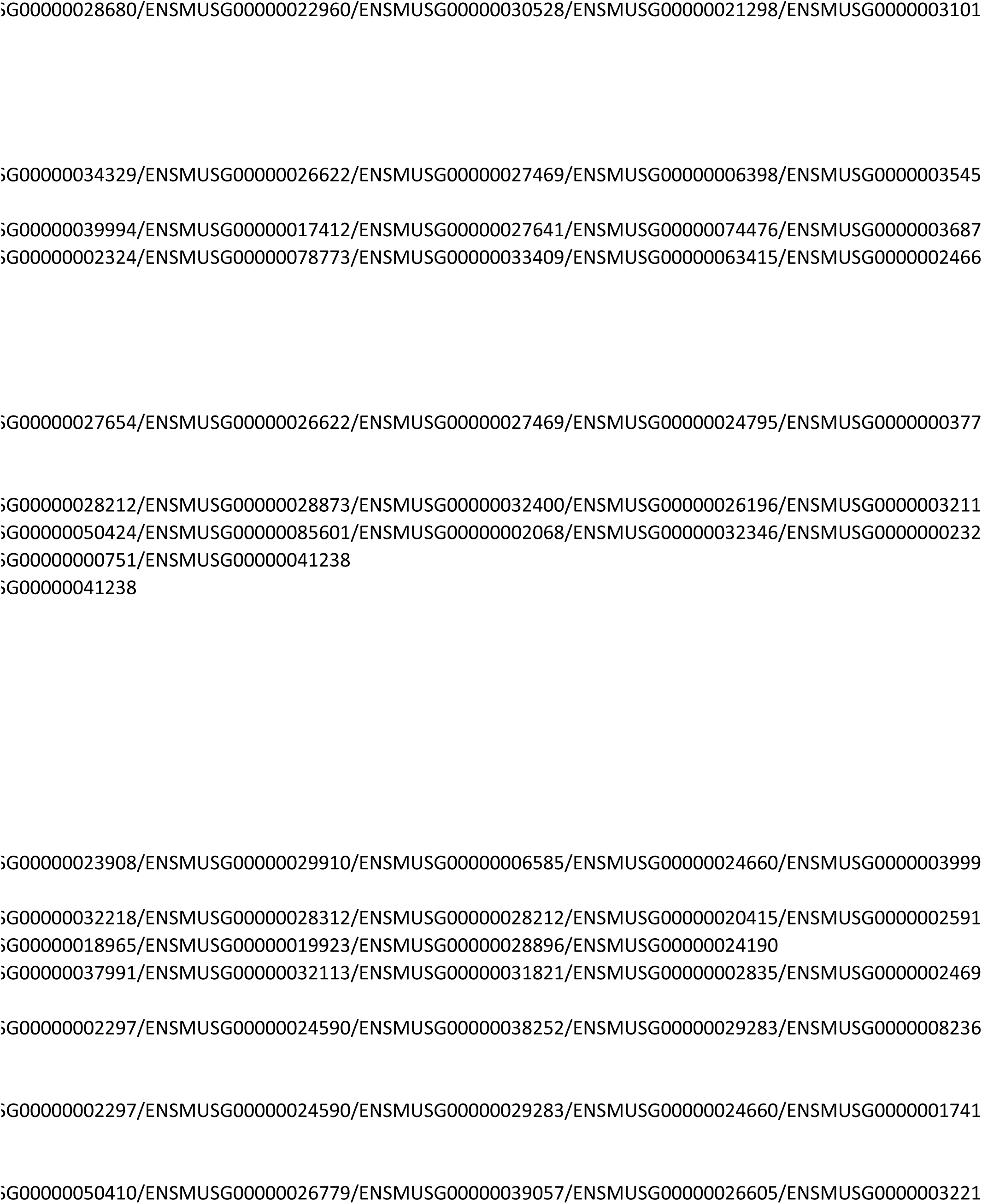
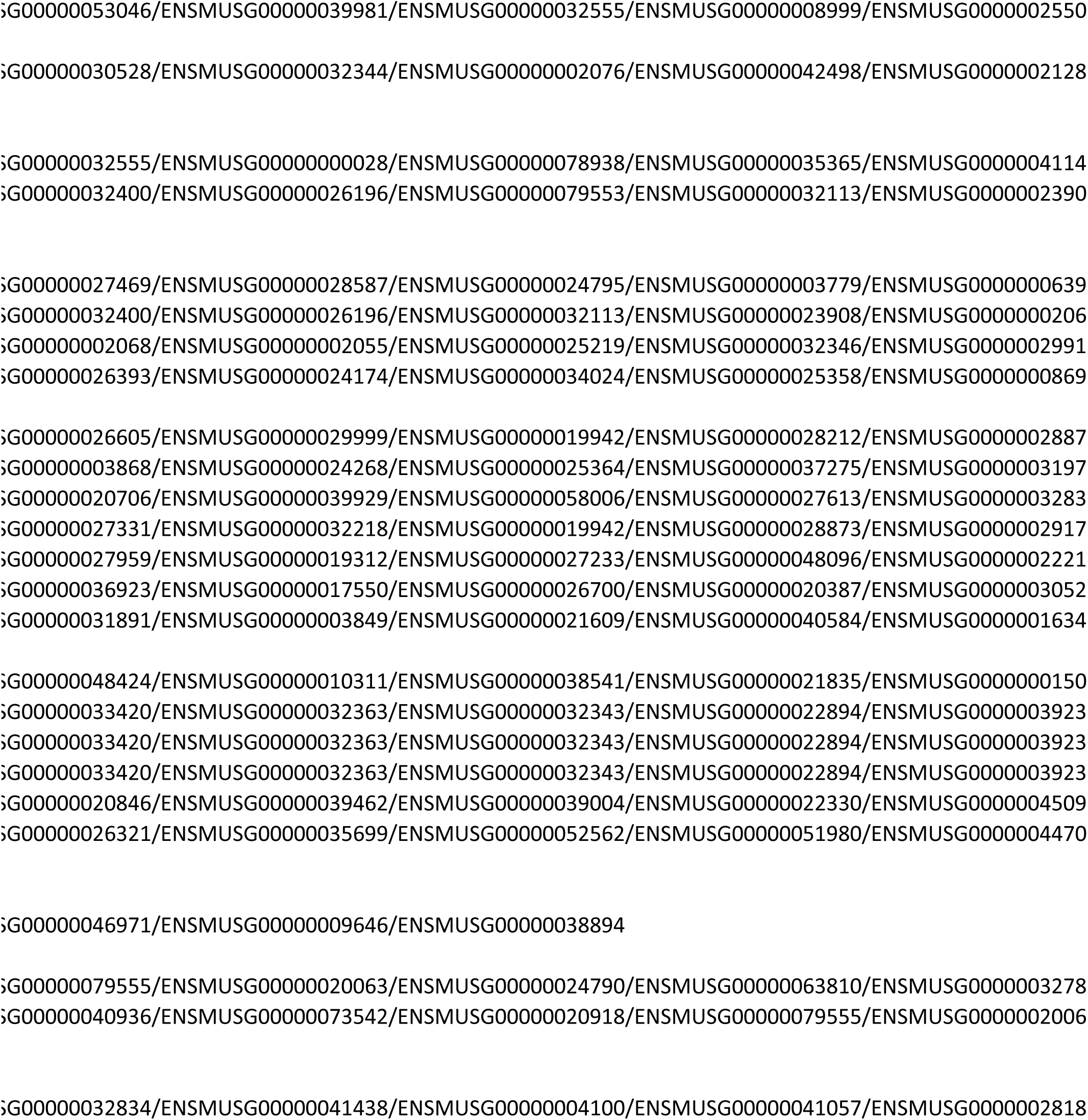
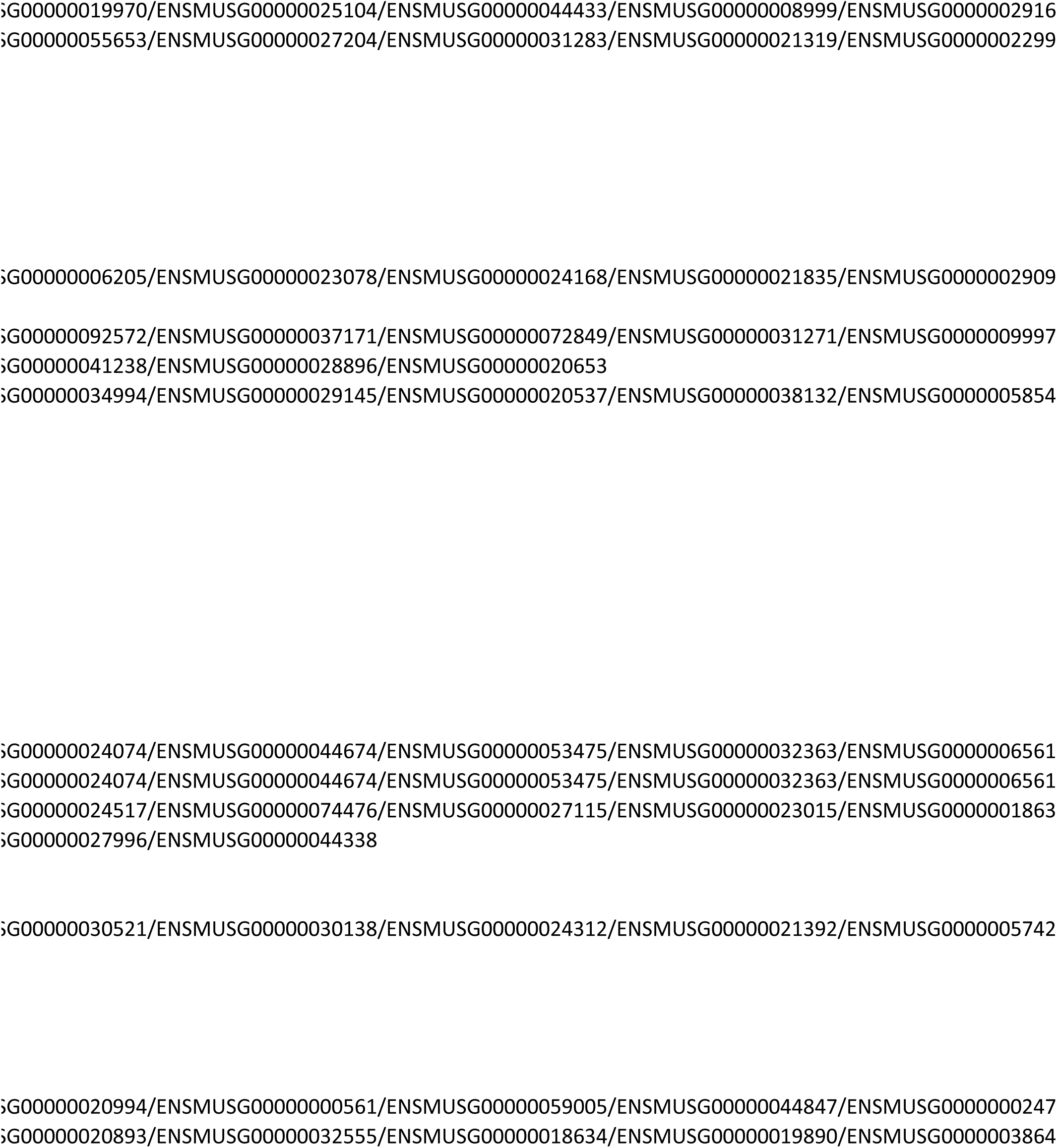
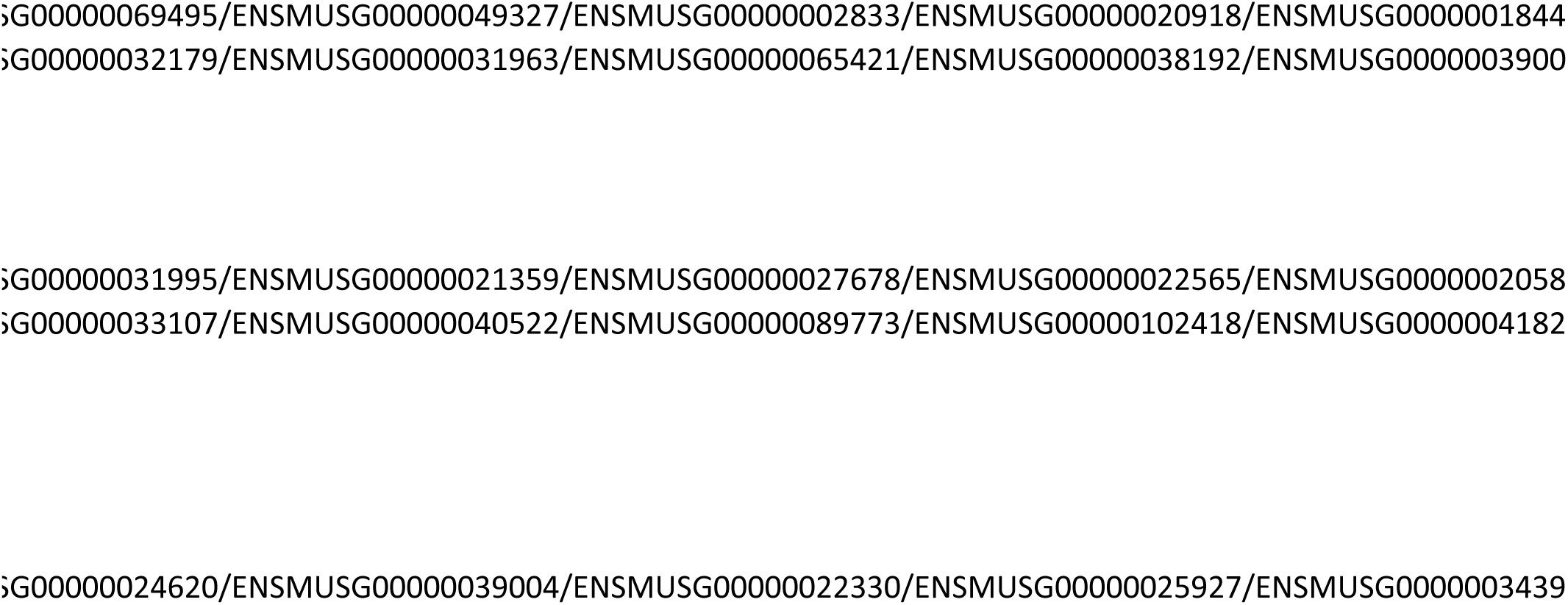
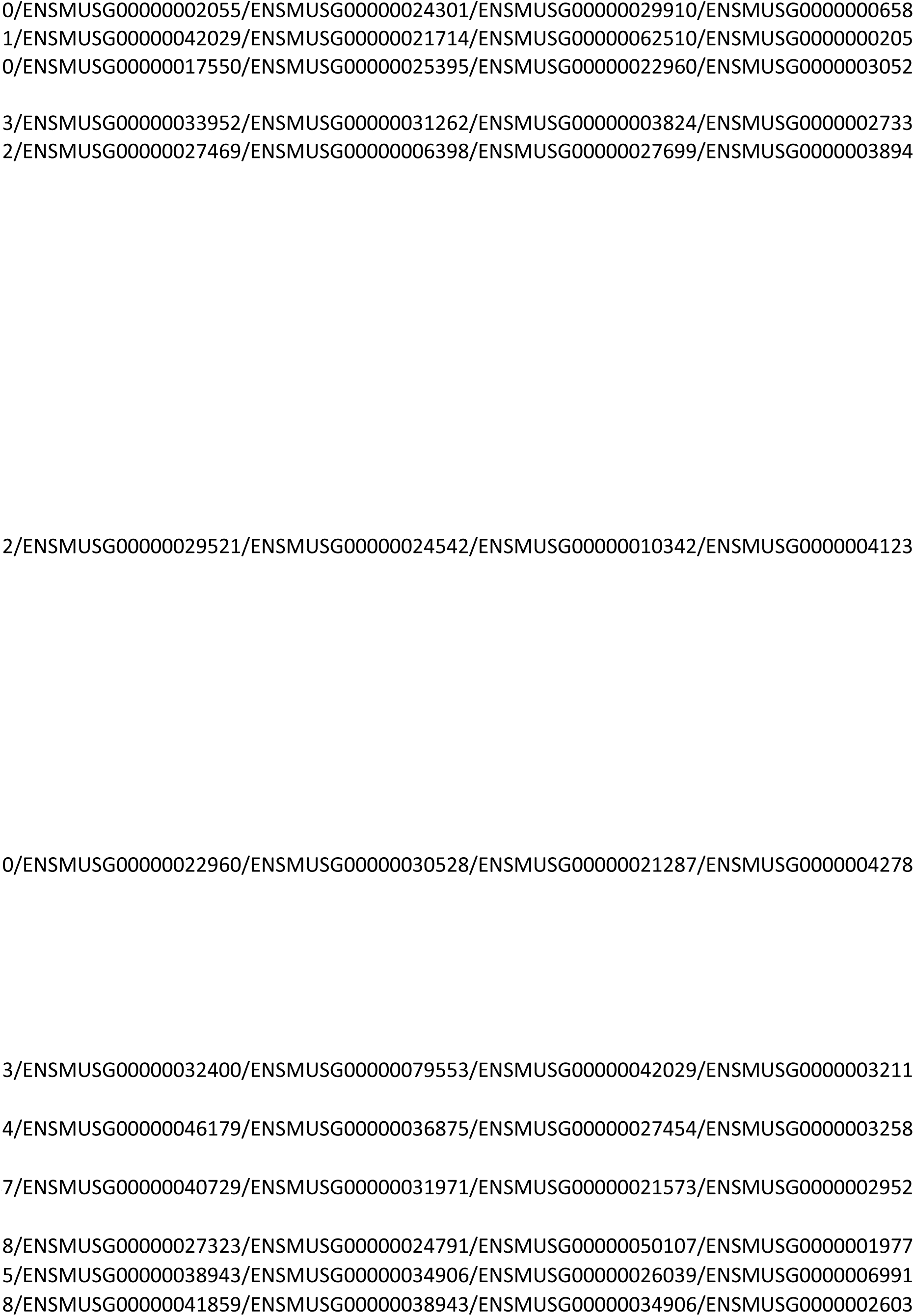
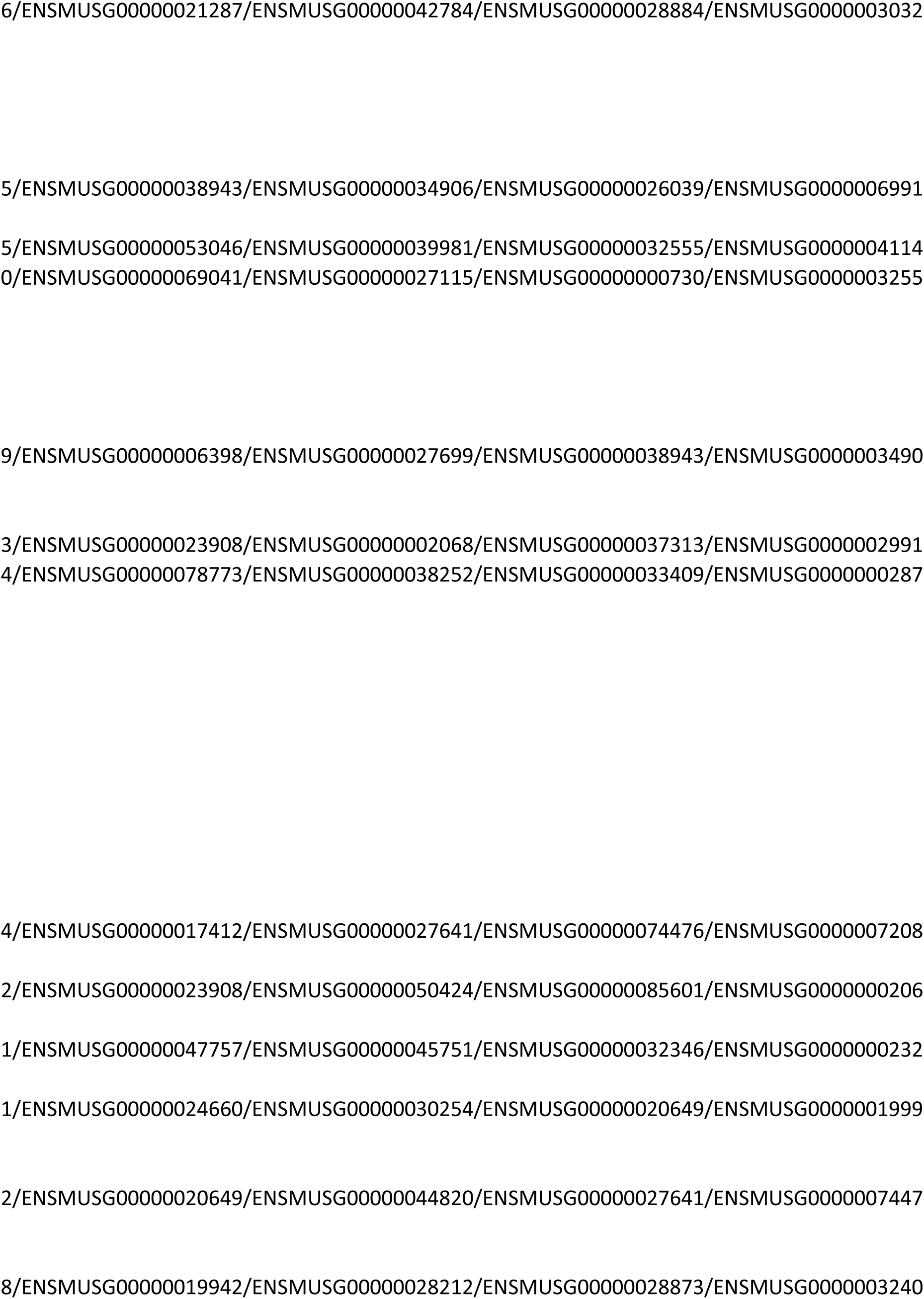
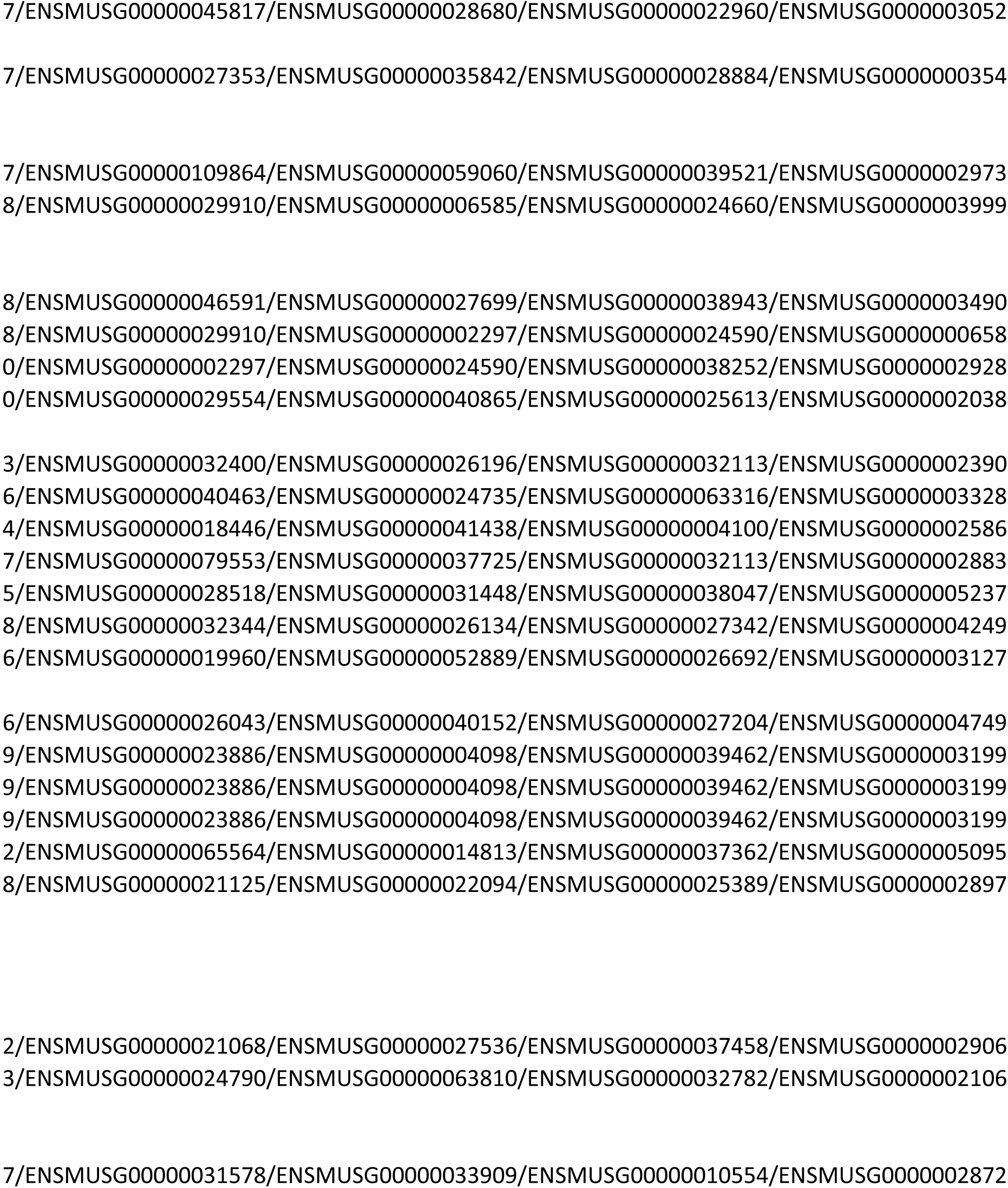
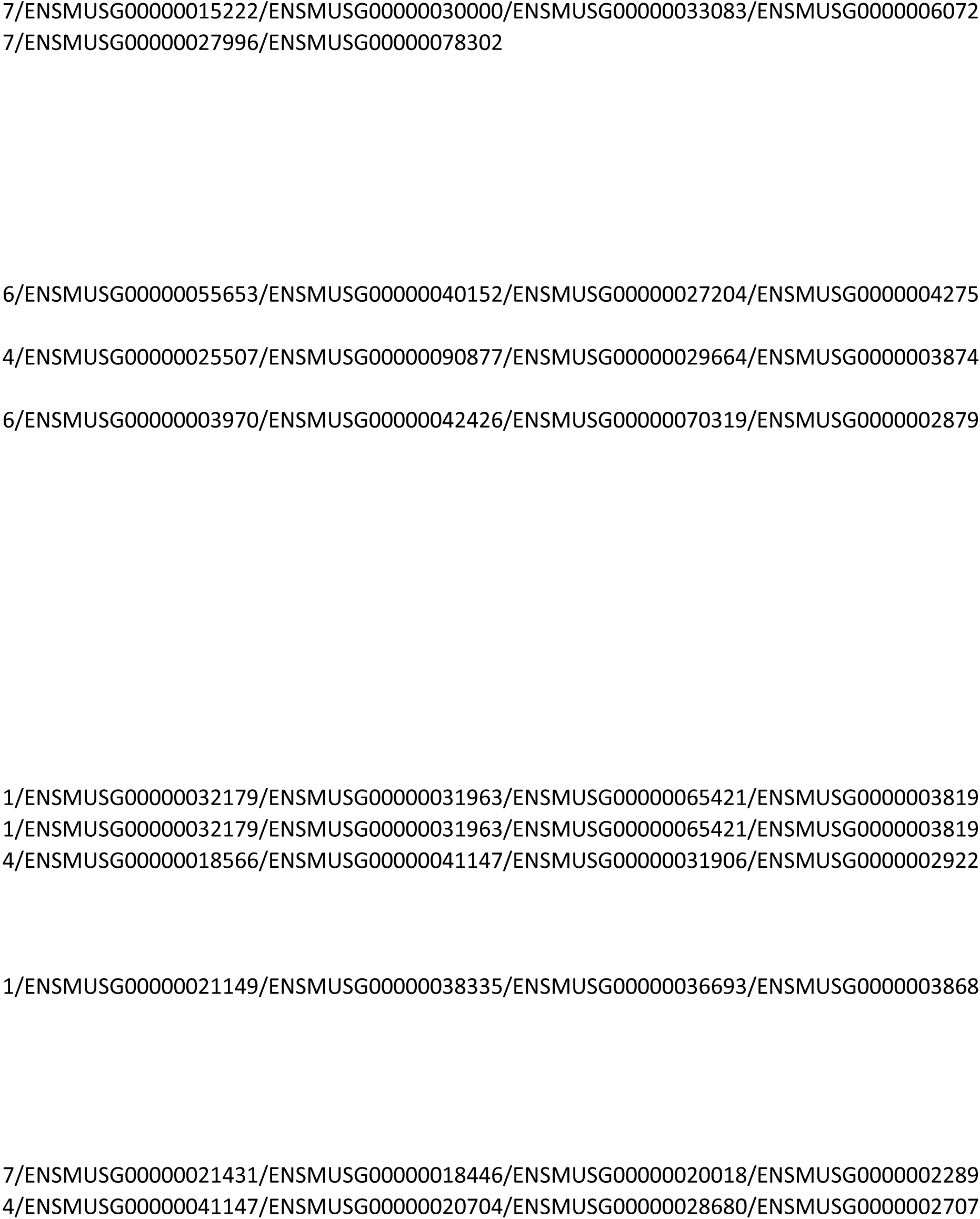
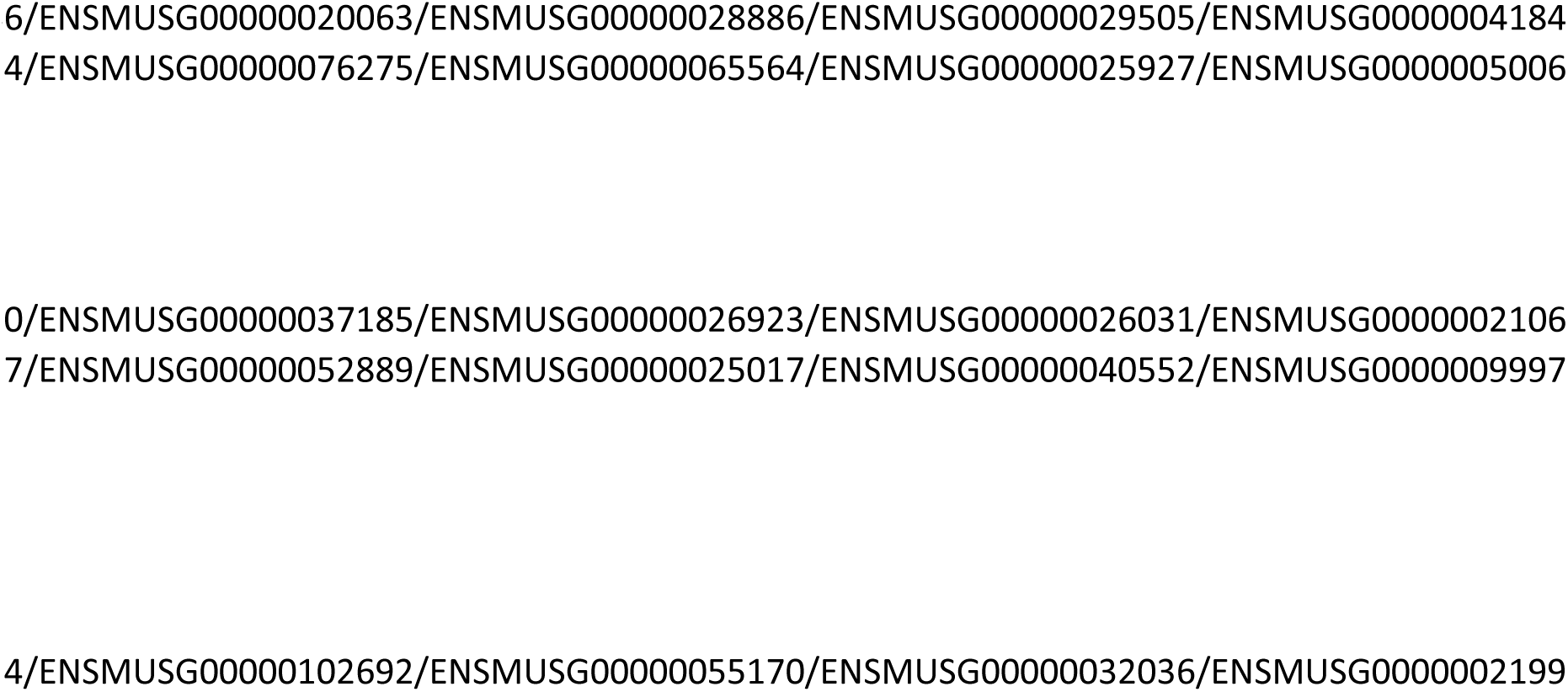

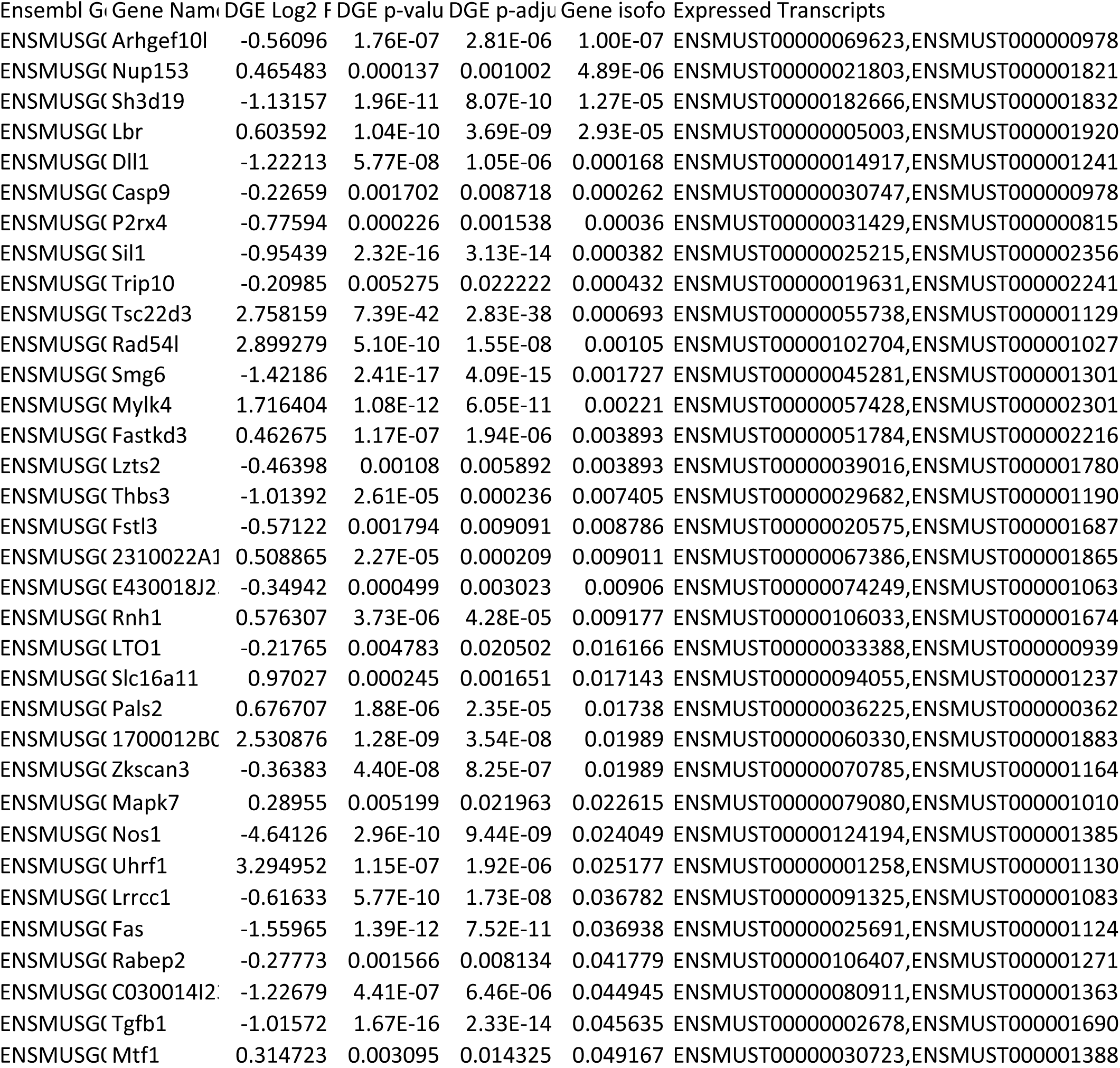
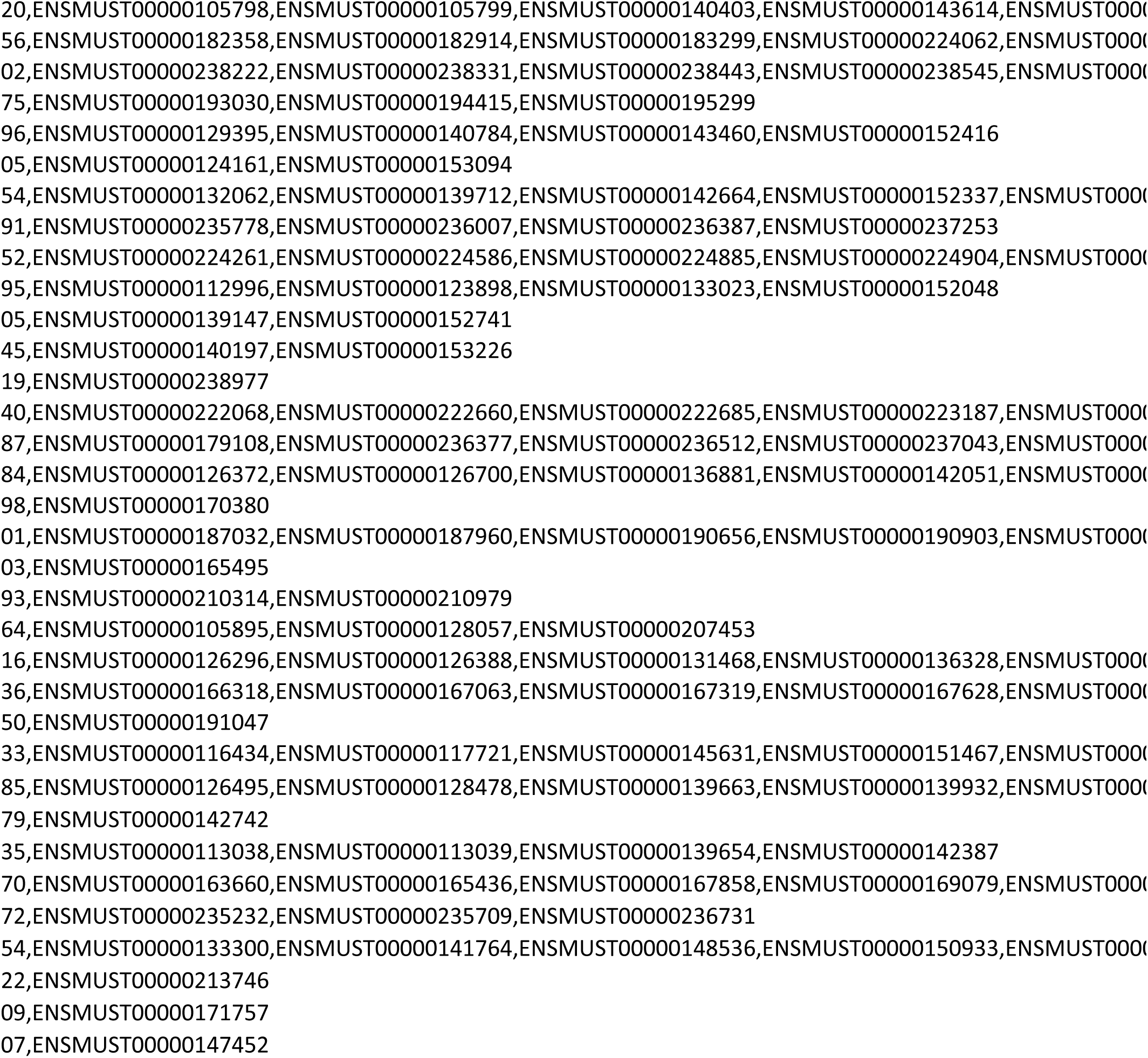
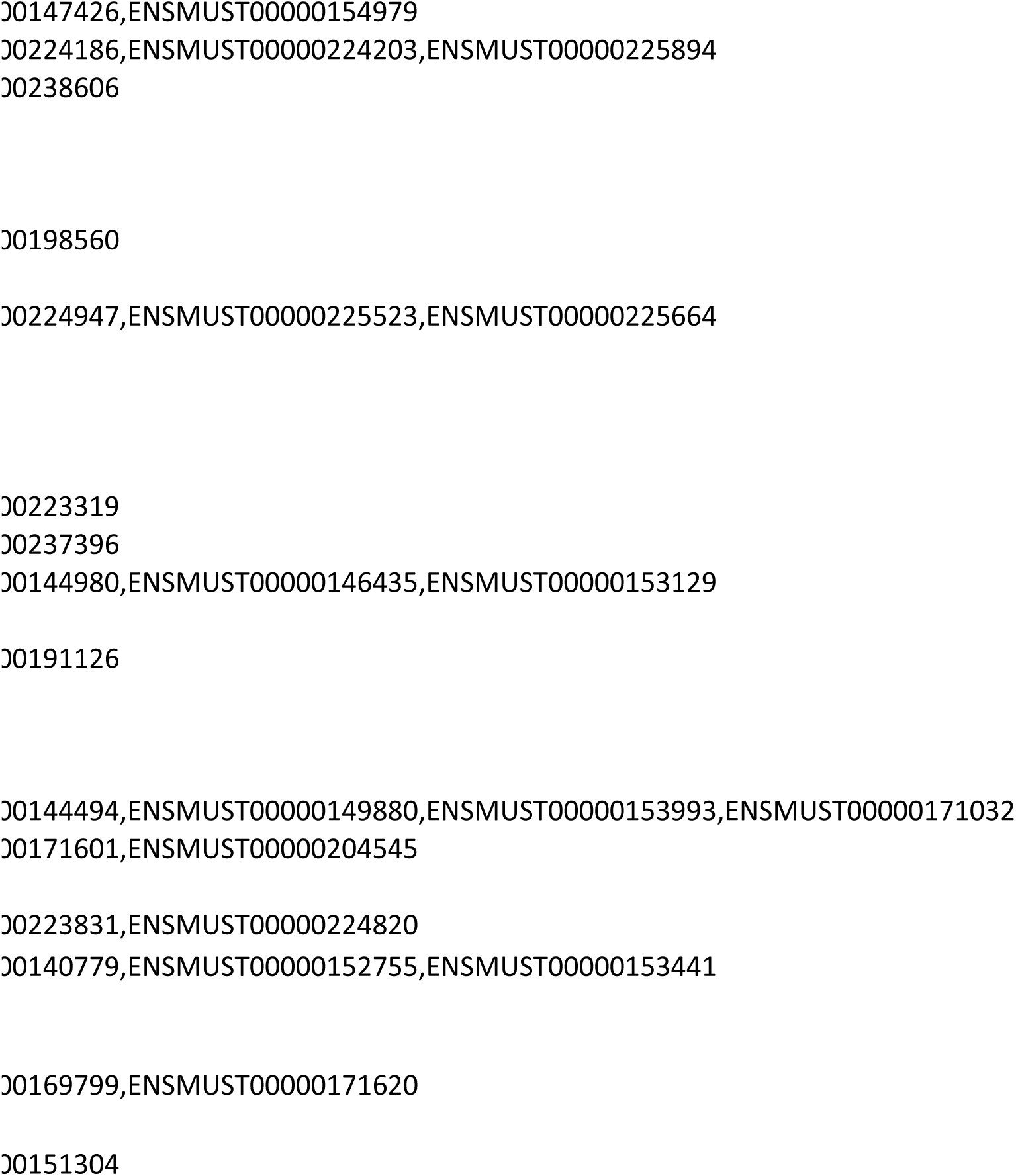

